# Task-optimized models of sensory uncertainty reproduce human confidence judgments

**DOI:** 10.1101/2025.10.31.685933

**Authors:** Lakshmi Narasimhan Govindarajan, Sagarika Alavilli, Josh H. McDermott

## Abstract

Sensory input is often ambiguous, leading to uncertain interpretations of the external world. Estimates of perceptual uncertainty might be useful in guiding behavior, but it remains unclear whether humans explicitly represent uncertainty in naturalistic settings, and whether any such representations are normatively correct. Progress has been hindered by the absence of stimulus-computable models that estimate uncertainty. We developed a class of task-optimized models that generate probability distributions over perceptual estimates. To assess whether human uncertainty representations align with the model’s, we compared human confidence judgments, which might indirectly reflect uncertainty representations, to confidence judgments extracted from the model’s uncertainty. In both sound localization and pitch perception, human confidence varied systematically, being lower for stimuli that produced more variable estimates across trials. Human confidence tracked model confidence across conditions, suggesting that human uncertainty representations accurately reflect the actual uncertainty of perceptual estimation. The modeling framework is extensible to other perceptual domains.

## Introduction

Our senses sample the environment with measurements that are noisy, and that only indirectly reflect the structure of the world around us. Perceptual estimates of this distal structure are thus inevitably uncertain. The extent of this uncertainty depends on the stimulus. For instance, when we view a tree trunk in broad daylight, its distance can be estimated with low uncertainty. But if you were to glimpse the tree through fog, the uncertainty would be much higher, because the sensory evidence would be consistent with a relatively wide range of possible distances.

There is much reason to expect humans and other organisms to monitor the uncertainty of their perception. Our actions are dependent on our perception of the world, and acting on uncertain estimates of the world could be costly. On these grounds one might expect an organism to have explicit internal representations of the uncertainty of its perception. For instance, in addition to or instead of deriving a single best estimate of the variables that they infer, sensory systems could compute a probability distribution over the variable^1–3^. If this distribution were to accurately reflect the actual probability of different values of the variable given a stimulus, it could enable quantitative estimation of uncertainty, for instance via the spread of the distribution. On the other hand, accurately deriving such a distribution might not always be tractable.

There is considerable evidence that estimation uncertainty is implicitly reflected in perceptual judgments. For instance, cue combination is driven by the uncertainty associated with different cues^4,5^, and observers are biased by prior information to an extent that depends on the fidelity of sensory information^6^. Representations of uncertainty might also be explicitly reflected in the feeling of confidence often associated with decisions. Confidence is typically viewed as expressing an internal estimate of the probability that a decision is correct^7^. An extensive literature has documented that humans can reliably report confidence in experimental settings^8–11^. What is less clear is whether these confidence judgments are derived from representations of the uncertainty of perceptual estimation, and whether any such representations accurately reflect uncertainty.

In restricted psychophysical settings, as when discriminating the orientation of a grating, confidence judgments align both with the consistency of a decision^12–16^ and with model estimates of decision uncertainty^14–16^, consistent with the idea that observers accurately represent uncertainty and convert those representations into confidence judgments. However, the models typically used to establish this uncertainty cannot be applied to actual sensory stimuli, and so their predictions have been confined to simple settings, leaving it unclear whether this alignment extends to more complicated settings. It remains possible that in more realistic settings observers are forced to rely on heuristics rather than deriving confidence from representations of uncertainty, or that an observer’s uncertainty representations do not faithfully represent the true uncertainty of their perception.

In a parallel literature, deep learning has revolutionized models of sensory systems^17^. Neural networks optimized for real-world tasks often produce good alignment with human behavior in domains such as object recognition^18^, word, voice, and environmental sound recognition^19–21^, attentional selection^22^, sound localization^20,23^, and pitch perception^24^. However, such models have thus far been poorly suited for studying uncertainty. In particular, the cross-entropy objective function standardly used to train large-scale models of sensory perception incentivizes models to be overconfident^25–27^. As a result, although current neural network models provide normative insights into patterns of performance^20,22–24^, as typically implemented they cannot make normative predictions of perceptual uncertainty.

To facilitate the study of explicit representations of uncertainty in rich real-world domains, we developed a class of neural network model that outputs the parameters of a distribution over the variables it estimates. These parameters can then be optimized to maximize the likelihood of the training data, forcing the model to accurately represent the true uncertainty of its estimates^28^.

We first applied the model to the problem of sound localization, which is rife with uncertainty due to the constraints of estimating a sound’s location in the world using only two microphones (the ears)^29^. We measured human estimates of uncertainty by asking participants to place bets on localization judgments. We then asked whether these judgments were reflective of the actual uncertainty of the estimated location by comparing them to judgments of uncertainty extracted from the model. We subsequently tested the generality of the framework by extending it to pitch perception. The results show that human confidence can accurately reflect uncertainty even in relatively unconstrained real-world settings. The modeling approach provides a general computational framework for incorporating uncertainty into working models of perception.

## Results

### Overview

In the sections that follow we motivate a computational framework for models of uncertainty. We then apply the model to two perceptual domains, in each case measuring human confidence judgments and asking whether they are consistent with the uncertainty predicted by a task-optimized model.

### Challenge of learning to represent uncertainty in neural networks

The dominant paradigm in deep learning is to train models to classify stimuli using a cross-entropy loss^30,31^. Such models typically have an output stage with different units representing different classes, and the unit activations are passed through a soft-max function that can be interpreted as a probability distribution over classes (Fig. 1a). Naively one might expect to be able to estimate the uncertainty of a classification decision from the width of this distribution. However, the cross-entropy loss function tends to cause such distributions to be biased in favor of the chosen class. Specifically, for any given training example, only the true class contributes to the loss, with the loss being minimized when the probability of the true class is maximized. The loss thus pushes a model to being overconfident (Fig. 1b), rather than representing the true uncertainty that may exist in a classification problem^25–27^.

**Figure 1.**
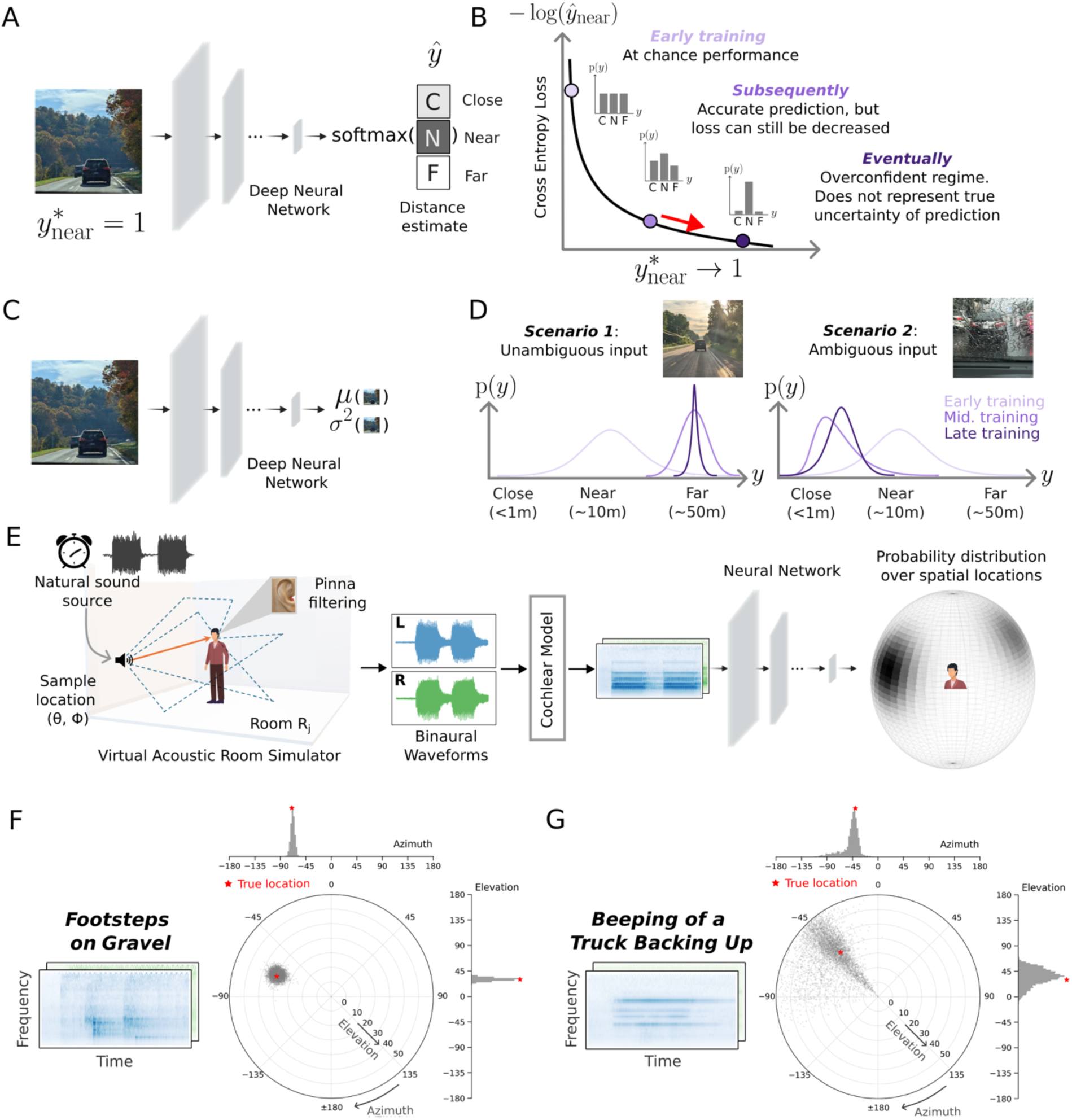
Conceptual sketch and application of the proposed computational framework to sound localization. **A**. Canonical neural network for classification. **B**. Schematic of consequence of training a model with the cross-entropy loss function (model becomes overconfident). **C**. Neural network that outputs stimulus-dependent parameters of a univariate Gaussian. **D**. When optimized with a maximum likelihood loss function, the model should learn to predict narrow distributions when actual uncertainty is low, and wider distributions when actual uncertainty is high. **E**. Model framework applied to sound localization. Training data is generated using a room simulator that yields the binaural audio signals that would enter the ears of a person at a particular location in a room listening to a sound at another particular location. The audio signals are passed through a fixed model of the cochlea followed by a neural network. The neural network predicts the parameters of a von Mises mixture distribution of locations over a sphere surrounding the listener. **F**. Posterior distribution for trained model for the sound of footsteps on gravel. This sound can be localized relatively precisely, and the model accordingly produces a narrow posterior. The posterior is visualized with Monte Carlo samples plotted in polar coordinates, with azimuth on the circular axis, elevation on the radial axis, and marginals shown as top (azimuth) and right (elevation) histograms. **G**. Posterior distribution for trained model for the beeping of a truck backing up. This sound’s location is relatively ambiguous, and the model accordingly produces a broader posterior.

### Challenge of estimating continuous variables in neural networks

A related challenge concerns the estimation of continuous variables. Deep learning models are universal function approximators and in principle can perform regression in addition to classification, producing a continuous valued output rather than a categorical decision. However, in practice, regression is more sensitive to outliers^32^, and tends to be less sample-efficient^33^ (i.e. requiring more training data to reach a criterion level of performance), often leading to worse performance than classification-based approaches. As a result, classification is commonly used even for problems involving inherently continuous variables^23,24,34,35^. One consequence of this approach is that the underlying topology of a continuous variable is lost: all erroneous classes are treated equally incorrect even when some may be much closer to the true value than others.

Because the topology defines similarity relationships within the target space, discarding it could lead to uncertainty estimates that fail to capture the graded structure of possible outcomes.

### Computational framework for stimulus-computable uncertainty representation

To build models that can learn to represent the actual uncertainty of a variable given a stimulus, it was necessary to enable a model to produce a distribution whose properties could vary depending on the stimulus (being tighter in some cases and broader in others), and to train the model using a loss function that would yield the correct distribution given the stimulus. We achieved this by treating the model outputs as parameters of a mixture distribution, and optimizing the model using a maximum likelihood loss function. This general idea was proposed by Bishop in 1994 for univariate mixtures of Gaussians^36^, but the neural networks that could be trained in that era were small, and could not be applied to real-world sensory problems. The idea has been revisited in the deep learning era for various engineering applications (including speech synthesis^37^, pose estimation^38^, and video prediction^39^), but has not been connected to human perception or cognition.

The principle underlying the approach can be understood in the simple setting of estimating a continuous, one-dimensional variable *y* from sensory input *x* (Fig. 1c, here representing the distance of an object from the camera, as an illustrative example). The negative log-likelihood of *y* under a univariate Gaussian distribution with mean and variance parameters predicted by a neural network ϕ is given as

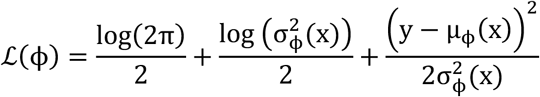

We can make two observations from this equation. First, if the variance term is not stimulus-dependent (i.e., constant), the loss reduces to the mean squared error objective standardly used in regression. Second, there are two complementary ways to reduce this loss. One way is to reduce the prediction error (i.e., to make µ_ø_(x) close to y). When the prediction error is close to zero, the loss can be further decreased by shrinking the variance (because in this regime, the effect of the variance on the 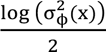 term will be greater than its effect on the 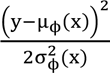 term). However, when the prediction error is large, the loss can be reduced by increasing the variance (decreasing the 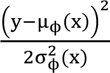 term), but only to the extent that it doesn’t outweigh the 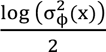 term. This balance couples accurate estimation of the mean with calibrated estimation of the variance, forcing a model to learn to produce small variance when the true value of the variable is unambiguous and can be estimated accurately, and larger variance when the variable is ambiguous and cannot be estimated with precision (Fig. 1d).

Although the actual distributions predicted by the models were more complicated than this univariate Gaussian (being mixture distributions), the underlying principle is this same balance between accuracy and uncertainty. The only labels the model received during training are point locations for each training example. However, the log-likelihood loss function will cause the model to learn to predict the true posterior distribution (in the limit of infinite training data, and successful optimization, and assuming the model architecture is sufficiently expressive). See Methods for a proof that the model’s output distributions will approximate the true posteriors given these assumptions. In this sense the model is an ideal observer for posterior distributions over point source locations given sound data drawn from the training distribution. In practice, the model is trained on a finite sample of training data, the model architecture places some limitations on what can be learned or expressed, and the optimization is not guaranteed to find a global minimum. However, the mixture distribution is sufficiently flexible that for large enough training sets the model should approximate the ideal observer for distributions over point source locations. We note that for sounds not well described as point sources (e.g. those that are spatially diffuse), the model cannot be an ideal observer, a point we return to later in the paper.

### Sound localization as a model problem for studying uncertainty

The framework is applicable to any perceptual variable, but we initially applied it to the problem of sound localization. This problem seemed like a useful test case for modeling uncertainty because the ambiguity of estimating location from only two sensors (the ears)^29^ creates considerable uncertainty in many real-world conditions. Moreover, internal representations of uncertainty seem likely to be important for behavior, as humans and other animals often base actions on where they think sounds are coming from^40–42^. In addition, sound localization is a relatively well studied perceptual problem, with a wealth of known perceptual effects that can be used to design experiments and validate models.

We first chose a functional form for the model output distribution that made sense for the problem, with the distribution defined over a sphere (centered at the listener’s head). Given an audio stimulus, the model produced the parameters of a von Mises mixture distribution. A von Mises distribution can be thought of as a circular analogue of a Gaussian distribution and so is a natural choice for a distribution over a sphere. The mixture distribution had 5 components, each defined by a two-dimensional mean and a two-dimensional concentration along with a mixture weight. Depending on the parameters, this distribution can be narrowly peaked at one location, broadly peaked, or multi-modal. The model was optimized for training data generated by rendering natural sounds in rooms at different positions relative to the listener (Fig. 1e; see Supplementary Table 1 for model architecture details). The sounds in the training data were superimposed on diffuse background noise.

The maximum likelihood loss function should cause the model to output narrow distributions in conditions of low uncertainty, and broader and/or multi-modal distributions in conditions of higher uncertainty. Inspection of the trained model’s output on example sounds suggested that the loss function had the intended effect. Fig. 1f&g plot the model posterior for two sounds: the sound of footsteps and the beeping of a truck backing up. The footsteps are broadband (with energy at a wide range of frequencies), and should be relatively accurately localized. Accordingly, the model posterior is narrow, with a peak near the true sound location. By contrast, the beeping truck has energy concentrated at a few frequencies, and would be expected to be difficult to localize^43^. The resulting model posterior is broad and multi-modal, with a peak in the front hemifield as well as in the back.

### Experiments 1 & 2: Model replicates accuracy of human sound localization

Because the model class was new, it was not clear a priori whether the model would perform well once optimized. We began by assessing localization performance, asking whether the model could replicate human localization performance with natural sounds. We obtained localization judgments from the model by taking the maximum a posteriori estimate from the posterior distribution predicted by the model (estimated by Monte Carlo sampling; see Methods).

We first evaluated model localization of natural sounds in quiet conditions (Supplementary Fig. 1a). Human listeners heard recorded natural sounds played from one speaker within a large array. The model performed the same experiment in a simulation of the experimental chamber. Model accuracy was close to that of human listeners and captured a substantial fraction of the variation in localization accuracy across different natural sounds (Supplementary Fig. 1b). We also evaluated the model with an experiment measuring localization of natural sounds from the same speaker array but accompanied by spatially diffuse background noise^20^ (Supplementary Fig. 1c), at several different signal-to-noise ratios (SNRs). Here the model again performed on par with human listeners, with a similar dependence on SNR (Supplementary Fig. 1d). Model performance was marginally worse than humans at low SNRs, plausibly because humans can learn the nature of the background noise over the course of the experiment^44^ (the model, by contrast, may sometimes erroneously base its judgment on the noise rather than the target sound). The model also replicated a range of classic signatures of human sound localization abilities (frequency-dependent use of interaural time and intensity cues^45^, dependence of bandwidth on accuracy^46^, use of high-frequency pinna cues to elevation^47^, the coarse spectral resolution^48^ and ear-specificity^49^ of such elevation cues^48^, and the precedence effect^50–52^; see Supplementary Fig. 2 for all these results, which replicate other recent neural network models of sound localization^20,23^). These results indicate that a model optimized to represent uncertainty can match the localization accuracy of human listeners.

### Logic of confidence experiments and analysis

As in prior work, we hypothesize that confidence is an estimate of the probability that a perceptual decision is correct^7^. Confidence should thus be a function of both the task an observer is asked to perform and of the uncertainty of estimated variables underlying the decision (e.g. the estimate of a sound’s location). To test whether humans estimate normatively correct representations of uncertainty, we asked whether human confidence judgments for sound localization were consistent with task-specific confidence judgments extracted from the model’s representation of uncertainty over location.

To our knowledge confidence judgments for sound location had not been systematically measured in any previous study. The one previous attempt of which we are aware measured confidence in an exploratory experiment with a relatively small sample size^53^, and did not find substantial variation in confidence with the stimulus conditions that were tested.

In a series of experiments described in the following sections, we concurrently measured conventional human localization or discrimination judgments along with confidence judgments. The main goal was to collect new human confidence judgments in order to compare them to those predicted by the uncertainty represented in the model posterior. We also measured accuracy to confirm that the participants performed as expected given prior work, and to confirm that the model reproduced human accuracy patterns. Based on the hypothesis that confidence should align with decision consistency^12,15,16^, we also measured localization precision across single-trial judgments, and compared it to confidence. We exclusively measured confidence judgments for azimuthal location, because it seemed likely that elevation judgments would be somewhat specific to an individual listener’s pinnae^54^.

In lieu of explicit judgments of confidence (e.g. ratings), we asked participants to place bets on their judgments. Participants were told that correct and incorrect bets would increase or decrease their overall compensation, respectively. Post-decision wagering is often recommended as a measure of subjective confidence, on the grounds that an observer seeking to maximize expected earnings should place bets proportional to their uncertainty in their decision^9^.

### Modeling confidence judgments

Confidence is often proposed to be an internal estimate of a decision’s reliability^12,15,16^. To extract model confidence judgments from the model’s distribution over location, we accordingly used the model’s posterior distribution to calculate a measure of uncertainty for the decision required by the task. For instance, for absolute localization judgments, we took the model confidence to simply be a summary measure of the spread of the model posterior. Whereas for experiments with binary judgments (front vs. back discrimination), we averaged the model posterior over front and back locations, and used the entropy of this binomial distribution. The resulting trial-level measure of confidence was subsequently mapped onto a 1–5 betting scale using an inverse cumulative distribution function (CDF) transform derived from the empirical distribution of the measure across the trials of an experiment.

### Experiment 3: Human bets reveal uncertainty representations for location

We first ran an experiment with synthetic sounds that we expected to vary in localization precision based on previous work^43,46^. Human listeners heard synthetic tones or noise bursts played from one speaker within a large array (Fig. 2a). On each trial listeners localized the sound within the horizontal plane (by selecting the location on a tablet) and indicated their confidence by placing a bet on their judgment (between 1 and 5 cents). The stimuli were selected to vary in the precision with which they could be localized, spanning broadband noise (high precision) narrowband noise (intermediate precision) and pure tones (single frequencies, with low precision). Localization precision is expected to increase with the source bandwidth^46^, other things being equal, as more localization cues become available. Localization precision is also expected to be higher at the midline, and lower at peripheral locations^55^, as binaural localization cues change more with position near the midline^29^. Example model posteriors are consistent with these differences in precision, and with the natural sound examples from Fig 1: the posterior for a white noise burst is unimodal and narrow, whereas that for a narrow-band noise is substantially broader (Fig. 2b).

**Figure 2.**
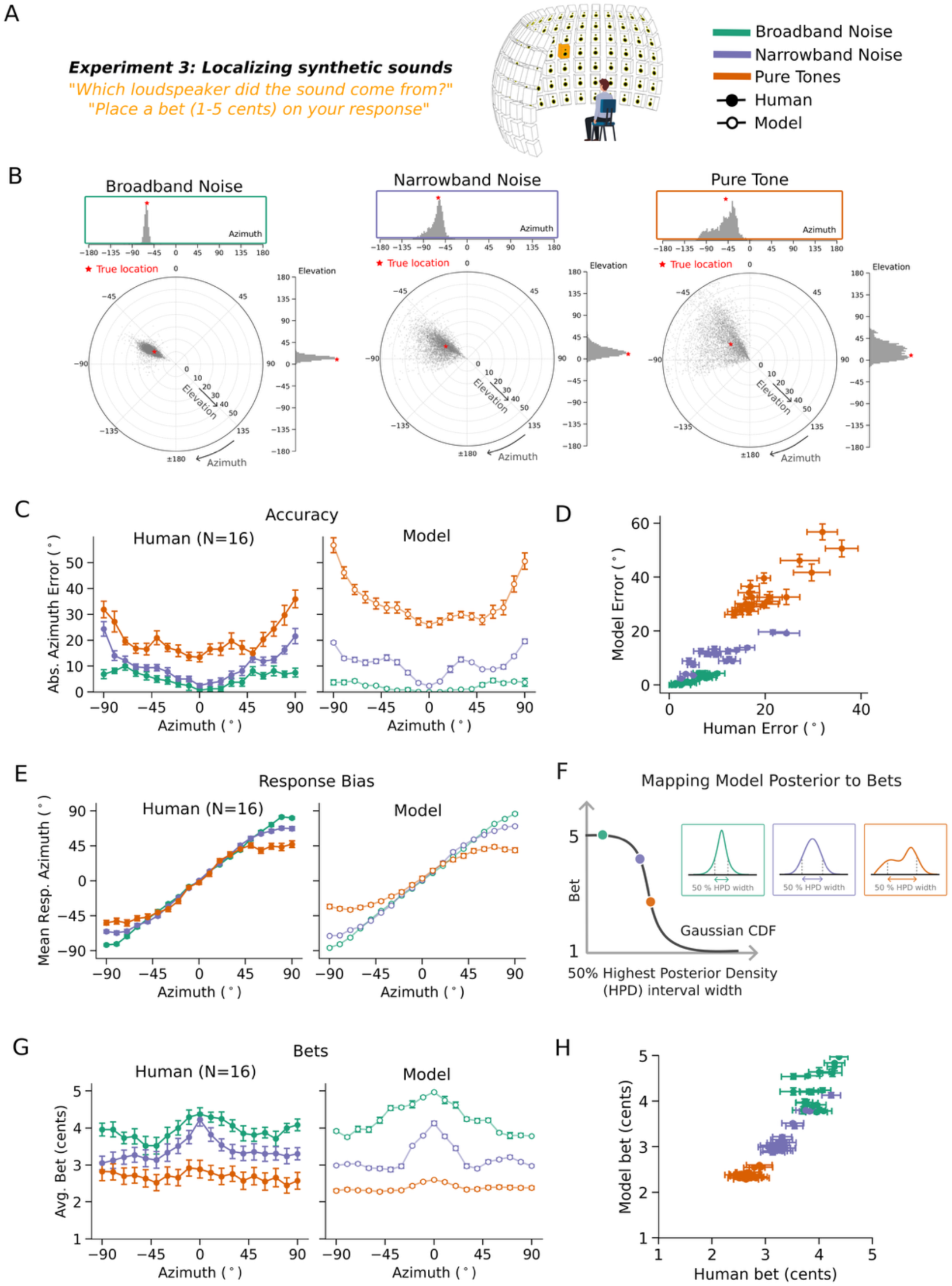
Experiment 3: Localization confidence judgments for synthetic stimuli in the frontal hemifield. **A**. Experiment measuring localization of noise bursts and tones in the frontal hemifield. Noise bursts and tones were presented from one of 133 locations, and the participant judged the azimuthal location. **B**. Example model posteriors for the three types of synthetic stimuli. The model posteriors are narrow for broadband noise, and broader for the stimuli that contain a smaller range of frequencies. **C**. Localization accuracy for each stimulus type at each azimuthal position. Here and in other panels, error bars plot standard error. **D.** Comparison of human and model localization error**. E**. Average judged location for each stimulus and azimuthal position. **F**. Model bets were generated from the width of the model posterior. **G**. Human and model bets for each stimulus type and azimuthal position. **H.** Comparison of human and model bets.

Human localization accuracy varied with stimulus and position as expected (Fig. 2c, left; main effect of stimulus type: F(2,30)=307.33, p<.001; main effect of position: F(18,270)=12.86, p<.001), replicating many previous experiments^46,55^. Results were similar for different pure tone and narrowband noise frequencies, and so are averaged across frequency here and in all subsequent analyses. Human localization judgments also showed biases at peripheral locations (Fig. 2d, left), as expected given prior work^56^. The average bet also varied with stimulus and position (Fig. 2e, left; main effect of stimulus type: F(2,30)=47.51, p<.001; main effect of position: F(18,270)=5.81, p<.001), but with the opposite trends. The accuracy effects are replications of many previous results, but the confidence results have not been previously reported.

### Experiment 3: Model replicates human confidence in azimuthal localization

Because the task in Experiment 3 was an absolute localization judgment, we simulated model confidence with a summary measure of the spread of the posterior distribution for a stimulus. As a summary measure we used the width of the 50% highest posterior density (HPD) interval^57^ (i.e., the width of the narrowest region containing 50% of the posterior mass; see Methods, and see Supplementary Figure 3 for results with alternative summary measures). Higher values of this measure should correspond to less reliable decisions, and thus lower confidence.

The model results replicated human results both for accuracy (Fig. 2c, right) and bias (Fig. 2d, right), but also for bets (Fig. 2f, right) with high human-model correlation for all measures (accuracy: r=0.91, p<.001; bias: r=0.99, p<.001; bets: r=0.94, p<.001). This latter result suggests that human confidence judgments are consistent with the actual uncertainty of their localization percept, on the assumption that the posteriors produced by the model correctly indicate this uncertainty.

### Experiment 4: Humans exhibit front-back uncertainty

The front-back dimension of sound localization is also known to have considerable ambiguity^40^. Humans have some ability to differentiate front and back locations, but often make errors^58,59^, in part because interaural cues can be identical for corresponding pairs of front and back sound locations (due to geometric symmetry). We assessed whether humans represent ambiguity in this setting by again measuring bets along with localization judgments.

Humans heard noise bursts or tones played from positions in front or behind the head (Fig. 3a). On each trial participants made a binary front-back judgment and placed a bet on their judgment. Performance was quantified as d’, calculated separately for each position (from 0 to 80 degrees). As expected^60^, human front-back accuracy was higher for broadband noise than tones (Fig. 3b, left; main effect of stimulus type: F(1,9)=578.15, p<.001). Human bets were also higher for noise bursts than for tones (Fig. 3c, left; main effect of stimulus type: F(1,9)=58.80, p<.001). This result has not been previously reported. Human bets were also higher at central than peripheral locations, as in Experiment 1 (main effect of position: F(8,72)=2.79, p<.01).

**Figure 3.**
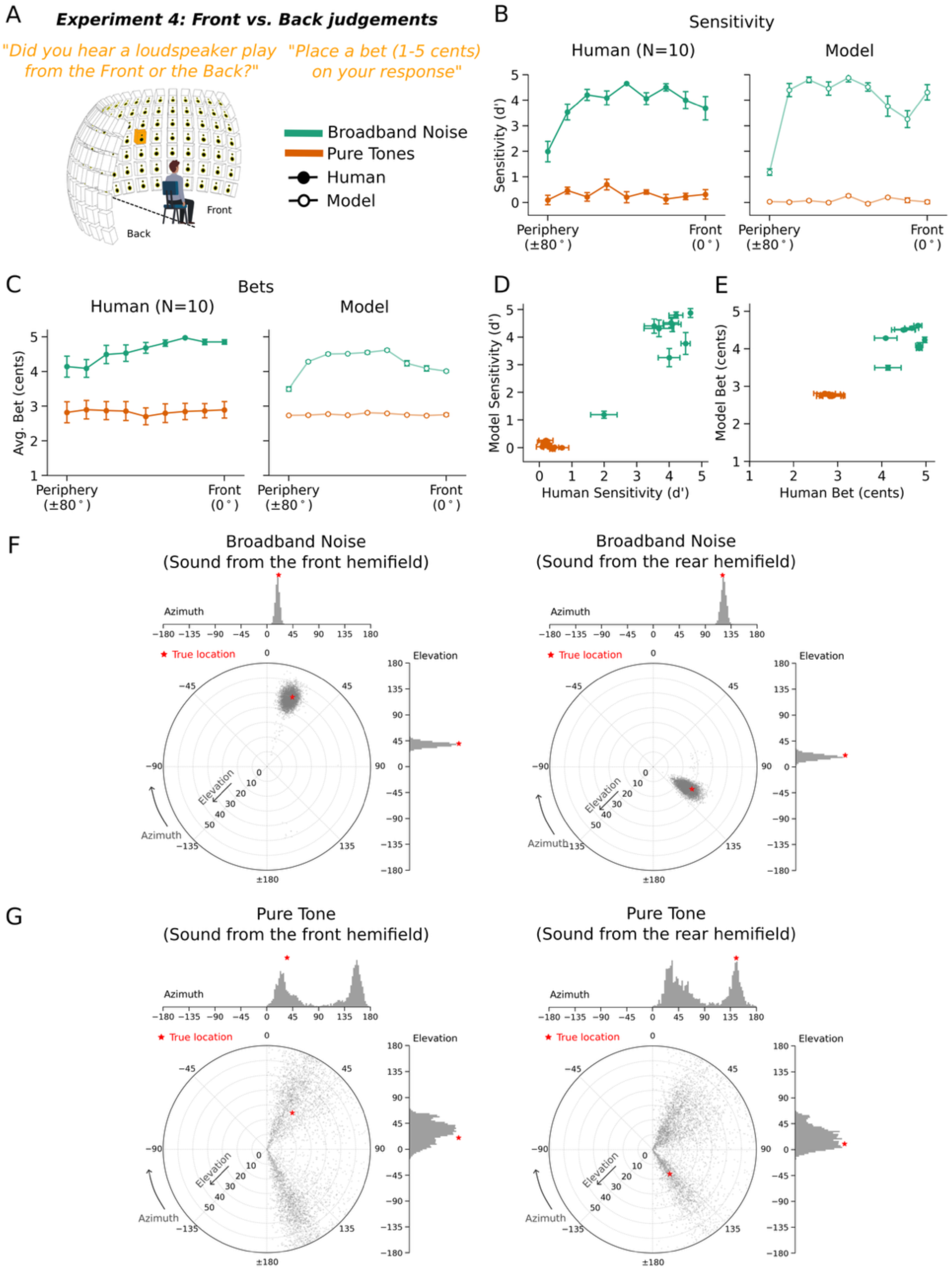
Experiment 4: Confidence judgments for front-back localization. **A**. Experiment measuring front-back localization of noise bursts and tones. Stimuli were presented in the left hemifield of the listener. The participant judged whether the sound was in the front or rear quadrant. **B**. Discrimination accuracy for each stimulus type at each azimuthal position. Here and in C, error bars plot standard error. **C**. Human and model bets for each stimulus type and azimuthal position. **D&E**. Comparison of human and model sensitivity and bets. **F**. Example model posteriors for noise bursts presented in a front or back location. The noise location is relatively unambiguous, and the model posteriors are unimodal. **G**. Example model posteriors for pure tones presented in a front or back location. Here the model posteriors are multimodal, reflecting the substantial front-back ambiguity for this stimulus type.

### Experiment 4: Model replicates human confidence in front-back localization

Because the task involved a binary judgment, we derived model confidence from the uncertainty of this judgment given the model’s posterior. Specifically, we integrated the posterior over front and back locations to yield a binomial distribution, and took the model’s confidence to be the entropy of this distribution. The model results qualitatively replicated human results in both front-back accuracy (Fig. 3b, right) as well as bets (Fig. 3c, right). Lower human bets coincided with wider and bimodal model posteriors, typically with peaks at both front and back (Fig. 3d&e). Human-model correlations were high for both sensitivity (r=0.97, p<.001) and bets (r=0.95, p<.001). We note that an alternative measure of the uncertainty of the full posterior across all locations did not reproduce human confidence (Supplementary Fig. 3b), underscoring the importance of a measure that specifically captures the uncertainty over the decision variable.

### Experiments 5&6: Model predicts human confidence in natural sound localization

To further test whether the model captured human confidence in natural conditions, we obtained model-derived uncertainty estimates for 160 unique natural sounds (Fig. 4a). From these predictions, we selected twenty sounds: 10 with high uncertainty and 10 with low uncertainty. We avoided the sounds with the highest and lowest uncertainty to avoid floor and ceiling effects on bets (and accuracy) and so selected the 10 sounds whose uncertainty was closest to the 5th and 95th percentiles of the model’s uncertainty distribution, respectively. Because we planned to run an absolute localization experiment, we used the same measure of uncertainty used to simulate model bets in Experiment 3 (the width of the 50% highest posterior density (HPD) interval). We then ran an experiment with human participants to assess their confidence in localizing these sounds, by obtaining both localization judgments and bets. We ran the same experiment on the model.

**Figure 4.**
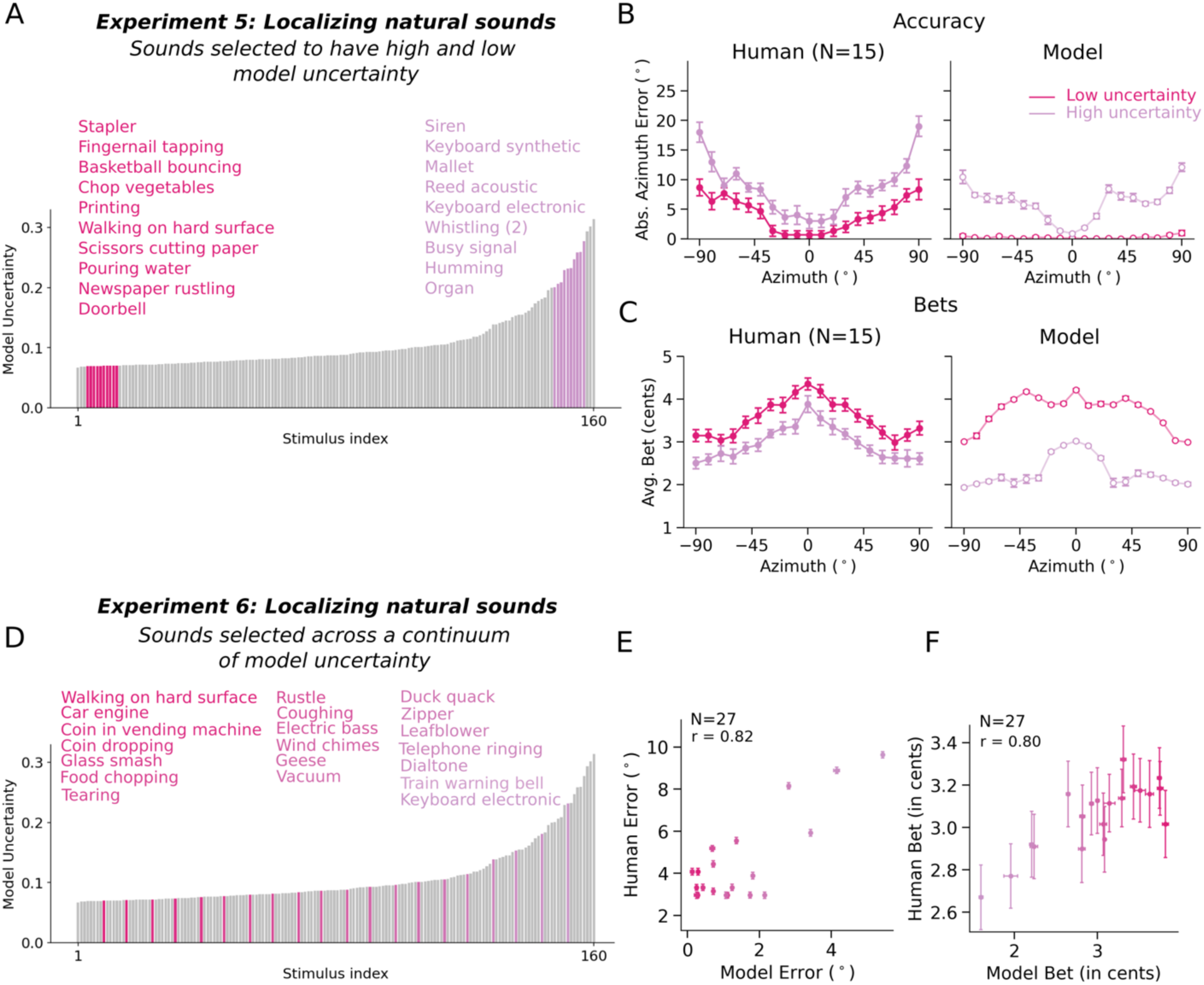
Experiments 5&6: Human and model confidence judgments for natural sounds. **A**. Model sound localization uncertainty for 160 natural sounds. For Experiment 5, we selected 10 sounds with low uncertainty and 10 sounds with high uncertainty and used them in an experiment with human participants. **B**. Results of Experiment 5: human and model localization accuracy for low- and high-uncertainty sounds. **C**. Results of Experiment 5: human and model bets for low- and high-uncertainty sounds at each azimuthal position. **D**. For Experiment 6, we selected 20 sounds that were evenly spaced when ordered by model uncertainty. Here and in E and F, color denotes model uncertainty. **E**. Results of Experiment 6: human and model localization accuracy for each sound (averaged across azimuthal position). **F**. Results of Experiment 6: human and model bets for each sound (averaged across azimuthal position).

Human localization accuracy differed systematically between low- and high-uncertainty stimuli (Fig. 4b, left; F(1,14)=181.45, p<.001), with accuracy also better at the midline compared to the periphery, as expected (main effect of position: F(18, 252)=18.93, p<.001). The average human bet showed the opposite trend, with sounds and locations with lower model uncertainty producing higher bets, and vice versa (Fig. 4c, left; main effect of stimulus group: F(1,14)=57.85, p<.001; main effect of position: F(18,252)=21.68, p<.001). The model results showed similar trends (as expected given how the sounds were selected; Figs. 5b&c, right).

**Figure 5.**
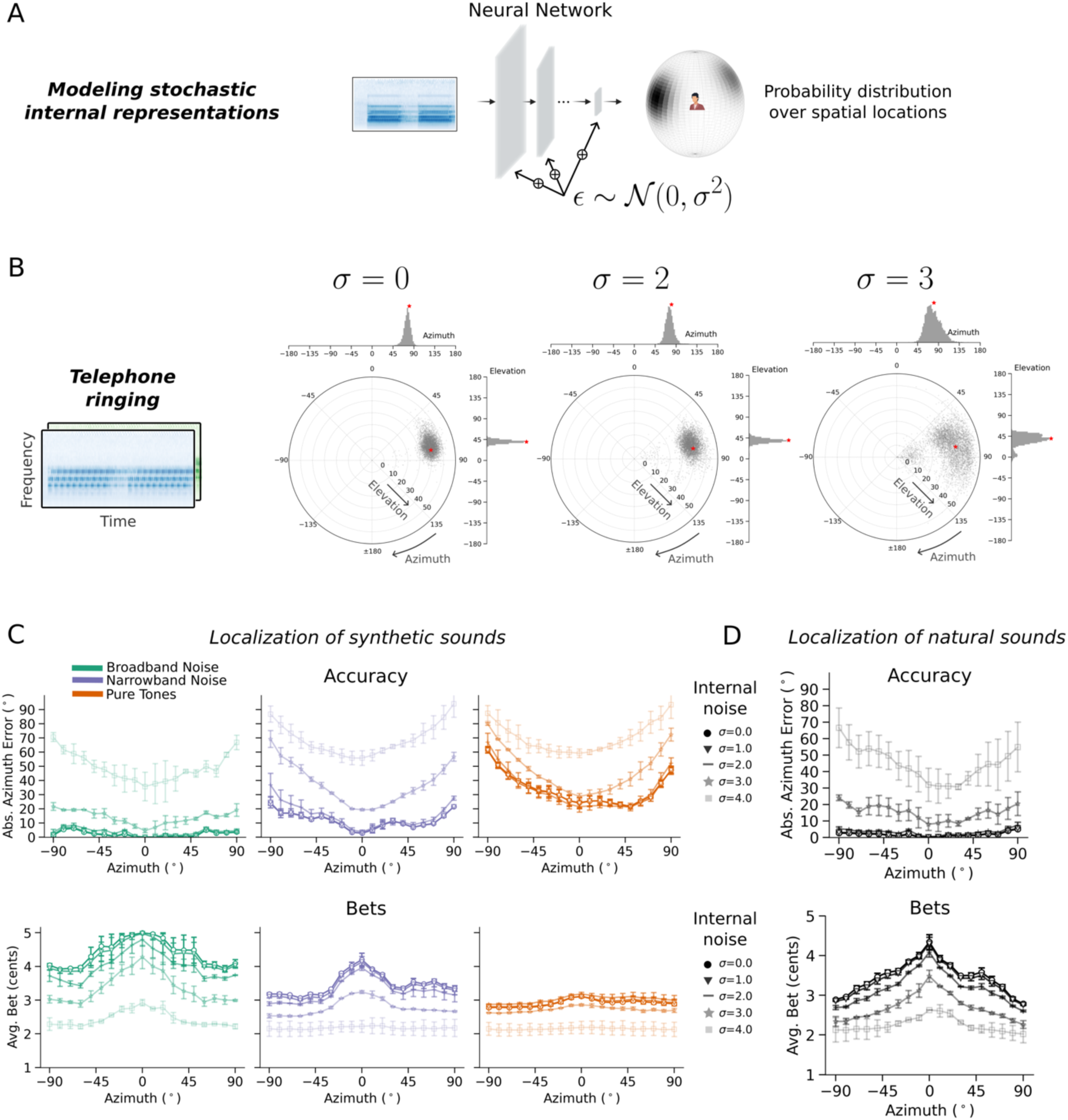
Effect of internal noise on model representations of uncertainty. **A**. Internal noise was added to model representations during training and evaluation. **B**. Example posteriors for different levels of internal noise. **C**. Model accuracy and bets for Experiment 3, simulated with different levels of internal noise. **D**. Model accuracy and bets averaged over the full set of natural sounds from which the stimuli in Experiments 5 and 6 were drawn, simulated with different levels of internal noise.

To test whether the human-model correspondence would persist for sounds drawn across a continuum of model uncertainty estimates, we conducted an additional experiment (Experiment 6) in which we selected twenty sounds evenly spaced from the 5^th^ to the 95^th^ percentile of model uncertainty (using the same set of 160 natural sounds; Fig. 4d). The sounds were again presented at different locations, and participants made localization judgments along with bets. As shown in Fig. 4e&f, localization accuracy and confidence vary smoothly across the uncertainty spectrum, with fairly good correspondence between human and model bets (r=0.80, p<.001; split-half reliability of the human bets: r=0.84).

Overall, these results suggest that humans represent location uncertainty even for complex natural sounds, and that these uncertainty estimates are captured by the model.

### Model uncertainty varies across many different stimulus manipulations

To assess whether the model’s uncertainty over sound location could potentially be explained by a small number of heuristics, we generated bets for the battery of classic experiments used to validate the model’s localization behavior in Supplementary Fig. 2. These experiments show that model bets vary not only with sound bandwidth^46^, but also as high-frequency pinna cues to elevation are removed with filtering^47^, if these cues come from ears that the model was not trained with^49^, and as the spectral resolution of such cues is reduced^48^ (Supplementary Fig. 4). These results suggest that uncertainty is influenced by any stimulus variable that affects the ability to infer location, rather than being dictated by a small number of heuristics.

### Model captures internal as well as external sources of uncertainty

In the version of the model presented thus far, the model’s representation of uncertainty is a deterministic function of the stimulus, exclusively reflecting external sources of uncertainty. However, human confidence is known to also reflect internal sources of uncertainty. For instance, the precision of the represented stimulus varies from trial to trial even when the stimulus is held constant, with humans reporting higher confidence on trials with greater representational precision^61^. To investigate whether the modeling framework could also account for internal sources of uncertainty, we trained a variant of the model in which noise was added to the model activations (Fig. 5a). The noise level varied across training examples. Once trained, we then ran the model in Experiment 3. We also measured accuracy and bets for the full set of 160 natural sounds from which the stimuli in Experiments 5 and 6 were drawn (those shown in Fig. 4).

Fig. 5b shows example posterior distributions with different levels of internal noise. The distributions become broader with higher levels of noise. Fig. 5c&d show accuracy and bets from Experiment 3, and for the set of 160 natural sounds, simulated with different levels of noise. Internal noise causes accuracy to degrade, and model uncertainty to increase. This result suggests that the modeling framework can capture the effect of internal sources of uncertainty, enabling it to account for another key property of human confidence. The deterministic formulation of the model used in most of this paper should thus be interpreted as a special case (i.e., zero internal noise) within a broader framework.

### Human confidence is not reproduced by a model trained with cross-entropy

The modeling framework described here was motivated by known shortcomings of the uncertainty representations of models trained to perform classification using a cross-entropy loss^25–27^. To confirm that such models would not suffice to explain human confidence, we trained a standard classification model (referred to henceforth as the “softmax” model) to perform sound localization. The softmax model had the same architecture and was optimized over the same training data as the uncertainty-aware model, differing only in its output stage and training objective. Sound location was discretized into 504 bins, and the model’s output stage had a unit for each of the 504 classes, the activations of which were passed through a softmax function to yield a categorical probability distribution. The model was optimized using a standard cross-entropy loss (as in previous neural network models of sound localization^20,23^).

Once trained, we ran the softmax model in Experiments 3-5. The model’s classification decision provided its localization judgment, and we extracted a model confidence judgment from the softmax distribution over its output activations. Because the distribution was over discrete classes, we took the model’s confidence to be the entropy over the possible location labels for Experiment 3 and 5. For Experiment 4, we measured the front-back entropy, as with the uncertainty-aware model. We omitted Experiment 6 from this analysis as it did not have enough trials per condition (sound x azimuth) to give reliable condition-specific results. We then compared both models’ similarity to human accuracy and confidence across the three experiments (Pearson correlation). As shown in Fig. 6a&b, the uncertainty-aware model exhibited a significantly better match to the pattern of human accuracy across experimental conditions (r = 0.87 vs. r = 0.82, significantly different by a one-tailed paired bootstrap test, p<.001) as well as a substantially better match to human confidence judgments across conditions (r = 0.9 vs. r = 0.81, significantly different by a one-tailed paired bootstrap test, p<.001). Inspection of the results of Experiment 3 (Figure 6b) reveals that the softmax model exhibits fairly similar patterns of accuracy as the uncertainty-aware model but fails to reproduce the detailed pattern of confidence across conditions (specifically, its dependence on azimuthal position) seen in humans and in the uncertainty-aware model. See Supplementary Fig. 5 for softmax model results for Experiments 4 and 5.

**Figure 6.**
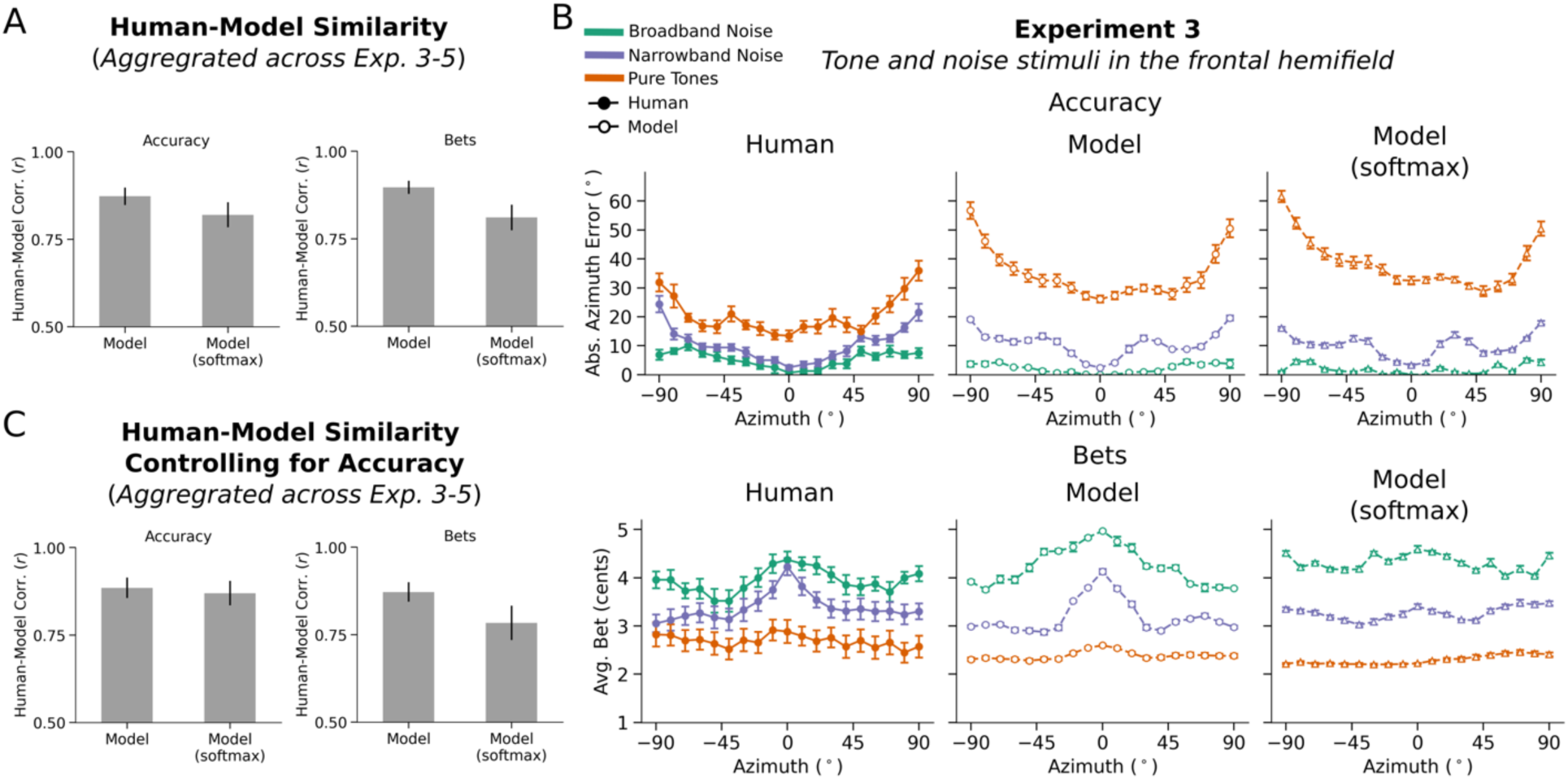
Comparison of human-model similarity between the main model featured in this paper (i.e., an uncertainty-aware model trained with a log-likelihood loss function) and a classification model trained with cross-entropy loss. **A**. Left: Aggregate human-model accuracy similarity (correlation between human accuracy and model accuracy across all conditions of Experiments 3-5). Error bars here and in B plot SEM, derived via bootstrap across conditions. Right: Aggregate human-model confidence similarity (correlation between human best and model bets across all conditions of Experiments 3-5). **B**. Comparison of uncertainty-aware model and classification model results for Experiment 3. Error bars plot SEM. See Supplementary Fig. 5 for softmax model results for Experiments 4 and 5. **C**. Human-model similarity for accuracy and bets for a subset of conditions selected in independent data to yield matched human-model similarity for accuracy. The difference between human-model bet similarity persists even for this subset of conditions.

To assess whether the improved match to human confidence was a side effect of better matching accuracy, we subsampled experimental conditions to match the human-model accuracy correspondence of the two models. We used half of the human participants and half of the model trials to select the subset of conditions, and then measured the human-model correspondence with the other half of participants and model trials. This resulted in human-model accuracy correspondence that was similar for the two models (Fig. 6c; r = 0.89 vs r = 0.87, though this difference reached statistical significance, one-tailed bootstrap test, p = .02). However, the human-model correspondence for confidence judgments remained substantially higher for the uncertainty-aware model (r = 0.87 vs r = 0.78, p = .005 via one-tailed bootstrap test).

Overall, these results provide further evidence that standard neural network models are not well suited to provide models of uncertainty representation, and that the modeling framework introduced here helps to fill this gap.

### Modeling uncertainty in human pitch perception

To assess the generality of our proposed framework for modeling perceptual confidence, we also applied it to the problem of pitch perception. Pitch is typically conceived as the perceptual correlate of the fundamental frequency (f0) of a sound^62,63^. Pitch is ecologically important given its central role in speech, music, animal vocalizations and environmental sounds. In practice, however, the f0 of many natural sounds may be ambiguous due to background noise, concurrent sounds, or the resolution limits of cochlear filters. This ambiguity makes pitch another useful test case for models of uncertainty.

As with localization, we chose a functional form for the model output distribution that was domain-appropriate. The model outputs were taken to be parameters of a distribution over the logarithm of the f0. Given an audio stimulus, the model produced the parameters of a Gaussian mixture distribution (5 components, each defined by a one-dimensional mean and variance along with a mixture weight), which could be narrowly peaked at one frequency, broadly peaked, or multi-modal, depending on the parameters (Fig. 7a). The model was trained on excerpts of speech and music superimposed on noise^24^.

**Figure 7.**
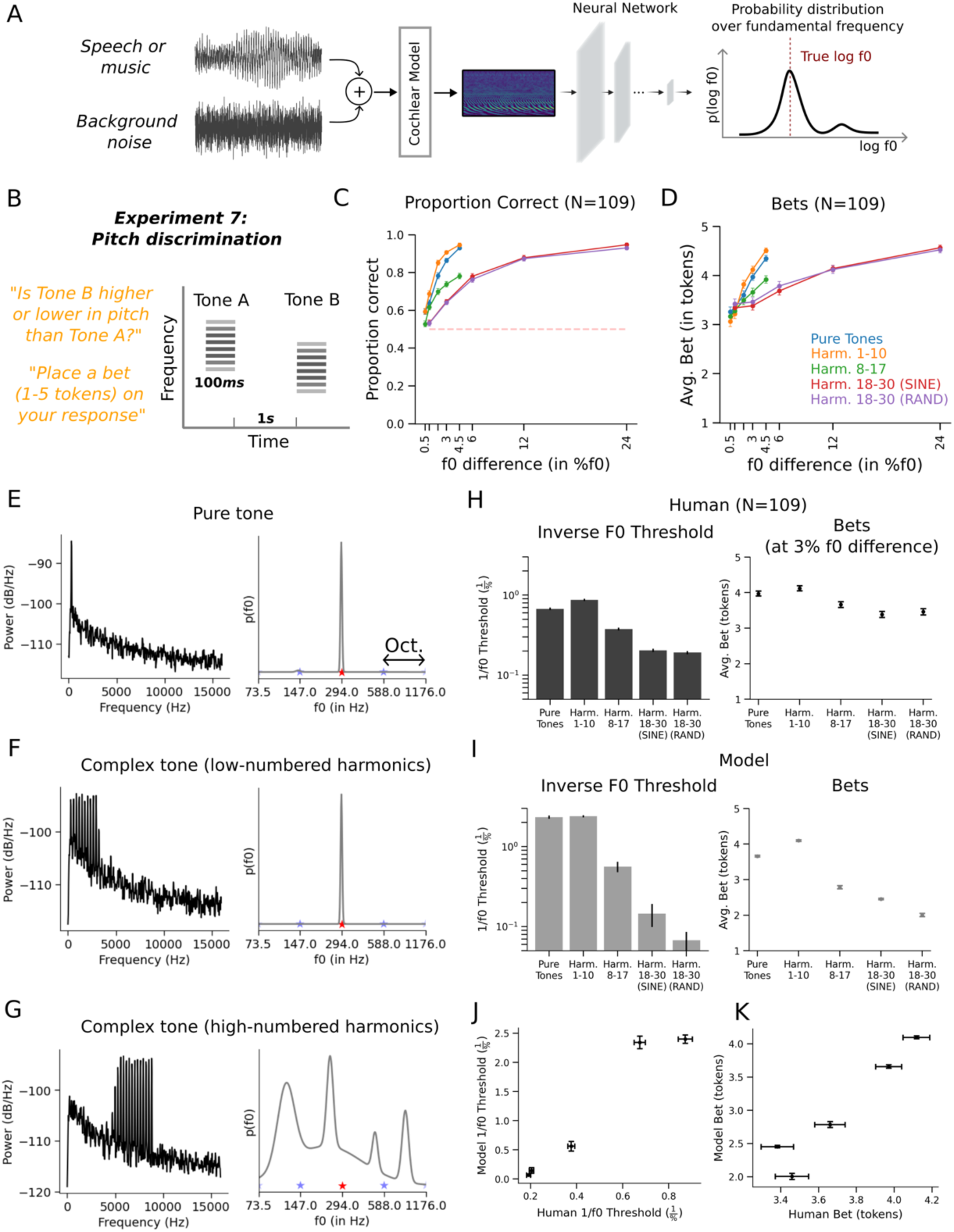
Experiment 7: Confidence in pitch perception. **A**. Model framework applied to pitch. Training data was generated by superimposing excerpts of speech or musical instrument notes on background noise. Audio signals were passed through a fixed model of the cochlea followed by a neural network. The neural network predicts the parameters of a Gaussian mixture distribution over log-frequency. **B**. Pitch discrimination experiment. Participants heard two tones, judged whether the second tone was higher or lower in pitch than the first tone, and placed a bet on their judgment. **C**. Average discrimination performance of human participants in each experimental condition. Here and elsewhere, error bars plot standard error. **D**. Average bet placed by human participants in each experimental condition. **E**. Stimulus spectrum for a pure tone (top) and the corresponding output distribution for trained model (bottom). Here and in F&G, red asterisk plots true f0, and blue asterisks plot f0s an octave above and below true f0. All stimuli were embedded in noise intended to mask distortion products that might otherwise provide a cue to the f0. **F**. Stimulus spectrum for a complex tone with low-numbered harmonics (top) and the corresponding output distribution for trained model (bottom). **G**. Stimulus spectrum for a complex tone with high-numbered harmonics (top) and the corresponding output distribution for trained model (bottom). **H**. Human f0 discrimination thresholds (left) and average bets (right). Bets are plotted for the 3% f0 difference, which was run for all tone types. **I**. Model f0 discrimination thresholds (left) and average bets. **J**. Comparison of human and model thresholds. **K**. Comparison of human and model bets.

Once trained, we assessed the model on a set of classic signatures of human pitch perception. As with previous neural network models of pitch^24^, the model reproduced the human dependence of pitch discrimination on harmonic number^64^, the effect of harmonic phase on pitch for stimuli containing only high-numbered harmonics^65^, the shifted pitch for shifted inharmonic complex tones^66^, poor discrimination of transposed tones^67^, and the pitch shift from mistuned harmonics^68^. These results, shown in Supplementary Fig. 6, indicate that the model captured the accuracy dependence of human pitch perception.

### Experiment 7: Human bets reveal uncertainty representations for pitch

To our knowledge human confidence judgments had not previously been measured for pitch perception, and so we measured them using an experiment with human participants. The stimuli were synthetic sounds intended to vary in discriminability (based on extensive previous work^64,65,69^). The question was whether the stimuli would produce variation in confidence, and whether any such variation would be mirrored by the model. We used a discrimination task based on the expectation that relative pitch judgments would be more natural for untrained human participants than absolute pitch judgments. On each trial, participants heard two tones in succession (in noise, to mask potential distortion products^70,71^) (Fig. 7b). They judged whether the second tone was higher or lower in pitch than the first and indicated their confidence by placing a bet on their judgment (between 1 and 5 tokens). Participants were again told that correct and incorrect bets would increase or decrease their overall compensation, respectively.

The stimuli consisted of pure tones and harmonic complexes with one of three different ranges of harmonics (low-, medium-, or high-numbered). In the high-numbered harmonic condition, we additionally varied the phases of harmonics to be aligned in sine phase or randomized. Pitch perception is known to be worse for high-numbered harmonics^64,65,69^, for which discrimination also tends to depend on phase^64,65^ (which affects the salience of the envelope fluctuations believed to underlie discrimination in such conditions). But unlike sound localization, pitch discrimination with pure tones is relatively good provided the frequency is not too high or too low^72^. We measured discrimination performance for each stimulus type for a range of f0 differences chosen to span near-change to near-ceiling performance for each condition.

Human discrimination varied with stimulus type as expected, replicating many previous experiments^64,65^, with the exception of the absence of a phase effect for these stimuli, for reasons that are not clear (Fig. 7c). Human bets also varied with the stimulus condition, being higher for larger f0 differences, but also varying across tone conditions (Fig. 7d). This result has not been previously reported.

### Experiment 7: Model uncertainty replicates human uncertainty for pitch

To compare model performance to that of humans, we simulated trials for each tone type with a range of f0 differences and generated psychometric functions. To yield a single summary measure of discrimination performance that could easily be compared to that of humans, we estimated thresholds from the average psychometric function for each tone type.

The dependence of human bets on both stimulus type and f0 difference exposes the distinction between confidence and the uncertainty associated with individual stimuli: human confidence judgments clearly reflect the uncertainty of the decision required by the task (because they scale with the f0 difference between the two tones on a trial) in addition to any uncertainty over the estimate of the f0 of each tone. To simulate model confidence judgments, we could have attempted to calculate the uncertainty of the discrimination decision from the model posterior for each of the tones on a trial. However, this calculation is nontrivial due to the mixture distributions that are involved. Instead, because it seemed obvious that the uncertainty of the decision would scale monotonically with the f0 difference, we compared human and model confidence for only a single f0 difference, estimating the model’s confidence from its uncertainty over the f0 of individual tones of each type (using the variance around the posterior mode as a summary measure; see Methods). We then compared this model confidence estimate to human bets for a fixed f0 difference for each tone type. We chose the f0 difference of 3% as this produced above-floor and below-ceiling performance for all stimulus types. Example model posterior distributions showed higher uncertainty for some tones than others (Fig. 7e-g).

As shown in Fig. 7h-k, the model results largely replicated the human dependence of discrimination and confidence on stimulus type, with better performance and higher bets for stimuli with lower-numbered harmonics. One qualitative difference was evident in the results for the random phase condition, which yielded lower model performance than the sine phase condition. We do not have a definitive explanation for this discrepancy, but it is possible that it reflects the statistics of the model’s training data, which might have prominent envelope fluctuations, making the model especially sensitive to this manipulation.

Overall, these results demonstrate the generality of the uncertainty modeling framework and again suggest that human confidence judgments derive from accurate estimates of the actual uncertainty of the variables being estimated.

### Human and model confidence track perceptual precision

The similarity between human confidence judgments and measures of uncertainty derived from the model posterior lends support to the idea that human confidence judgments reflect estimation uncertainty. However, estimation uncertainty should also be reflected in objective measures of performance across multiple trials of the same stimulus. Specifically, uncertainty should limit discrimination of stimuli, and in absolute estimation tasks such as localization, it should determine the consistency of judgments across trials.

Figure 8 presents an explicit comparison of human and model bets to the relevant measure of objective performance: precision, for the two absolute localization experiments (i.e. the variance of the location estimate for a stimulus, independent of any bias), sensitivity, for the front-back discrimination experiment, and thresholds, for the pitch discrimination experiment. In each case the bets were strongly correlated with the objective measure of estimation precision (Experiment 3: Human (r=-0.93, p<.001), Model (r=-0.92, p<.001); Experiment 4: Human (r=0.97, p<.001), Model (r=0.99, p<.001); Experiment 5: Human (r=-0.95, p<.001), Model (r=-0.94, p<.001); Experiment 7: Human (r=0.99, p<.01), Model (r=0.97, p<.01)). This result further supports the idea that confidence judgments in these experiments indeed reflects estimation uncertainty. See Supplementary Fig. 7 for human and model precision as a function of azimuth for Experiments 3 and 5.

**Figure 8.**
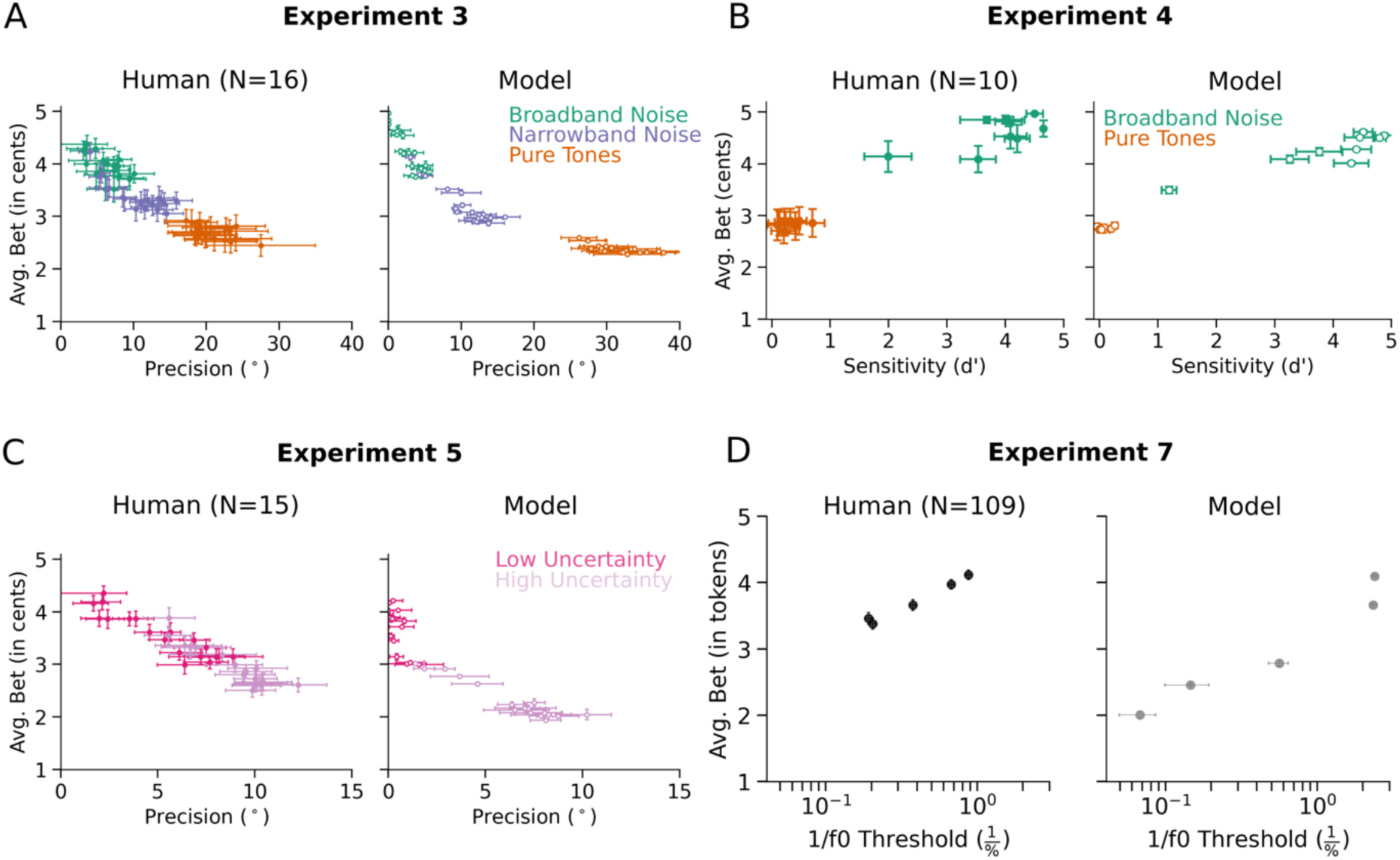
Relationship of confidence to precision. Each panel plots a measure of response precision vs. the average bet for each condition of an experiment. Error bars plot standard error. We note that colors indicate classes of conditions (mirroring the conventions of the other results figures in this paper), and that the individual dots depict different locations, such that each dot represents a distinct stimulus. **A.** Experiment 3: Tone and noise stimuli in the frontal hemifield. **B.** Experiment 4: Front-back localization of tones and noise stimuli. **C.** Experiment 5: Natural sounds screened by model uncertainty. **D.** Experiment 6: Pitch discrimination of tones.

We note that a critical distinction between measures of objective precision and confidence is that precision is only evident across multiple trials. By obtaining judgments over multiple trials with the same stimulus, we can indirectly infer estimation uncertainty. But a confidence judgment (assuming it accurately reflects uncertainty) provides access to uncertainty at the level of a single trial. Indeed, that is why access to representations of uncertainty is presumably useful to an observer.

### Humans and model exhibit metacognitive sensitivity

Under many conditions human confidence judgments reliably distinguish better from worse decisions within the same experimental condition^73,74^. To ask whether this “meta-cognition” was evident in the model as well, we re-analyzed the experiments, splitting the trials for each condition into those with high and low bets (this analysis was only possible for Experiments 3, 4, and 7, as the other experiments did not have enough trials per conditions for individual participants). As shown in Figure 9, performance was systematically better on high-bet trials than on low-bet trials. Notably, this was true for both humans (Experiment 3: F(1,14)=31.08, p<.001; Experiment 7: F(1,92)=223.53, p<.001; the trend was also clear in Experiment 4 but only five of the participants had data in all the confidence/condition bins, precluding a statistical test), but also the model (Experiment 3: F(1,4)=301.72, p<.001, including only the narrowband and pure tone conditions; Experiment 4: F(1,4)=1967.28, p<.001; a statistical test was likewise not possible for Experiment 7 as there were no low-bet model trials for some conditions). In addition, we calculated several proposed metrics of metacognitive ability (meta-d’/d’^75^, the Brier score^76^, confidence delta^77^, and expected calibration error^77^) from the responses for Experiments 4 and 7 (as these experiments used binary judgments as required by the metrics) where possible (see Methods). These analyses indicate that both humans and the model are metacognitively sensitive (Supplementary Table 2). We note that the model has complete access to its posterior, which could explain why it tends to be more meta-cognitively sensitive than humans in these analyses.

**Figure 9.**
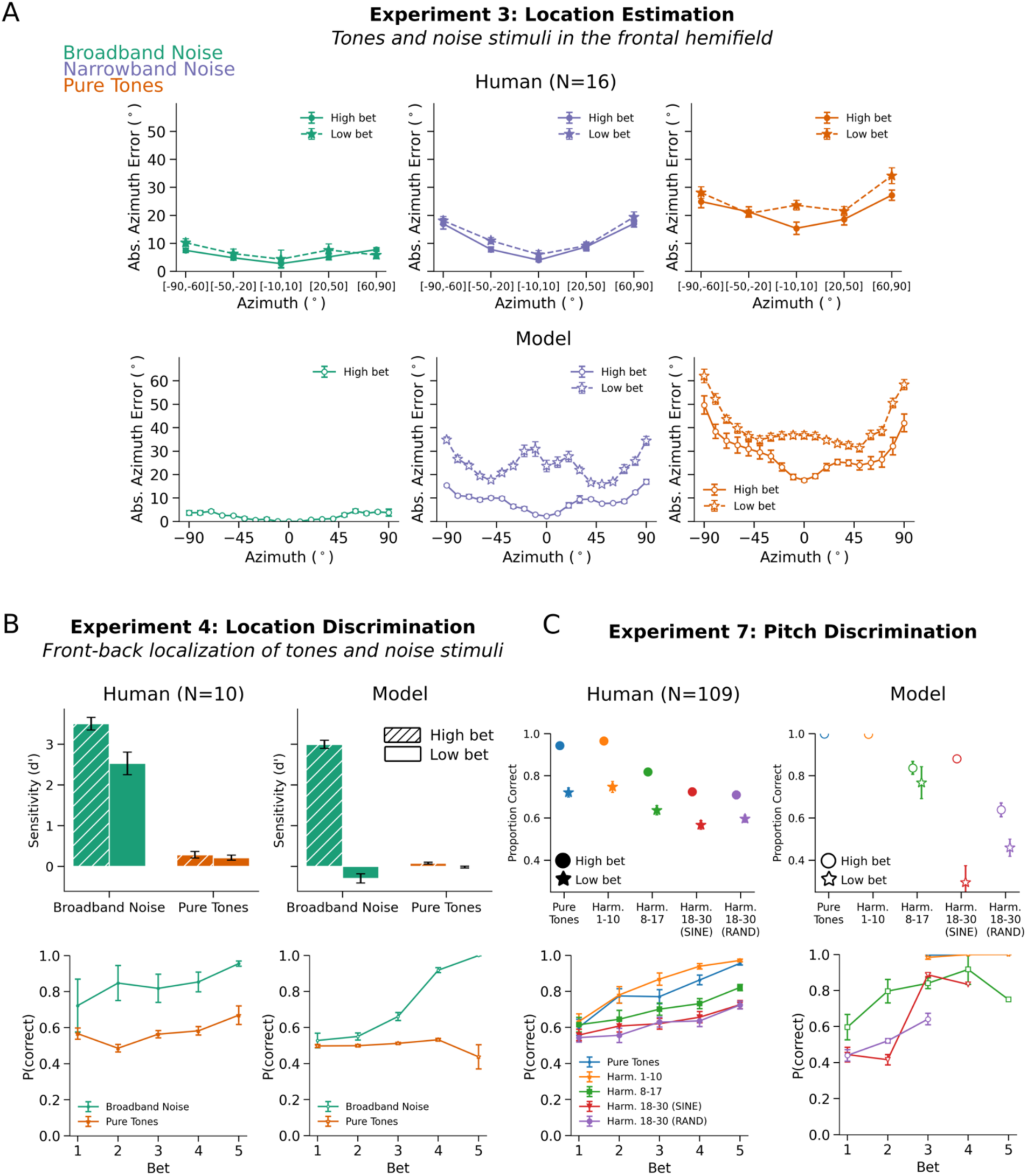
Comparison of human and model meta-cognitive sensitivity. In each experiment in which there were enough trials per condition to perform the analysis, trials for each condition were split into those with high and low bets. **A**. Results of Experiment 3, split into high- and low-bet trials. Here and elsewhere, error bars plot SEM. Note that there were no low bets for the model for broadband noise. **B**. Results of Experiment 4, split into high- and low-bet trials (top), and for each of the five bet values (bottom). The more granular analysis here and in C was possible due to the large number of trials per condition, and appropriate given the 2AFC tasks used in these experiments. **C**. Results of Experiment 7, split into high- and low-bet trials (top), and for each of the five bet values (bottom). Note that the model did not have data in all of the bet/condition bins.

### Experiments 8&9: Uncertainty is distinct from spatial diffuseness

The models we present here represent a distribution over a single quantity, with the spread of the distribution representing the uncertainty of that quantity. This representation assumes that sounds are generated by point sources. However, objects in the world are often not point sources with respect to the various dimensions that define them. For instance, many sound sources are spatially diffuse (e.g. a swarm of bees), and others have a fundamental frequency that varies over some range (Fig. 10a). Similarly, visually presented objects have a location as well as a spatial extent.

**Figure 10.**
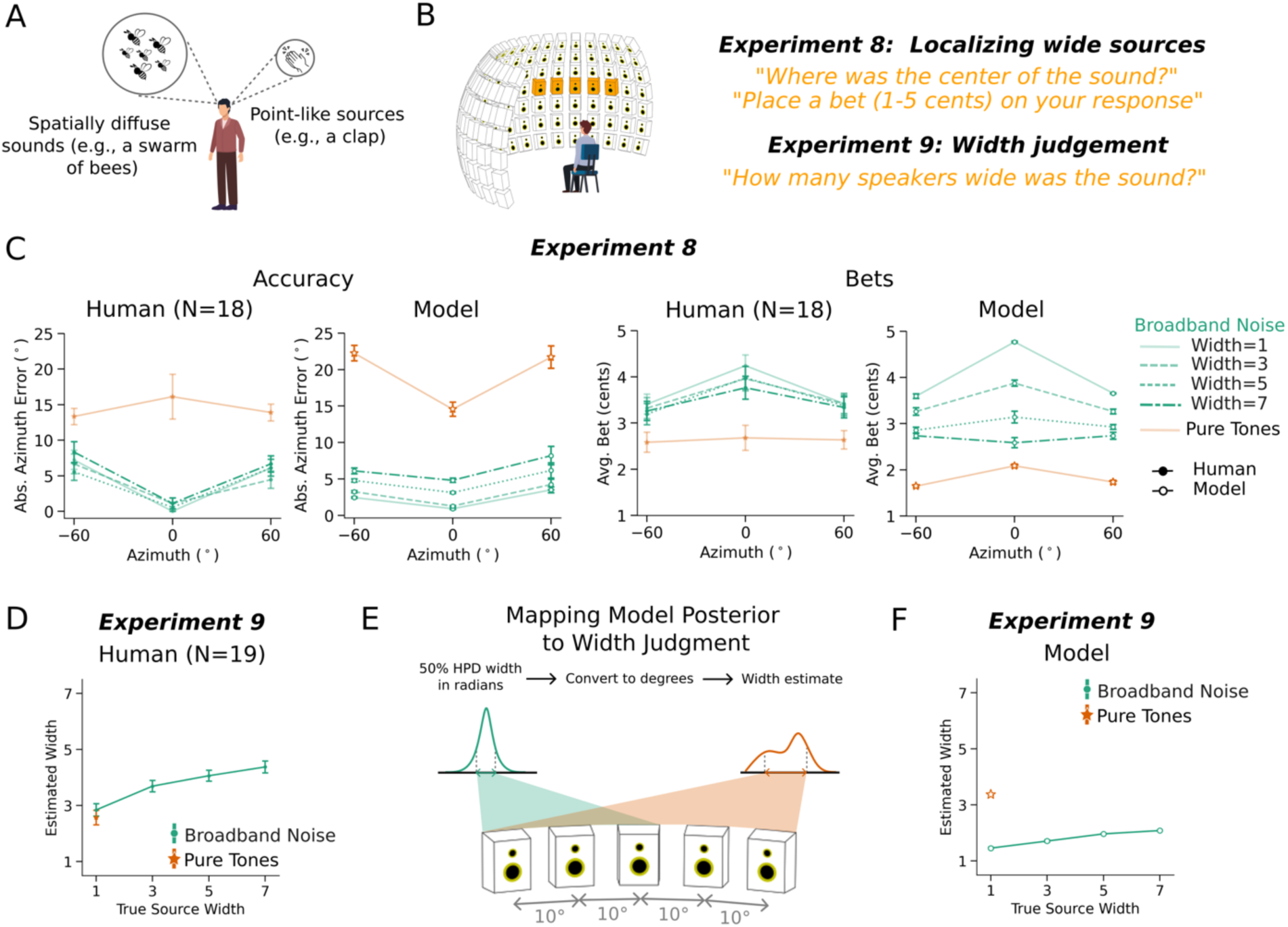
Experiments 8&9: Perception of source width is distinct from perception of location uncertainty. **A**. Sound sources in the world vary in their spatial extent. **B**. Width perception experiments. In Experiment 8, participants localized sources that could span up to 7 adjacent speakers, and placed bets on their location judgment. In Experiment 9, participants judged the width of the presented sound, by specifying the number of speakers from which the sound appeared to be played. **C**. Results of Experiment 8: localization accuracy (left) and average bet (right) for each stimulus condition, for humans and model. Here and in E, error bars plot standard error. **D**. Results of Experiment 9: average estimated sound source width vs. the true source width, for humans. **E**. Model “width” judgments were derived from the spread of the model posterior. **F**. Results of Experiment 9, for model. Because the model does not represent source width as distinct from the uncertainty of a source’s location, its “width” judgments simply reflect the uncertainty over location, unlike width judgments of humans.

Humans have some ability to perceive the spread of a sound source in space^78–81^ or other dimensions, but the relationship to representations of uncertainty is unclear. The models presented here brought this issue into focus, as they cannot distinguish between the width of a sound source along a dimension of interest and the uncertainty in that dimension, raising the question of whether humans can do so. We explored this question with respect to the spatial width of sound sources in the world, as it was straightforward for humans to report this width using a speaker array that we used in the preceding experiments.

We conducted two experiments. In Experiment 8, humans heard pure tones or noise bursts, judged the location, and placed a bet on their judgment (Fig. 10b). The noise bursts could be played from 1, 3, 5, or 7 adjacent loudspeakers (independent samples of noise from each speaker). When played from more than one speaker the noises were perceptibly wider in spatial extent. The tones were always played from a single loudspeaker (due to way that sinusoids superimpose, multiple pure tones from different speakers are physically indistinguishable from a tone from a single speaker). Participants reported the perceived center of the sound source.

In Experiment 9, a separate set of subjects heard the same stimuli from the same locations, but were told that the sounds could originate from multiple adjacent loudspeakers, and judged the perceived width of the sound (in speakers). The question was whether judgments of source width would dissociate from judgments of uncertainty (as reflected in the participant bets).

As shown in Fig. 10c, localization accuracy and bets for these stimuli showed the expected trends, with noises more precisely localized than tones, and with higher bets on noises than tones. The model replicated these qualitative trends. However, participants judged the physically wider noise stimuli to be wider than the pure tones (Fig. 10d). Thus, a stimulus perceived to have high uncertainty (pure tones) is not necessarily perceived to be spatially diffuse, and spatially diffuse stimuli are not necessarily perceived to have uncertain locations. These findings indicate that perceived width and location uncertainty are distinct perceptual dimensions.

The present model cannot represent this distinction as the distributions it outputs are over point locations. To underscore this point, we extracted “width” judgments from the model by estimating the spread of the posterior in degrees, and converting to the corresponding number of speakers (Fig. 10e). As expected, when the model’s output is interpreted in this way, it judges the pure tone to be wider than any of the noises, unlike humans (Fig. 10f). This result is not surprising given the limits on what the model can represent, but underscores that future models will need more expressive representations in order to account for the richness of human uncertainty.

## Discussion

We developed a computational framework for building sensory models that represent the uncertainty of perception. The models output the parameters of mixture distributions over the variables they estimate. We tested the framework on the problems of sound localization and pitch perception. We then used the resulting models to ask whether human confidence judgments are reflective of the actual uncertainty of perceptual estimation in these domains, based on the assumption that the models learn to correctly predict the true distribution over location or pitch given a stimulus. Human confidence, as measured by bets on individual trials, closely tracked the model’s uncertainty in both domains. The models also identified natural sounds producing percepts of high and low uncertainty, that in turn produced distinct levels of confidence in human observers. The results indicate that human perceptual confidence judgments are normatively appropriate, in the sense that they are consistent with the actual uncertainty of perceptual estimation in the conditions we tested. We also found that the uncertainty estimates of models trained with internal noise reflected both internal (determined by the level of noise) and external (determined by the stimulus) sources of uncertainty. The framework we propose is general and can be applied to any perceptual estimation problem, which should help enable the study of confidence in realistic perceptual domains.

### Relation to prior work on confidence

Our work builds on a large literature documenting confidence judgments in humans performing psychophysical tasks^7–12,15,16^. Our main contributions relative to this literature were 1) to measure confidence judgments in a real-world task (localization of sounds in space), 2) to develop a stimulus-computable model of perceptual estimation uncertainty that could make predictions in this real-world domain, 3) to demonstrate correspondence between human confidence judgments and the actual uncertainty of perceptual decisions in a real-world domain, 4) to show that the same modeling framework also accounted for confidence judgments for pitch, 5) to show that the same modeling framework could encompass internal as well as external sources of uncertainty, and 6) to demonstrate that perception of uncertainty in a dimension is distinct from perception of the spread of a source in that dimension. We also studied confidence within the sense of audition^82^, in contrast to most prior work on perceptual confidence, which has largely focused on vision. We chose to work with auditory tasks because they often feature a high degree of uncertainty in everyday listening conditions, and so seemed a good test bed for real-world theories of confidence.

### Relation to prior work on models of sensory systems

Our work also builds on a growing literature describing models of perception built using deep learning^18–20,22–24,83–85^. This line of work has repeatedly found that models optimized for real-world tasks replicate human performance characteristics, at least to first order, provided they are optimized under natural conditions^23,24^. This general result is consistent with the idea that humans are fairly well optimized for important natural tasks^86^, such that their behavioral characteristics are reproduced to a considerable extent by machine systems optimized for similar tasks^87^. The results here provide another example of optimized models converging to human-like behavior, in this case in the domain of confidence. The present work innovates on previous work using task-optimized models in two key respects.

First, the models we developed here give predictions over a continuous rather than discrete space. Prior models have largely used classification-based objectives, in some cases discretizing a continuous output dimension into bins to allow them to be fit into a classification framework^20,23,24^. This approach was necessitated by the poor performance of models trained with traditional regression objectives (in our experience it was difficult to achieve human levels of performance with such models). The present approach of predicting distributions over continuous variables in practice enabled high-performing regression-based models. This innovation should open the door to new models in a wide range of domains that involve the estimation of continuous variables.

Second, the models produced representations of uncertainty that were incentivized to be normatively correct. We trained the models with point labels for location or fundamental frequency (without explicit labels for uncertainty), and relied on the log-likelihood objective to drive the models to produce appropriate probability distributions (see Methods for proof).

### Relation to previous neural network models of density estimation

The idea of estimating mixture distribution parameters with a neural network was proposed over 30 years ago^36^, but at the time was limited to small-scale problems. The idea has been revisited sporadically in the deep learning era but has been restricted to engineering applications^37–39^ and has not made contact with work in perception and cognition. Our contribution was to integrate the idea of deep mixture density estimation with models of perceptual problems and to show how they can be used to understand confidence. Our models were fairly straightforward to optimize, and enabled large architectures trained on large datasets that in practice performed well on realistic problems, and seem a promising approach for future work in this area.

### Relation to literature on calibration and uncertainty estimation in AI

The alignment between a model’s confidence and the true likelihood of correctness, known as “calibration”, has become increasingly important as modern AI systems are deployed in safety-critical domains such as autonomous driving and medical diagnosis. A growing body of work has shown that most modern neural networks are poorly calibrated by default^88^, often producing overconfident estimates. More recently, similar effects of miscalibration have also been consistently observed in large language models^89–91^ (LLMs). Our results with softmax models trained on the domains we studied confirm these problems (Fig. 6). A range of post-hoc calibration techniques^88,92–94^ have been developed to adjust model outputs after training to better align with empirically correct uncertainty.

A parallel line of research has sought to model uncertainty directly either via novel training objectives^90,95,96^ or by training models to predict not only a point estimate but also the parameters of a higher-order distribution that captures the model’s uncertainty about its own predictions. This latter family of methods, known as evidential learning, has been developed for both classification^25^ and more recently for regression problems^97^. The latter approach (from Amini et al.) is most closely related to what we propose here. The main difference is that Amini et al. assume a stereotyped (single Gaussian) form for the model’s output distribution in order to be able to also predict a prior over distribution parameters, allowing their models to express second-order uncertainty (uncertainty about the predicted distribution itself). By contrast, our models output a mixture distribution that can take on many forms. This flexibility is critical for many real-world problems that may yield multi-modal distributions, but comes at the cost of being less amenable to the estimation of second-order uncertainty (at present, we do not have a way to estimate the uncertainty of the mixture distribution produced by the model).

### Relation to probabilistic models in AI

Our approach is also distinct from other types of probabilistic models in AI. Unlike models such as variational autoencoders, which learn distributions over the input stimulus itself and aim to capture the full data-generating process^98^, our model produces a distribution only over the estimated target variable. The correspondence between the model’s uncertainty and human confidence suggests that humans represent the uncertainty of the variables we estimate; it remains unclear whether and in what conditions the brain represents the full data-generating process. Another form of uncertainty representation can be found in Bayesian neural networks, which express uncertainties over model parameters^99^. Such models could quantify decision uncertainty indirectly (by running multiple forward passes that each could yield distinct outputs) but have not been shown to scale reliably to high-dimensional, real-world tasks^99^. By contrast, our method directly produces output-level uncertainty in a single pass through the model, and scales to large problems.

### Limitations

The models we described here assume point sources (in space, or in f0). Experiment 9 shows that this assumption is too impoverished to fully account for human perception, in that humans can distinguish the spatial width of a source from the uncertainty over the source’s location. The models also did not represent the distance of a source from the listener, another source property humans are known to represent^100^. The modeling framework could be straightforwardly extended to incorporate richer representations of sound sources (three-dimensional locations as well as spatial extent, for instance), with distributions that capture uncertainty over each of these dimensions. The model also can only report the location for a single source, whereas listeners have some ability to localize multiple concurrent sources^101^. It remains to be seen whether humans represent the uncertainty associated with multiple concurrent sources (as opposed to, say, just the source at a focus of attention), but if so that would require representations that bind uncertainty to particular sources^1^.

The models implemented here have no temporal dynamics. They operate on brief excerpts of audio and produce a single distribution, and are free to process the entire stimulus in parallel to generate a decision. Biological sensory systems, by contrast, receive inputs that are continuous in time, and produce responses that evolve over time. These temporal dynamics can subserve evidence accumulation, which might be a way to reduce uncertainty^13^. At present our model cannot reproduce such effects, but could be straightforwardly extended to incorporate recurrent processing for this purpose. Confidence is also sometimes correlated with reaction time, with lower confidence corresponding to longer reaction times^102,103^. A model with temporal dynamics could be used to relate reaction times to uncertainty.

We used the models presented here in a manner akin to classical ideal observer models, treating the output of their optimized solutions as approximations of the normatively correct posterior distribution for a stimulus, and using them to assess whether human confidence judgments reflect the actual uncertainty of perception. In the limit of infinite training data and successful optimization, a sufficiently expressive model architecture should converge to this ideal observer. However, because the models are empirically optimized over a finite training set, and because the model architectures cannot perfectly express all possible distributions, they likely do not reach this ideal state. We view the training data as likely to limit the model robustness. In the early stages of the project, we experimented with models trained on less data, and obtained more idiosyncratic results (for instance, localization accuracy that varied less smoothly with position). We subsequently adopted a larger training set, but the models are nonetheless almost surely undertrained relative to what would be needed to converge to a fully ideal observer for data from the training distribution. The models are also limited by the training data generation process, which is impoverished relative to the full distribution of sensory data that likely shaped human perceptual systems over evolution and development.

### Future directions

Uncertainty is relevant to many aspects of perception, and the ability to create working models that represent uncertainty should open many doors. One setting where confidence could be functionally important is in the control of attention. If uncertainty is high, an observer might benefit from exerting more attention. Models that combine computational instantiations of attention^22^ with representations of uncertainty could clarify the relationship between attention and uncertainty, and could be further extended to model attentional control. Uncertainty estimates could also aid learning, by allowing larger belief updates in settings where estimates of the environment are more certain. The models developed here could be incorporated into reinforcement learning frameworks to explore this issue.

Another setting in which uncertainty is important is in the perception of music. Music perception has long been argued to be the result of an internal model that generates predictions of upcoming events, with uncertainty creating tension that drives the listener’s interest^104,105^. Measures of uncertainty derived from generative models of music have shown some ability to predict brain responses to music^106,107^, but as in other domains, generative models trained with a cross-entropy loss may not consistently capture true uncertainty. Modifying such models to produce well calibrated output distributions could enable better models of human music perception and stronger tests of the role of uncertainty in this domain.

Uncertainty should be objectively higher when sensory input is degraded, as happens routinely following hearing loss, potentially because decreased sensory input leads to effectively higher levels of internal noise. However, it remains unclear whether confidence remains well calibrated following hearing loss (i.e., with lower confidence accompanying the increase in uncertainty). Our modeling framework could be extended to models with simulated peripheral hearing loss^108^, or cochlear implants^85^, which could allow predictions of the confidence judgments that should be expected from a system that correctly estimates uncertainty in these settings.

Although we found confidence to be related to perceptual precision, and thus to be correlated with performance, there are some situations where performance and confidence dissociate^102,109–111^. Applying our modeling framework to these settings could help clarify whether these situations can be understood in normative terms. For instance, observers might learn heuristics for summarizing the overall uncertainty of a distribution that in most settings are efficient and correct but that produce biases in some conditions (see Supplementary Fig. 3b for an example of how alternative summary measures of uncertainty can alter patterns of confidence across conditions). The model could also help clarify whether humans can be fooled by confidence “illusions”, misestimating uncertainty in some settings. More generally, the model offers the opportunity to test alternative accounts of how confidence in different settings is generated from representations of uncertainty over estimated variables.

## Methods

### Model training data – Sound localization

Model training data for sound localization was generated using methods similar to those in a previous publication^20^. The primary difference from the methods of that previous publication is that we used a larger training set as we found this led to more robust model behavior. We used a virtual acoustic room simulator^112^ to generate spatialized scenes in different rooms with the listener at random positions and angles within a room. We first generated binaural room impulse responses (BRIRs) for each room, source location, and listener position, and then used these BRIRs to render scenes (by convolving BRIRs with source waveforms, and with noise waveforms, and then adding the results to get a training example waveform).

1800 rooms were used in training and a different set of 200 rooms were used for validation. Room lengths and widths were sampled log-uniformly between 3 m and 30 m, and room heights were sampled log-uniformly between 2.2 m and 10 m. For each room, we generated BRIRs for one randomly chosen listener position and all possible source directions about that listener position. The listener’s head position was sampled uniformly within each room, with the constraints that the listener was at least 1.45 m away from all walls and no higher than 2.0 m from the floor. The listener’s head angle was then randomly sampled between 0° and 90°. Source locations were generated for two distances from the listener, in each case at every 5 degrees in azimuth (spanning 0 to 355 degrees) and every 10 degrees in elevation (spanning 0 to 60 degrees). The two source distances included a fixed distance of 1.4 m, and a second distance that was independently sampled for each room, drawn uniformly between 1.0 m and *d*, where *d* was 0.1 m less than the distance from the listener to the nearest wall. BRIRs were rendered at a sampling rate of 44.1 kHz.

Each training example was generated as a 1.5 s binaural audio clip, to allow for subsequent cropping of the cochleagram representation to 1 s (to avoid boundary artifacts). Each training example contained the foreground sound to be localized superimposed on a set of background noises. Foreground sounds were drawn from the GISE-51 data set^113^, which contains monaural waveforms of recorded environmental sounds drawn from 51 classes, sampled at 44.1 kHz. To increase stimulus diversity, 50% of the foreground sounds were bandpass-filtered using a Butterworth filter. The filter width was uniformly sampled between 2 and 4 octaves, the filter order randomly selected between 1 and 4, and the center frequency log-uniformly sampled between 160 Hz and 16 kHz. A 10 ms half-Hanning window was applied to both onset and offset. A temporal jitter of up to 500 ms was then added to the clip to ensure that onset times varied across training examples. This was achieved by zero-padding the signal by a duration uniformly sampled from between 0 and 500 ms, followed by cropping or padding the end to produce a 1s clip. The resulting signal was then zero-padded with 500 ms on each side (yielding a 2 s clip), spatialized in the target room at the target location, and finally cropped to the central 1.5 s segment.

Background sounds were sourced from a subset of AudioSet^114^ that was screened to include only stationary sounds. This screening procedure involved measuring texture statistics^115^ and removing all clips where statistics were not stable over time^116,117^. The final subset included 26,515 unique 10-second noise clips for the training dataset and 562 clips for the validation dataset. These sounds were also sampled at 44.1 kHz. Between 3 and 15 background sounds (random 1 sec crops of the same signal) from AudioSet, spatialized to randomly sampled locations within the same room, were added at an SNR drawn uniformly from [-15, +30] dB. The resulting binaural signals were RMS-normalized to 0.1. We generated a total of approximately 18M training scenes and 500K validation scenes.

To ensure balanced sampling of spatial cues across hemifields during training, the training data was augmented by mirroring each training example across the interaural axis (i.e., swapping the left and right audio channels). This produced a left–right reversed version of the original scene. The associated sound source location labels were transformed by inverting their azimuthal coordinates relative to the midline, with angular values wrapped within a 0–360° polar coordinate system. The final training set contained both the original and mirrored auditory scenes, for a total of approximately 36M training scenes.

### Deep neural network models of sound localization

#### Overview

Spatialized binaural waveforms were first processed by simulated cochlea corresponding to the left and right ears. The resulting two-channel cochlear representations were then passed to a deep convolutional neural network, which provided a non-linear transformation of the input and could be viewed as a functional instantiation of potential central auditory pathway computations. The network outputs a low-dimensional parameterization of a probability distribution over sound-source azimuth and elevation. Model parameters are optimized using stochastic gradient descent, with supervision provided by the ground-truth location of the foreground sound source in the auditory scene.

A key distinction from previous models of perception based on deep learning is that this framework supports continuous-valued outputs rather than discrete class labels, enabling readouts that are better aligned with the continuous variables that underlie many human perceptual judgments. This approach also mitigates label-dependence, in this case allowing the system to generalize to source locations not encountered during training.

Subsequent sections provide implementation details.

#### Peripheral auditory model

Binaural input waveforms were transformed into a cochlear representation intended to approximate cochlear and auditory nerve processing. Each channel (left/right) was passed through a finite-impulse-response approximation of a 40-channel gammatone filter bank, with impulse responses truncated to 25 ms for memory efficiency and characteristic frequencies spaced uniformly on an ERB-number scale^118^ between 40 Hz and 20 kHz. The resulting subbands were half-wave rectified and subjected to a compressive nonlinearity by raising amplitudes to the power of 0.3, simulating the compression from outer hair cells. The result was then lowpass filtered with a cutoff of ∼3.79 kHz to simulate the upper limit of auditory nerve phase locking and downsampled to 8 kHz. Lowpass filtering and downsampling was performed with Torchaudio’s resample method using a Kaiser-windowed sinc filter of width 64, rolloff 0.94759, and β = 14.76965. To avoid signal onset and offset artifacts, the middle 1 second was extracted from the full 1.5-second representation, producing tensors with dimensions 2 × 40 × 8000 (ear × frequency × time).

#### Neural network model

The 2 x 40 x 8000 cochlear representation was passed through a deep convolutional neural network backbone composed of hierarchically organized stages. Each stage implemented one or more standard operations, including two-dimensional convolutions, pointwise nonlinear rectifications, max pooling, batch normalization, linear transformations, and dropout regularization. For the sound localization task, we adopted a representative backbone architecture from Francl and McDermott (2022)^23^. The full specification of the architecture is provided in Supplementary Table 1.

#### Model readout

The neural network computes a latent representation *z* = *f*(*x*) from the input stimulus *x*, where *f* is parameterized by trainable convolutional kernels and biases. Instead of projecting *z* onto a deterministic point estimate of source location or a discretized spatial bin, we employ a readout function *g* that maps *z* to the parameters of a conditional distribution *p*(*y*|*x*) over the possible locations of the sound source. Depending on the ambiguity of the acoustic cues in the stimulus, the true posterior over source location may be sharply peaked, diffuse, or even multimodal; explicitly modeling a distribution over locations accommodates these forms of uncertainty. Model parameters were learned by maximizing the conditional log-likelihood, a strictly proper scoring rule^119^, thereby promoting posteriors that are not only accurate (tending to have highest probability for the true variable value) but also well-calibrated (tending to have a spread that reflects the actual uncertainty of the estimated variable).

The natural spatial coordinates for sound localization exhibit a non-Euclidean topology. To respect this geometry, we employed circular probability distributions, avoiding the boundary artifacts that would otherwise arise under Euclidean loss formulations. Concretely, we parameterized the distributional readout as a finite mixture of factorized bivariate von Mises densities. The von Mises family is well-suited for this setting, as it naturally encodes circular structure and can be described by a small number of parameters. The readout *g*(*z*) was implemented with three output heads: one producing the mixture weights, another the component means, and a third the component concentration parameters (analogous to the reciprocal of the variance parameters of normal distributions). The number of mixture components was treated as a hyperparameter; in all reported experiments, it was fixed at *K* = 5, which we found to provide sufficient expressivity. We now describe the parameterization of each head in detail.

The readout 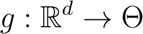 parameterizes a finite mixture of *K* independent bivariate von Mises components. For a target 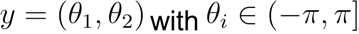, the conditional density is:

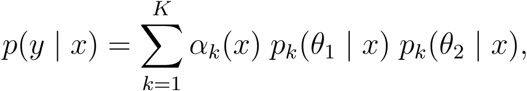

where for component *k*

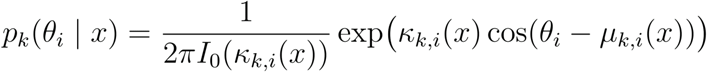

*I*_0_(·) is the modified Bessel function of the first kind (order zero). Thus, each component factorizes as follows.

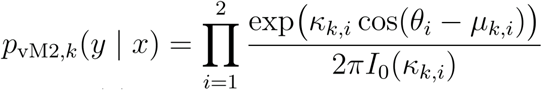

The three heads of the readout *g*(*z*) were used to predict the mixture weight (*α_k_*), distributional mean (*µ*_k_), and concentration parameter (*k*_k_) of each component *k*, respectively.

The first readout head produces logits 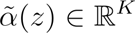, which are subsequently mapped to the simplex via the softmax function 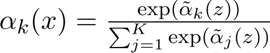 such that 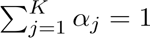. These correspond to the mixture weights.

The second readout head produced a 2D vector for each mixture component *k* and each dimension (azimuth/elevation) *i* as 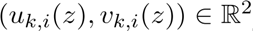. Here, we ensured the legitimacy of *u*(·), *v*(·) as sin and cosine components by passing them through the hyperbolic tangent activation function. They were then mapped to the circle via 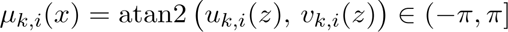.

The third readout head outputted the concentration parameters that control the spread of each mixture component. These parameters are subject to two constraints. First, they must be non-negative. For reference, a concentration of zero corresponds to a uniform density, and larger values correspond to narrower densities. Very large values yield extremely narrow densities and lead to numerical instabilities, and thus must be avoided. To enforce these constraints, we applied a modified rectifying nonlinearity that bounds the activations both below and above. In our experiments, we set the lower bound to *lb* = 1 and the upper bound to *ub* = 512. A concentration parameter of *k* = 512 yields a close approximation to a wrapped normal distribution with variance 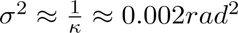, which is effectively a delta function in practice.

#### Justification and interpretation of training objective

While the location of sound sources can serve as “true” labels for the mean values of the distributional readout, no ground-truth information is available for the concentration parameters and the mixture weights. To enable the model to learn to correctly set these parameters given an input stimulus, we optimized the negative log-likelihood of the true source locations under the predicted von Mises density. The log-likelihood naturally couples the estimation of means and concentrations during training: high concentration values (resulting in low dispersion of probability, i.e., lower variance) are encouraged when predictions closely match the targets, while lower concentration values (resulting in higher dispersion, i.e., higher variance) are favored when uncertainty is high. This formulation allows the model to learn both accurate and appropriately calibrated distributions without requiring explicit supervision of the concentration parameters.

The proof of this general idea is straightforward, and we provide it below:

In the following proof, we will use *x* to denote data (auditory signals) and *y* to denote labels (spatial locations or fundamental frequency). Let us suppose that there is a true conditional distribution *p*^∗^(*y*|*x*) that specifies the relationship between data and labels and that our model predicts a conditional density *q*_4_(*y*|*x*), where *θ* refers to the model parameters. We show below that minimizing the expected negative log-likelihood of the true labels under the predicted distribution recovers the true conditional distribution.

#### Proof that the log-likelihood objective will cause the model to predict the true posterior

The expected negative log-likelihood is defined as follows:

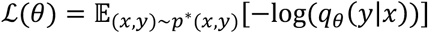

By expanding the expectation, we get:

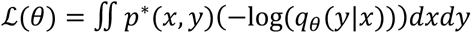

We can use the factorization *p*^∗^(*x*, *y*) = *p*^∗^(*x*)*p*^∗^(*y*|*x*) to obtain

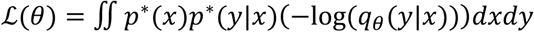

Rearranging the integrals,

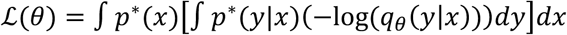

Adding and subtracting log(*p*^∗^(*y*|*x*)) to the term in the inner integral:

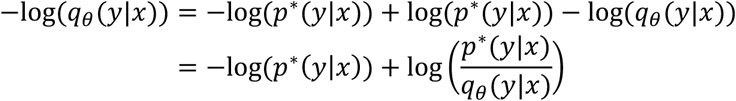

Substituting back into the loss formulation, we get the following:

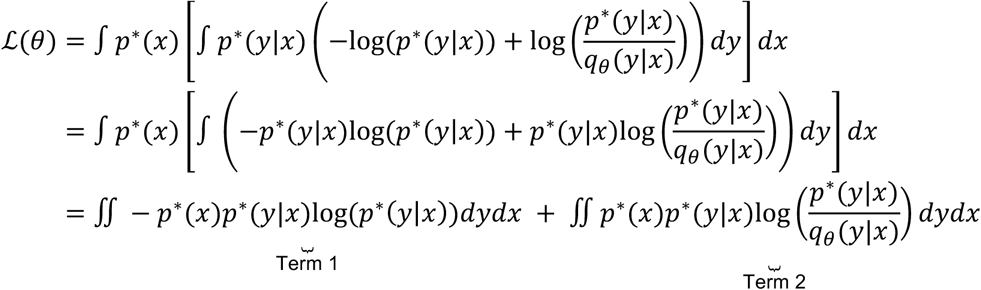

The first term is the conditional entropy *H_p_*_∗_(*Y*|*X*), which depends only on the true data-generating distribution *p*^∗^. Since it does not involve the model parameters *θ*, it is a constant with respect to optimization.

Justification: The entropy of a random variable *Z* is defined as:

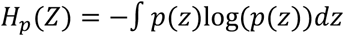

The conditional entropy of the labels *Y* given a particular value of a signal *x* is thus

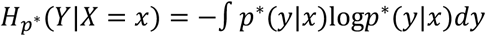

To get the overall conditional entropy, we have to consider all possible *x*:

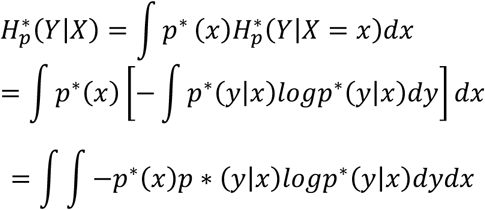

The second term is the expected value of the Kullback-Leibler (KL) divergence between the true conditional distribution *p*^∗^(*y*|*x*) and the model’s predicted distribution *q_θ_*(*y*|*x*). This term measures how different the model’s predictions are from the true conditional distribution.

Justification: the KL divergence is defined as:

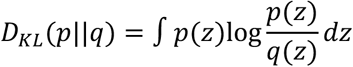

Applying this definition to our “true” and predicted distributions for a specific value of *x* we get the following:

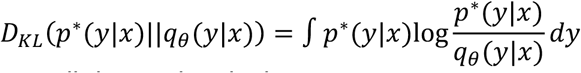

The expectation of this over all data points is thus

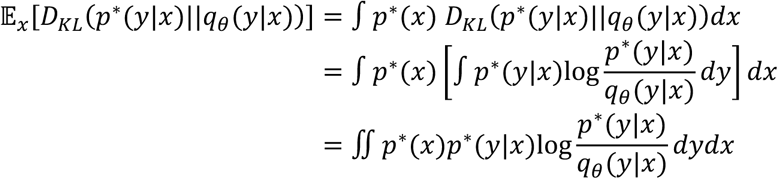

Therefore,

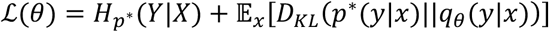

Because the first term is fixed and cannot be reduced by changing the model, minimizing ℒ(*θ*) is equivalent to minimizing the expected KL divergence between the true and predicted conditional distributions. Since the KL divergence is always nonnegative (*D_KL_*(*p*||*q*) ≥ 0), with equality if and only if *p* ≡ *q* almost everywhere, it follows that

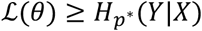

The lower bound is achieved precisely when *p*^∗^(*y*|*x*) = *q_θ_*(*y*|*x*). In other words, the minimum possible loss is attained when the model recovers the true conditional distribution.

In practice, achieving this optimum requires several idealized assumptions: (i) infinitely many training samples so that the empirical risk minimization approximates minimization of the population loss; (ii) a model class rich enough to represent the true conditional distribution *p*^∗^(*y*|*x*); and (iii) successful optimization that finds a global minimizer of the objective. Under these conditions, the learned distribution converges to the true conditional distribution, and the loss approaches the lower bound *H_p_*_∗_(*Y*|*X*) that is irreducible uncertainty.

#### Training objective

Given a dataset 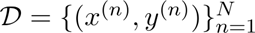, the conditional log-likelihood is defined as follows:

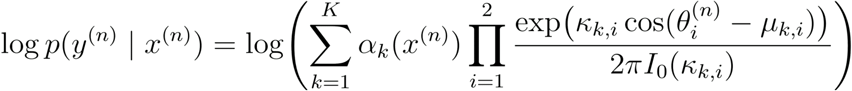

For numerical stability, we employ the log-sum-exp technique used in machine learning^120^. Parameters are learned by maximizing the total conditional log-likelihood, or equivalently, minimizing the negative log-likelihood, which is a strictly proper scoring rule that encourages calibrated posterior distributions.

We implement the logarithm of the modified Bessel function of the first kind, order zero, using piecewise polynomial approximations. For small arguments *k* = 3.75, we apply a polynomial expansion of *I*_0_ (*k*) and then take the logarithm. For large arguments, we use the asymptotic expansion 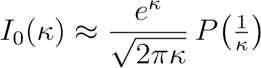, where *P*(1/*k*) is represented as a polynomial in 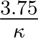. This implementation ensures numerical stability and differentiability across the full input domain. Polynomial coefficients were taken from a textbook^121^.

#### Training details

The model was trained using the Adam optimizer with an initial learning rate of *lr* = 0.001. We employed a warmup and decay learning rate scheduler, allocating 20% of the total training steps to a linear warmup phase and 20% to a decay phase, with a multiplicative decay factor of γ = 0.9. The nominal batch size was set to 64. However, due to batch-wise symmetric augmentation, the effective batch size was doubled to 128Training proceeded for a maximum of 5 epochs, with early termination if the training loss exhibited saturation. Training was performed on an NVIDIA A100 GPU processor. We trained 5 model instantiations, each with a different random seed.

#### Alternative model trained with cross-entropy loss

The training dataset and all training hyperparameter details were the same as for the uncertainty-aware model. The model architecture was also the same except that the final linear layer’s output was 504-dimensional. Each output unit corresponded to a location bin (5×5 degrees in azimuth and elevation, as in previous classification-based localization models^20,23^). The model was trained with a cross entropy loss. Training was performed on an NVIDIA A100 GPU processor and proceeded for a maximum of 5 epochs, with early termination if the training loss exhibited saturation. We trained 5 model instantiations, each with a different random seed. Model results were averaged across the 5 model instantiations.

### Human and model psychophysics – sound localization

#### Overview

All in-house human sound localization experiments were conducted using a custom-built loudspeaker array comprising 133 loudspeakers (KRK Classic 5), arranged to cover azimuths from –90° to +90° in 10° increments and elevations from –20° to 40° in 10° increments. The speakers had a flat frequency response between 46Hz and 30 kHz. Participants recorded their responses using an iPad interface that displayed a schematic of the loudspeaker array, with each loudspeaker position represented as a selectable button. Additional interface options were presented on the screen depending on the specific experimental requirements.

#### Model experimental stimuli

We simulated each experiment on the model using a virtual room designed to closely replicate the physical loudspeaker array room, based on measurements of the real space. The simulated room measured 4.66 m × 5.9 m × 2.48 m. Three of the walls were modeled as 1-inch-thick fiberglass acoustic treatments, with the frontmost wall modeled as painted concrete. Materials within the simulator were chosen to approximate the acoustic properties of the actual room. Binaural room impulse responses (BRIRs) were generated for 133 loudspeaker locations, each positioned 2 meters from the listener, matching the physical array configuration.

Each sound stimulus was convolved with the corresponding BRIR to create a virtual version of the set of trials presented to human participants. To minimize boundary artifacts, all sounds were zero-padded to 1.5 seconds prior to spatialization and then truncated to 1 second afterward, with the sound centered in the signal. Audio processing was performed at a sampling rate of 44.1 kHz. After spatialization, the sounds were upsampled to 48 kHz to match the input rate required by the models.

To approximate the ambient noise of the testing chamber (which was not acoustically isolated), we spatialized 10–20 randomly selected audio textures from the AudioSet dataset and placed them at random locations within the virtual room. Spatialized background noise was added at an SNR in [25, 35] dB, consistent with the measured noise levels in the physical array room (between 30 and 40 dB SPL, with stimuli presented at 65 dB SPL).

#### Model localization judgments

To obtain a location “response” from the model for a given stimulus (Experiments 1-3, 5, 6 & 8), we computed a maximum a posteriori (MAP) estimate from the stimulus-conditioned posterior as predicted by the model. Due to the mixture parameterization of the posterior, the MAP estimate does not admit a closed-form solution. Instead, we approximated the MAP estimate via Monte Carlo sampling. For each stimulus-specific posterior, we drew 5000 samples and constructed an empirical histogram with a bin width of 1 degree. The mode of this empirical distribution was then taken as the MAP estimate and used as the model’s location response.

For Experiment 4, which used binary front-or-back judgments, we first obtained a location estimate as for absolute localization experiments, and then determined whether this location was in the front or back hemifield.

#### Model confidence judgments

To obtain model confidence judgments, we chose measures that summarized the spread of the distribution over the presumptive decision variable for a task. For experiments with absolute localization judgments (Experiments 1-3, 5, 6 & 8), the posterior estimated by the model is the distribution over the decision variable, and so a measure of posterior spread was appropriate as a summary measure. As a robust, non-parametric measure of posterior spread, we chose the width of the 50% highest posterior density (HPD) interval of the empirical posterior obtained from Monte Carlo sampling (this is the narrowest interval containing 50% of the probability mass under the density). The 50% HPD is sensitive to local structure in the distribution while avoiding undue influence from low-probability tails. To convert these continuous uncertainty values into discrete “bets” akin to those made by humans, we mapped HPD widths to a five-point scale (1–5) using an inverse cumulative distribution function (CDF) procedure. HPD widths (for each experiment with its own set of trials) were sorted in descending order and assigned to bet values according to a Gaussian prior over the betting scale. Specifically, the mapping distribution was defined as a discrete normal distribution over the integers 1–5, centered at 3 with unit variance, and normalized to sum to one. The cumulative probabilities of this Gaussian prior were used to map the sorted HPD widths to discrete bets.

We make no claim that the 50% HPD is the only or best way to summarize the posterior for this purpose, and found that similar model results were obtained with alternative measures (Supplementary Fig. 3a).

For experiment 4, which used a binary front-or-back judgment, the posterior (again estimated from Monte Carlo sampling) was instead collapsed to two bins (integrating the posterior over all front locations, and all back locations), from which we computed entropy. We note that different patterns of results could be obtained from summary measures that summarized the full posterior rather than the distribution over the presumptive decision variable (Supplementary Fig. 3b).

### Localization model validation – Psychoacoustic experiments

We replicated the battery of psychoacoustic experiments used by Francl and McDermott (2022)^23^ to evaluate models of sound localization, employing the stimuli and analysis procedures as those used by Saddler and McDermott (2024)^20^. Here, we provide a brief overview of each of these experiments.

#### Macpherson and Middlebrooks (2002) – Duplex theory

In the original experiment^45^, the authors tested the relative contributions of interaural time (ITDs) and level differences (ILDs) to perceived azimuth for low-pass and high-pass filtered noise. The influence of imposed ITD or ILD biases on localization judgments was quantified with a perceptual weight (the slope relating the imposed ITD or ILD to the shift in the perceived location, with the shift expressed in terms of the change in ITD or ILD that would normally accompany a location shift of that magnitude). The original experiment found that humans and exhibited high perceptual weight for ITDs and low perceptual weight for ILDs with low-pass noise, with the opposite pattern for high-pass noise. These results provide support for the duplex theory, which posits that ITD and ILD cues are differentially weighted depending on the frequency range. The models reproduced the pattern of results seen in humans.

#### Yost and Zhong (2014) – Bandwidth dependence

In the original experiment^46^, the authors measured the effect of stimulus bandwidth on localization accuracy. Stimuli were presented every 15° azimuth at center frequencies of 250, 2000, and 4000 Hz, with bandwidths of 1/20, 1/10, 1/6, 1/3, 1, and 2 octaves. Humans exhibited a decrease in localization error as the bandwidth increased. The models reproduced this pattern of results.

#### Hebrank and Wright (1974) – Spectral cues to elevation

In the original experiment^47^, the authors measured human elevation judgments as a function of the frequency content of sound sources. Sounds were presented every 30° in elevation on the midline and were either high-pass or low-pass filtered noise bursts with cutoff frequencies ranging from 3.8 to 16 kHz. Human accuracy varied systematically with frequency content, being poor for sounds lacking high-frequency content. This result provides support for the importance of high-frequency cues to elevation. The model replicated the pattern of results seen in humans.

#### Hofman et al. (1998) – Ear alteration

In the original experiment^49^, the authors measured the effect of altering pinna shape on localization accuracy. White noise bursts were presented every 20° in both azimuth and elevation, and localization performance was assessed before and after the pinna shape was modified. Humans exhibited a reduction in elevation localization accuracy following the change in pinna shape, underscoring the importance of ear-specific pinna cues to elevation. The model replicated the pattern of results seen in humans.

#### Litovsky and Godar (1999) – Precedence effect

The precedence effect^50,52,122^ occurs when identical signals arrive from different locations with a short delay. If the delay is sufficiently short, humans typically hear a single sound that appears to come from the location of the initial sound. We simulated an experiment originally run in humans^51^. In this experiment, two identical pink noise bursts were played from loudspeakers. The leading noise burst came from one of six locations (±20°, ±40° or ±60°) and the lagging noise burst came from 0°. The lagging noise burst was delayed relative to the leading noise burst by 5, 10, 25, 50 or 100ms. For each pair of noise bursts, participants reported the apparent location of each perceived sound. The experimenters then calculated the mean localization error separately for the leading and lagging noise burst for each time delay. Humans perceived a single source at the leading location for small delays, as reflected by high azimuth error for the lagging sound under these conditions. The model replicated this pattern of results.

#### Kulkarni and Colburn (1998) – HRTF smoothing

In the original experiment^48^, the authors tested the spectral resolution of spectral localization cues by presenting humans and models with broadband noise bursts processed through HRTFs with varying levels of spectral smoothing. Human participants discriminated the processed sounds from sounds presented from loudspeakers. Humans were unable to distinguish the processed sounds from free-field presentation until the HRTFs were substantially smoothed, indicating that localization relies on relatively coarse spectral cues. To assess this effect in models we measured localization error as a function of HRTF smoothing. The models exhibited a similar dependence of accuracy on smoothing to that seen in humans.

We generated model bets for the four experiments that were analyzed in terms of absolute localization judgments of single sources. The procedure for obtaining bets from the model posterior was the same as that used for the other absolute localization experiments in this paper (e.g. Experiment 3), described above.

### Experiment 1: Localization of natural sounds in quiet

#### Human experiment

Participants localized sounds presented from an array of loudspeakers. Sounds were drawn from a set of 160 unique recorded natural sounds sampled at 44.1 kHz, each cropped to 1 second in duration, with 10 ms Hanning windows applied to the onset and offset. Sounds were presented at 65 dB SPL. Each participant heard each of the 160 sounds 7 times over the course of the experiment, each time presented from a different location.

Participants sat facing the loudspeaker array and began each trial with their head oriented toward the center of the array (0°, 0°). When ready, they initiated the trial by pressing a button, triggering the sound presentation. At the end of each trial, participants responded by selecting the label of the loudspeaker they believed produced the sound. During the response period, participants were permitted to turn their heads to identify the sound source location. Each participant completed 1,120 trials (160 sounds x 7 locations). Locations were pseudo-randomly assigned to sounds such that across the 18 participants, every sound was presented from every loudspeaker location at least once.

19 participants (9 female; ages 19-28) completed the experiment. Data for 18 of the 19 participants was collected by A. Francl and R.P. Hess, and originally included in Francl’s PhD thesis.

#### Model experiment

We simulated the experiment following the procedure described above, using the same set of 160 natural sounds presented to human participants. To maximize the reliability of the model results, the complete set of sounds and locations was generated 16 times, using background noise clips and SNR values that were randomly sampled for each repeat.

#### Analysis

For each of the 160 unique sounds used in the experiment, we computed the mean absolute azimuthal error and mean absolute elevation error for both humans and models. Human-model similarity was assessed by computing the Pearson correlation between these two measures, independently for azimuth and elevation. Raw correlations were corrected for attenuation^123^ using the reliabilities of each error measure (estimated as split-half correlations, corrected for sample size using the Spearman-Brown formula^124^).

### Experiment 2: Localization of natural sounds in noise

The human data for this experiment were collected and published in a previous paper^20^. The following methods description of the experimental paradigm is reproduced from that paper with minor edits.

#### Human experiment

On each trial a target natural sound was played from one of 95 loudspeakers while threshold-equalizing noise^125^ was played concurrently from 9 other loudspeakers. Target and noise locations were randomly sampled on each trial. The listener’s task was to report which loudspeaker produced the target by entering the loudspeaker’s label on a keypad. Listeners were instructed to direct their head at the loudspeaker directly in front of them for the duration of the stimulus. Once the stimulus ended, they could look at the loudspeaker where they thought the target had occurred to obtain the label. Target sounds were drawn from a set of 460 recorded natural sounds from the GISE-51^113^ evaluation dataset. Each target sound was presented once over the course of the experiment. Target sounds were presented at 60 dB SPL (A-weighted) and noise levels were determined such that the SNR of the target relative to the sum of the 9 noise sources was −13.6, −6.8, 0, +6.8, or +13.6 dB. All stimuli were sampled at 44.1 kHz and were 1 s in duration, including 15 ms onset and offset ramps (Hanning window). 11 normal-hearing listeners (5 female) with ages between 21 and 30 years each performed 460 trials.

#### Model experiment

Models were tested on all combinations of the 460 target natural sounds, 5 SNRs, and 95 target locations (218,500 total stimuli) used in the human experiments, replicating the model experiment in a previous publication^20^. Sources were spatialized in a virtual rendering of the loudspeaker array room human listeners were evaluated in. To match the task between human and models, we restricted model localization judgments to azimuth and elevations corresponding to the 95 loudspeaker locations.

#### Analysis

Human and model performance was quantified by measuring mean absolute spherical error (great circle distance).

### Experiment 3: Effect of stimulus bandwidth on localization and uncertainty judgments

#### Human experiment

Stimuli were designed to vary in localization accuracy. They included three types of sounds: broadband noise, pure tones (600, 2000 and 4000 Hz), and narrowband noise centered at the same frequencies as the pure tones (with one-octave bandwidths). Each sound was 750 ms in duration, modulated by linear on and off ramps (250 ms in duration). All sounds were sampled at 44.1 kHz and presented at 65 dBA. On each trial we jittered the center frequencies of tones and narrowband sounds within a half-octave range. For broadband sounds, we randomly varied the lower (uniformly at random between 20 and 60 Hz) and upper (uniformly at random between 8 and 16 kHz) cutoff frequencies.

Participants sat facing the loudspeaker array with their head oriented toward the center of the array (0,0) and ears aligned with −90 and 90 degrees azimuth. On each trial, participants were first cued with the elevation from which the sound would be presented. When ready, they initiated the trial by pressing a button, triggering the stimulus presentation. Participants were instructed to judge which loudspeaker produced the sound, with response options restricted to the speakers at the elevation specified at the beginning of the trial. After selecting a loudspeaker, participants placed a bet between 1 and 5 cents (integer values) on their response. Bonus payments were determined trial by trial, with the bet magnitude awarded for correct responses and subtracted for incorrect responses, and with total negative earnings floored at zero. For the purpose of calculating payments, a response was considered correct if the selected loudspeaker was within one position of the true source. During sound presentation, participants were instructed to maintain fixation on the loudspeaker located at 0° azimuth, 0° elevation (and to not turn their head). Stimuli were presented from azimuths spanning −90° to +90° in 10° steps. On each trial, the elevation was randomly assigned to be between 0° and 40° (in integer steps of 10°). Each participant completed a total of 399 judgments (7 stimulus conditions × 19 azimuths × 3 repetitions). No intermediate feedback was provided to the participants.

16 participants completed the experiment (8 female; mean age = 22.4 years, SD = 5.72 years).

#### Model experiment

We simulated the experiment in a virtual replica of the speaker array room, using the procedure described above and employing the same stimuli presented to human participants. To maximize the reliability of the model results, the complete set of sounds and locations was regenerated 128 times, with background noise clips and SNR values randomly sampled for each repeat.

#### Analysis

For both human participants and models, we quantified four metrics as a function of stimulus azimuth: localization accuracy, response bias, response precision, and response confidence. To calculate each metric, we first selected the subset of trials corresponding to a given target azimuth and stimulus type. From this subset, we computed participant- or model-level statistics, which were then aggregated across participants or models to obtain group-level values. Error bars indicate the standard error of the mean.

Localization accuracy was quantified as the absolute azimuthal error. For humans, we computed the median absolute error for each participant, then averaged these medians across participants. The median was used because it is more robust given the small number of trials per condition per azimuth. For models (which performed more trials than humans), we computed the mean absolute error per model and then averaged across models.

Response bias was captured by the mean reported azimuth for each azimuthal position. For each participant (or model), we computed the mean reported azimuth across trials, then averaged these means across participants (or models).

Response precision was quantified using the mean absolute deviation (MAD) of reported azimuths. For humans, MAD was calculated from data pooled across all participants to increase robustness, reflecting the assumption that participants perform the task similarly. Confidence intervals on the human MAD were obtained via bootstrapping across participants. For models, MAD was computed separately for each model and then averaged across models, and error bars were obtained via the standard error across models.

Response confidence was quantified as the average bet across the trials for a condition for each participant (or model), then averaged across participants (or models).

### Experiment 4: Front-back estimation and uncertainty judgments of controlled stimuli

#### Human experiment

Participants listened to sounds presented through a semicircular loudspeaker array. Half of the participants were seated with the loudspeaker array to their right; the remaining participants sat with the array to their left. In each case the participant sat facing the outermost speaker in the array, such that the speakers spanned 0 to 180 degrees to either the left or right of the participant. We included two types of stimuli: broadband noise and pure tones (600 Hz, 2000 Hz, and 4000 Hz). Each sound was 750 ms in duration, modulated by linear on and off ramps (250 ms in duration). All sounds were sampled at 44.1 kHz and presented at 65 dBA.

Stimuli were presented at all azimuths ranging from 0° through 180° in 10 degree steps (relative to the participants’ nose), omitting 90° (which lacks an unambiguous front–back label), and elevations 0° to 40° (10 degree steps) in a fully crossed experimental design. On each trial, participants were asked to report whether the sound originated from the front or back hemifield. Participants then placed a bet between 1 and 5 cents (integer values) on their response. Bonus payments were determined trial by trial, with the bet magnitude awarded for correct responses and subtracted for incorrect responses, with total negative earnings floored at zero. No feedback was provided to participants during the experiment. Each participant completed a total of 360 trials (4 stimulus conditions × 18 azimuths × 5 elevations).

10 participants completed the experiment (5 female; mean age = 26.4 years, SD = 6.7 years).

#### Model experiment

We used the same stimuli as in the human experiment, but zero-padded each sound to 1.5 seconds before spatialization, centered the stimuli within this window, and cropped stimuli to 800 ms. Spatialization followed the same procedure described above, but we generated a new virtual room configured to match the participants’ orientations during the experiment. The experiment was repeated 128 times, randomly varying background noise clips and SNR values across repetitions.

To remain consistent with the idea that confidence reflects decision uncertainty, we computed bets for this experiment by integrating the model posterior over front and back locations to obtain a binomial distribution. The model’s confidence was defined as the entropy of this distribution, which was then mapped to bets ranging from 1–5 cents using the same non-linear monotonic transformation employed in the other experiments.

#### Analysis

For both human participants and models, we quantified two metrics as a function of stimulus azimuth: front-back discrimination performance and response confidence. We report them as a function of azimuthal position.

Discrimination performance was assessed using sensitivity (d’) calculated for azimuthal pairs of location on the same cone of confusion (e.g., 0 and 180 degrees, 10 and 170 degrees, 20 and 160 degrees etc.). Hits were defined as “front” responses on trials where the sound came from the front location, and false alarms were defined as “front” responses on trials where the sound came from the back location. From the corresponding hit and false alarm rates, we computed a d’ value for each participant (or model), which was then averaged across participants (or models). Error bars indicate the standard error of the mean (s.e.m.). Response confidence was quantified by averaging bets separately for front and back trials for each location pair, then taking the mean of these two values. As with sensitivity, this measure was computed per participant (or model) and then averaged across participants (or models), with s.e.m. as the error bars.

### Experiment 5: Model-guided natural sound localization experiment

#### Model experiment

Stimuli were selected using the model’s uncertainty. We evaluated the model on a set of 160 unique natural sound stimuli that were not used during training (the same sounds used in Experiment 1), rendered in a virtual replica of the speaker array room as in other experiments. The goal was to identify subsets of stimuli with high and low uncertainty.

For each stimulus, we computed a mean uncertainty value by presenting it to the model at every azimuth (–90° to 90° in 10° steps) and elevation (0° to 40° in 10° steps) with each pair repeated 16 times with different background noise samples. At each azimuthal location, uncertainty was quantified as the width of the 50% highest posterior density (HPD) interval of the empirical posterior along the azimuthal axis. For each model, we averaged these uncertainty values across the different elevation presentations, then averaged across models. The resulting stimulus-level means were sorted, and we selected the 10 stimuli centered on the 5th and 95th percentiles, respectively. We drew stimuli from these ranges to avoid potential floor and ceiling effects in both model performance and bets (as might have occurred with the very top and bottom sounds).

The selected stimuli were as follows:

*Low uncertainty group*: Stapler, Fingernail tapping, Basketball bouncing, Chop vegetables, Printing, Walking on hard surface, Scissors cutting paper, Pouring water, Newspaper rustling, and Doorbell.

*High uncertainty group*: Siren, Keyboard synthetic, Mallet, Reed acoustic, Keyboard electronic, Whistling, Busy signal, Humming, and Organ.

#### Human experiment

We subsequently tested human participants in an experiment similar in design to Experiment 3, with the main difference that each participant was presented with all 20 selected stimuli at every azimuth (in randomized order). On each trial, the elevation was randomized between 0° and 40°, and participants were informed of the elevation in advance. Responses were restricted to azimuthal positions at the presented elevation. On each trial, participants provided both an azimuthal localization response and a confidence bet (1–5 cents). Sounds were presented at 65 dBA.

15 participants completed the experiment (11 female; mean age = 21.6 years, SD = 3.5 years).

#### Analysis

To evaluate human performance and its correspondence to model predictions, we calculated three metrics as a function of stimulus azimuth: localization accuracy, response precision, and response confidence. For each metric, we first selected the subset of trials corresponding to a given target azimuth and stimulus group (high vs. low uncertainty). Participant-or model-level statistics were then computed from these subsets and aggregated across participants or models to obtain group-level values. Error bars represent the standard error of the mean (s.e.m.).

As in Experiment 3, localization accuracy was quantified as the absolute azimuth error, and response precision as the mean absolute deviation (MAD) of reported azimuths. For human data, MAD was pooled across all participants and confidence intervals were obtained via bootstrapping. For models, MAD was computed separately for each model and then averaged across models, and error bars were computed as the standard error across models. Response confidence was measured as the mean bet across the selected trials for each participant (or model), followed by averaging across participants (or models).

### Experiment 6: Second natural sound localization experiment

#### Model experiment

Stimuli were again selected using the model’s uncertainty, using the same set of 160 natural sound stimuli from which the Experiment 5 stimuli were selected (the same sounds used in Experiment 1, and that were not used during training), rendered in a virtual replica of the speaker array room as in other experiments. The goal was to sample stimuli spanning the full range of model uncertainty. The only difference from the Experiment 5 selection procedure was that 20 stimuli were selected by uniformly sampling the distribution between its 5th and 95th percentiles.

The selected stimuli were as follows: Walking on hard surface, Car engine, Coin in vending machine, Coin dropping, Glass smash, Food chopping, Tearing, Rustle, Coughing, Electric bass, Wind chimes, Geese, Vacuum, Duck quack, Zipper, Leaf blower, Telephone ringing, Dial tone, Train warning bell, Keyboard electronic.

#### Human experiment

The experimental procedure was exactly the same as that of Experiment 5. 27 participants completed the experiment (17 female; mean age = 21.8 years, SD = 3.4 years).

#### Analysis

To evaluate human performance and its correspondence to model predictions, we calculated item-wise localization accuracy and response confidence. For each stimulus, participant- or model-level statistics were computed from the corresponding subset of trials and then aggregated across participants or models to obtain group-level values. Error bars represent the standard error of the mean (s.e.m.).

As in Experiments 3 and 5, localization accuracy was quantified as the absolute azimuth error and response confidence as the mean bet.

To assess human–model correspondence, Pearson correlations across stimuli were computed between model and human localization errors and between model and human confidence ratings.

### Model training data – Pitch estimation

Training and validation datasets used to train our deep neural network models of pitch estimation were adopted from a recent publication from our lab^24^. The following methods description of the dataset generation process is reproduced from that paper with minor formatting edits.

#### Overview

The main training set consisted of 50-ms excerpts of speech and musical instruments. This duration was chosen to enable accurate pitch perception in human listeners^126^, but to be short enough that the F0 would be relatively stable even in natural sounds such as speech that have time-varying F0s. The F0 label for a training example was estimated from a “clean” speech or music excerpt. These excerpts were then superimposed on natural background noise. Overall stimulus presentation levels were drawn uniformly from a 60 dB range, up to a maximum possible root-mean-squared amplitude of 0.632. All training stimuli were sampled at 32 kHz.

#### Speech and music training excerpts

We used STRAIGHT^127^ to compute time-varying F0 and periodicity traces for sounds in several large corpora of recorded speech and instrumental music: Spoken Wikipedia Corpora (SWC)^128^, Wall Street Journal (WSJ), CMU Kids Corpus, CSLU Kids Speech, NSynth^129^, and RWC Music Database. STRAIGHT provides accurate estimates of the F0 provided the background noise is low, as it was in each of the corpora. Musical instrument recordings were notes from the chromatic scale, and thus were spaced roughly in semitones. To ensure that sounds would span a continuous range of F0s, we randomly pitch-shifted each instrumental music recording by a small amount (up to ±3% F0, via resampling).

Source libraries were constructed for each corpus by extracting all highly periodic (time-averaged periodicity level > 0.8) and non-overlapping 50ms segments from each recording. We then generated our natural sounds training dataset by sampling segments with replacement from these source libraries to uniformly populate 700 log-spaced F0 bins between 80 Hz and 1000 Hz (bin width = 1/16 semitones = 0.36% F0). Segments were assigned to bins according to their time-averaged F0. The resulting training dataset consisted of 3000 exemplars per F0 bin for a total of 2.1 million exemplars. The relative contribution of each corpus to the final dataset was constrained both by the number of segments per F0 bin available in each source library (the higher the F0, the harder it is to find speech clips) and the goal of using audio from many different speakers, instruments, and corpora. The composition we settled on is:

*F0 bins between 80 Hz and 320 Hz*

● 50% instrumental music (1000 NSynth and 500 RWC clips per bin)
● 50% adult speech (1000 SWC and 500 WSJ clips per bin)

*F0 bins between 320 Hz and 450 Hz*

● 50% instrumental music (1000 NSynth and 500 RWC clips per bin)
● 50% child speech (750 CSLU and 750 CMU clips per bin)

*F0 bins between 450 Hz and 1000 Hz*

● 100% instrumental music (2500 NSynth and 500 RWC clips per bin)

#### Background noise for training data

To make the F0 estimation task more difficult and to simulate naturalistic listening conditions, each speech or instrument excerpt in the training dataset was embedded in natural background noise. The signal-to-noise ratio for each training example was drawn uniformly between −10 dB and +10 dB. Noise source clips were taken from a subset of the AudioSet corpus^114^, screened to remove nonstationary sounds (e.g., speech or music). The screening procedure involved measuring auditory texture statistics (envelope means, correlations, and modulation power in and across cochlear frequency channels)^115^ from all recordings, and discarding segments over which these statistics were not stable in time, as in previous studies^116,117^. To ensure the F0 estimation task remained well defined for the noisy stimuli, background noise clips were also screened for periodicity by computing their autocorrelation functions. Noise clips with peaks greater than 0.8 at lags greater than 1 ms in their normalized autocorrelation function were excluded.

### Deep neural network models of pitch perception

#### Overview

50 ms monaural waveforms were processed by a simulated cochlea, then passed to a deep convolutional network that could be viewed as a functional instantiation of potential central auditory computations related to pitch. The network produced a low-dimensional parameterization of a probability distribution over the fundamental frequency axis, optimized via stochastic gradient descent with supervision from ground-truth F0 values.

#### Peripheral auditory model

The same transformation sequence described for the sound localization pipeline was also applied to monaural inputs, but with parameters adjusted to match a previous model of pitch^24^. Waveforms were filtered with a finite-impulse-response approximation of a 100-channel gammatone filter bank, with characteristic frequencies uniformly spaced on an ERB-number scale between 60 Hz and 16 kHz. The resulting subbands were half-wave rectified and lowpass filtered with a cutoff of 3 kHz to simulate the upper limit of auditory nerve phase locking, compressed with the same 0.3 power-law nonlinearity, and then downsampled to 20 kHz. For boundary handling, the central 50 ms were taken from the full 150 ms inputs, yielding tensors of size 100 × 1000 (frequency × time).

#### Central auditory model

The 100 x 1000 cochlear representation was passed through a deep convolutional neural network backbone composed of hierarchically organized stages. As with the localization models, each stage here was implemented using one or more standard operations, including two-dimensional convolutions, pointwise nonlinear rectifications, max pooling, batch normalization, linear transformations, and dropout regularization. For the pitch estimation task, we adopted a representative backbone architecture from Saddler et al. (2021)^24^. The full specification of the architecture is provided in Supplementary Table 3.

#### Model readout

The underlying spirit of the model readout here is similar to that of the localization model, but with a distributional form that was suited to the specific characteristics of the F0 estimation problem. F0 is a one-dimensional, continuous, real-valued variable. Unlike unconstrained regression problems, however, there is no need to represent F0 values that fall outside the perceptual limits of human hearing. At the same time, the distribution over possible F0 values, given a natural sound input, is likely to be multi-modal due to well-known ambiguities such as octave confusions. These ambiguities motivate a distributional form with enough flexibility to capture complex, structured uncertainty. Taking these considerations into account, we designed the readout to produce parameters of a Gaussian mixture distribution defined over the log-F0 axis.

Let *s* = log *f*_0_ (*f*_0_ > 0), and let the readout parameterize a finite mixture of *K* independent univariate Gaussian components over *s*. For a given input *x*, the conditional density of *s* is

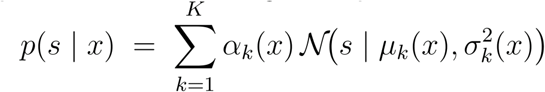

where

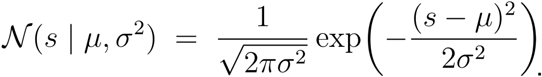

As with the localization model, the pitch model had three readout heads: one for the mixture weights, one for the component means, and one for the component variances, each with custom constraints.

The mixture weights were constrained by 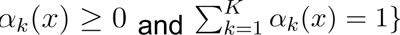. We enforced this constraint by applying the softmax function to the readout head that produced the mixture weights.

To constrain the component means to lie within the perceptually relevant range of log *F*_0_, we use a modified ReLU activation with both lower and upper bounds, which we denote as “relux”. Specifically,

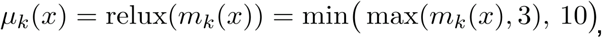

Where *m_k_*(*x*) is an unconstrained linear output of the network. The lower bound of three and upper bound of 10 correspond to the approximate log-frequency range of human hearing.

To ensure strictly positive variances, we applied a softplus transformation with a small epsilon correction:

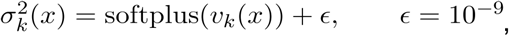

where *v_k_*(*x*) is an unconstrained linear output of the network. The epsilon correction prevents degenerate variances and avoids invalid logarithms during likelihood evaluation.

#### Training objective

We optimized the negative log-likelihood of the true F0 value under the predicted Gaussian mixture density, which naturally couples the estimation of means and concentrations. Given a dataset 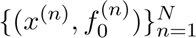, the negative log-likelihood (NLL) objective is

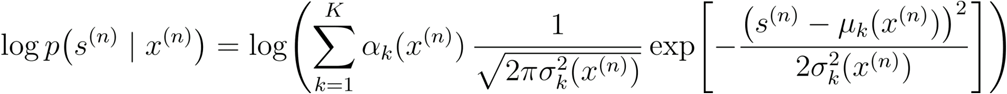

where 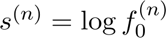,
and where the mixture component weights, component means, and component variances are model outputs. For numerical stability, we employ the log-sum-exp trick commonly used in machine learning^120^. The model parameters were then optimized by minimizing the total conditional negative log-likelihood across all training examples.

#### Training details

The model was trained using the Adam optimizer with an initial learning rate of *lr* = 0.001. We employed a warmup and decay learning rate scheduler, allocating 10% of the total training steps to a linear warmup phase and 20% to a decay phase, with a multiplicative decay factor of γ = 0.99. We used a batch size of 128 for our experiments. Training proceeded for a maximum of 25 epochs, with early termination if the training loss exhibited saturation. Training was performed on an NVIDIA A100 GPU processor. We trained 5 model repeats, each with a different random seed.

#### Pitch model validation – Psychoacoustic tests

We evaluated our models using the suite of pitch psychoacoustic experiments described in Saddler et al. (2021)^24^, employing the same stimuli and analysis procedures to provide qualitative validation. These experiments span several classic paradigms in pitch perception, including the effect of harmonic number and phase on pitch discrimination^64^, the pitch of alternating-phase harmonic complexes^65^, the pitch of frequency-shifted complexes^66^ and complexes with individually mistuned harmonics^68^, and frequency discrimination with pure and transposed tones^67^. For detailed descriptions of these experiments and their corresponding stimulus manipulations, we refer the reader to Saddler et al. (2021)^24^. Here, we provide a brief overview of each of these experiments.

#### Bernstein and Oxenham (2005) – Effect of harmonic number and phase on pitch discrimination

In the original experiment, the authors measured f0 discrimination thresholds as a function of lowest harmonic number and phase. Stimuli were bandpass-filtered harmonic complexes embedded in masking noise to control the lowest audible harmonic and mask distortion products, with harmonics presented in either sine or random phase. Human listeners performed a three-alternative forced-choice task on each trial within an adaptive procedure for each condition (defined by the combination of lowest harmonic number and phase). On each trial, participants heard three tones: two with the same f0, and the third with a higher f0. They were instructed to identify the tone with the higher pitch. The original experiment found that human f0 thresholds increased with the lowest audible harmonic number, with lower thresholds for sine-phase than random-phase complexes at higher, but not lower, minimum harmonic numbers. The models reproduced the pattern of results seen in humans.

#### Shackleton and Carlyon (1994) – Pitch of alternating-phase harmonic complexes

In the original experiment, the authors employed a pitch-matching paradigm to examine how harmonic resolvability and phase influence pitch perception. Stimuli consisted of bandpass filtered harmonic complexes embedded in masking noise, to constrain the lowest audible harmonic and mask distortion products. Harmonics were presented in an alternating sine–cosine phase. On each trial, participants adjusted the f0 of a sine phase control tone to match the perceived pitch of the alternating phase test stimulus. The matched f0 served as an estimate of the test stimulus’s perceived pitch. The study reported a “pitch-doubling” effect: alternating phase stimuli were judged to be an octave higher than sine phase stimuli, but only when the audible harmonics were high-numbered. No such shift occurred when low-numbered harmonics were audible. The models reproduced this pattern.

#### Moore and Moore (2003) – Pitch of frequency-shifted complexes

In the original experiment, the authors used a pitch-matching paradigm to measure the perceived pitch of frequency-shifted complex tones. The test stimuli were harmonic complexes containing either low- or high-numbered harmonics in cosine phase, rendered inharmonic by shifting all component frequencies by a fixed number in Hz. The study found that the matching f0 shifted linearly with the imposed frequency shift, but only for tones containing low-numbered harmonics. The models reproduced this effect.

#### Moore et al. (1985) – Pitch of complexes with individually mistuned harmonics

In the original experiment, the authors used a pitch-matching paradigm to measure the perceived pitch of complex tones with individually mistuned harmonics. The test stimuli were harmonic complexes containing 12 equal-amplitude, sine-phase harmonics, one of which was mistuned. The study found that small mistunings applied to low-numbered harmonics produced measurable pitch shifts in human listeners. The model reproduced both the magnitude of these shifts and their dependence on harmonic number, though it did not capture the non-monotonic pattern observed in human data.

#### Oxenham et al. (2004) – Frequency discrimination with pure and transposed tones

In the original study, the authors assessed the contribution of temporal cues to pitch perception by measuring discrimination thresholds for pure tones and transposed tones. Transposed tones were generated by multiplying a half-wave rectified, low-frequency sinusoid (the “envelope”) with a high-frequency sinusoidal “carrier”. This procedure preserved the temporal structure of the low-frequency stimulus while shifting its spectral energy to a higher place on the cochlea. The study reported that discrimination thresholds were higher for transposed tones than for pure tones, consistent with the idea that the use of temporal cues is place-dependent. The models reproduced this effect.

### Experiment 7: Pitch Discrimination

#### Human experiment

We measured human uncertainty representations of pitch through a pitch discrimination task. Participants were recruited online via the Prolific platform. Eligibility criteria required the use of headphones, which was verified with a standard headphone-screening test^130^. As an additional headphone check measure, participants also completed an auditory lateralization task that required discrimination of interaural time differences. Only participants who passed both screenings proceeded to the main experiment. Extensive research conducted in our laboratory has found that online data can generally match the quality of data collected in traditional laboratory settings^19,131–136^ provided measures are taken to ensure standardized sound presentation and participant compliance.

Each experimental session began with a screening block using complex tones with harmonics 1–15. Participants were required to achieve at least 70% accuracy in this block to continue with the main experiment. These same practice stimuli were also used as catch trials during the experiment, allowing us to monitor and screen ongoing performance without providing extra practice on the experimental conditions.

The main task used a pitch discrimination and post-decision wagering paradigm. On each trial participants heard two tones: a target followed by a probe, each 100 ms long (10-ms linear on and off ramps) and separated by a 1-s inter-stimulus interval. Participants judged whether the probe tone was higher or lower in pitch relative to the target. The 1-s interstimulus interval was chosen to increase the likelihood that participants would base their judgment on a comparison of f0 estimates, rather than comparing individual frequency components of the stimuli^133,137^. Participants were then asked to place a bet of between 1 to 5 tokens on their response. Correct responses yielded the bet amount of tokens while incorrect responses resulted in the loss of that number of tokens. The total number of tokens a participant accumulated by the end of the experimental session was translated into a bonus cash payment (floored at 0 in the case of negative winnings) at the rate of 0.25 cents per token. No inter-trial feedback was provided.

The stimuli were designed to feature different pitch cues and to produce varying levels of discrimination performance, based on extensive prior work on pitch perception. We employed five stimulus types: (1) pure tones; (2) complex tones with low-numbered harmonics (harmonics 1 – 10); (3) complex tones with intermediate harmonics (harmonics 8 – 17); (4) complex tones with high-numbered harmonics (harmonics 18 – 30) with sine-phase components; and (5) complex tones with high-numbered harmonics (harmonics 18 – 30) with random-phase components. Target and probe tones were offset by fixed F0 differences determined a priori to sufficiently vary task difficulty. For the screening trials, we use F0 differences of 1%, 2%, 3%, and 4.5%. For the main experimental trials, the F0 differences depended on the stimulus type (in order to span a range of performance levels and avoid large numbers of conditions where performance was either near-chance or near-ceiling). For pure tones and complex tones with low or intermediate harmonics, we used differences of 0.5%, 1%, 2%, 3%, and 4.5%. For complex tones with high-numbered harmonics, we used differences of 1%, 3%, 6%, 12% and 24%. Twice as many trials were presented with 3% differences compared to other magnitudes. On each trial, the target F0 was randomly and uniformly sampled within 6% of one of two nominal values (150 or 300 Hz), and the probe was shifted up or down (uniformly at random) from the target according to the chosen F0 difference.

To minimize spectral edge cues, we applied a trapezoidal filter with slopes of 50 dB/octave to all sounds. The filter cutoff was defined relative to the maximum of the target and probe F0s on each trial, translated to the lowest desired harmonic number of that trial type. All sounds were embedded in threshold equalizing noise^125^ with the level set to mask distortion products^70,71^. This was achieved by setting each frequency component of each complex tone to be 10 dB above the level at which a 1 kHz tone was inaudible when presented in the noise. Pure tones were set 20 dB above this audibility level, to ensure that they were easily heard.

Each participant completed a total of 350 trials in the main experiment. Of these, 300 trials (5 stimulus types × 6 F0 differences × 2 nominal F0s × 5 repetitions) comprised the main analyzed dataset. The remaining 50 trials (5 F0 differences × 2 nominal F0s × 5 repetitions) were practice-type catch trials interspersed with the main experiment.

#### Human data analysis

143 participants completed the online experiment. Of these, 34 participants were excluded based on their performance on the catch trials (less than 70% accuracy). We analyzed data from the remaining 109 participants. Psychometric curves were constructed by plotting the mean proportion correct and mean bets as a function of the %F0 difference between the target and probe tones. Error bars represent the standard error of the mean (s.e.m.).

To estimate discrimination thresholds, we fit normal cumulative distribution functions to the mean psychometric function. The F0 threshold was defined as the F0 difference (in percent, capped at 100%) corresponding to a proportion correct of 0.707. Confidence intervals for the thresholds were obtained by bootstrapping across participants (1024 bootstrap samples).

#### Model experiment

To simulate the human experiment on the model, we obtained model F0 estimates for the two stimuli on a trial, and selected the stimulus that yielded the higher F0 estimate. To obtain a pitch estimate from the model for a given stimulus, we computed a maximum a posteriori (MAP) estimate from the stimulus-conditioned posterior distribution predicted by the model. We constrained the estimate to lie within a ±1-octave range centered on the *nominal* F0 value, based on the assumption that humans expected stimuli in the same trial to have relatively close F0s. We obtained model predictions on the target and probe tone independently.

We simulated pitch discrimination trials in this way for 121 target F0s for each of the two nominal F0s (150 Hz and 300 Hz). For each nominal F0, the 121 target F0s were uniformly spaced over ±6%. For each of a set of F0 differences (0.2 to 6 %F0 in steps of 0.2%F0), we then generated a probe tone at that F0 difference from each target F0. We simulated trials for each of the resulting stimulus pairs. Stimulus types, spectral filtering, and the addition of threshold equalizing noise were matched exactly to the human experiment.

Psychometric curves were constructed from the model’s proportion correct responses in the same way as for human participants. Discrimination thresholds were then defined as the F0 difference (in percent, capped at 100%) at which performance reached a proportion correct value of 0.707, using fits of the normal cumulative distribution function to the psychometric data. Inverse threshold data plotted in the manuscript were computed as the mean over 5 different model runs. Error bars represent the standard error of the mean (s.e.m.).

Because the model could only process one stimulus at a time, there was no way to make model bets depend on the %F0 difference between target and probe tones without introducing an ad hoc method of combining the posteriors for the two tones. We deemed it preferable to instead calculate the uncertainty for individual tones and to compare the associated bets to those of humans for a particular F0 difference. This choice was rooted in the assumption that uncertainty would scale monotonically with F0 difference for all stimulus types, such that the diagnostic variation in confidence was that driven by the stimulus type. To summarize the uncertainty of a model prediction that was likely to constrain discrimination, we used a measure of the width of dominant mode of the distribution. The rationale was that determining the f0 difference between tones, when the difference is small, is likely to depend on a comparison of the modes of the posteriors for the two tones. Specifically, we computed the posterior variance within a one-octave window centered on the posterior mode. For each trial, we averaged the posterior variances from the target and probe presentations. This trial-level value was then mapped onto a 1–5 betting scale using an inverse cumulative distribution function (CDF) transform for the distribution of these uncertainties across all trials in the experiment. The mean bets plotted in the manuscript were computed as the mean of 5 different model runs. Error bars represent the standard error of the mean (s.e.m.).

### Analysis of meta-cognitive sensitivity

#### Binning trials by confidence

We defined high- and low-confidence trials based on each participant’s (or model’s) normalized betting behavior. Specifically, we z-scored bets within participant (or model) using the mean and standard deviation computed across all stimuli and locations. Trials were then sorted based on these z-scored values, and those in the upper and lower 40% of the distribution were designated as high- and low-confidence trials, respectively, excluding the central 20% to avoid the median range.

In Experiment 3, the number of trials per subject x stimulus x location combination was limited, and splitting trials into high- and low-confidence conditions further reduced per-bin data. We therefore combined data from adjacent azimuths prior to analysis, rather than computing separate metrics for each azimuth tested. The models had no trials for broadband noise in which bets fell in the low-bet bin, and so the results graph (in Figure 9a) plots only the results for high bets for that stimulus type (and the ANOVA was run on only the narrowband and pure tone conditions).

In Experiment 4, 5 of the 10 participants had at least one confidence x stimulus bin that contained no trials. Mean and SEM d’ values presented in the bar chart in Figure. 9a (Human) are computed over available data in each bin (different numbers of participants, depending on the bin). The models however did not have this issue.

In Experiment 7, several model conditions had data in only some confidence bins. Data are shown for bins that contained more than one trial.

#### Metrics of meta-cognitive sensitivity

We calculated several commonly proposed metrics of meta-cognitive sensitivity, that each in different ways assess the extent to which confidence judgments reflect the quality of sensory evidence: Meta-d’/d’^75^, Brier score^76^, confidence delta^77^, and expected calibration error^77^. Because these metrics rely on the presence of both correct and incorrect trials and on variation in confidence judgments, we only calculated them for the conditions for which performance was above-chance and below-ceiling.

All metrics were computed separately for each participant (or model) and stimulus condition, and subsequently summarized at the group level as mean ± SEM across participants (or models).

Meta-d′/d′ is a signal-detection-theoretic measure of metacognitive efficiency, quantifying how well confidence discriminates between correct and incorrect responses relative to perceptual sensitivity. We calculated the measure using the metadpy Python toolbox^138^, which operates on the observed frequencies of response-confidence combinations. A value of 1 indicates good meta-cognitive sensitivity.

The Brier score measures the mean squared deviation between predicted confidence-derived probabilities and actual outcomes. Confidence judgments (bets) were first mapped to empirical probabilities of correctness on a subject-specific basis. The Brier score was computed as:

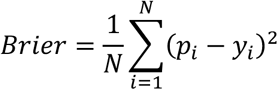

where, *p_i_* is the predicted probability of being correct for trial *i*, and *y_i_*, equal to 0 or 1, indicates whether the response was correct. Lower values indicate better calibration between subjective confidence and objective accuracy.

Confidence delta quantifies whether confidence systematically differs between correct and incorrect trials:

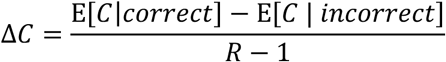

where C is the discrete confidence rating and R is the number of confidence levels. Higher values indicate stronger separation between confidence assigned to correct versus incorrect responses.

Expected calibration error (ECE) measures the discrepancy between confidence and empirical accuracy across confidence bins. Confidence judgments (bets) were first mapped to empirical probabilities of correctness on a subject-specific basis, and were then discretized into K equally spaced bins. For each bin k, let *B_k_* denote the set of trials in that bin. ECE is defined as:

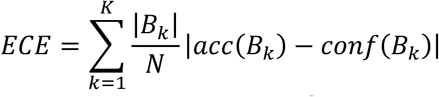

where acc(*B_k_*)is the mean observed accuracy and conf(*B_k_*)is the mean confidence within bin *k*. ECE therefore reflects a weighted average of absolute calibration error across confidence levels, with lower values indicating better calibration.

### Experiment 8: Localization of sources varying in width

#### Human experiment

Participants sat facing a semicircular loudspeaker array, such that their head was centered at 0 ° azimuth and 0 ° elevation. As in Experiment 3, participants were first cued with the elevation from which the sounds would be presented. Elevations were varied in a fully crossed manner, ranging from 0° to 40° in 10° increments. This procedure was used to isolate ambiguity in azimuth perception by holding elevation constant. On each trial, participants indicated the perceived center location of the sound and placed a bet from 1 to 5 cents on their response. No feedback was provided.

We used two types of stimuli: broadband noise and pure tones (of frequencies 600 Hz, 2000 Hz, and 4000 Hz). For the broadband noise condition, we manipulated source width by presenting independently sampled white noise bursts from multiple adjacent loudspeakers in the azimuthal plane. Stimuli were presented at three azimuths (−60°, 0°, and 60°), with source widths of 1, 3, 5, or 7 loudspeakers, always centered on the chosen azimuth. Only odd-numbered widths were included to ensure the presence of a true center. The most peripheral azimuthal location was limited to 60° so that the highest-width condition could be implemented without exceeding the physical bounds of the array. For the pure-tone condition, stimuli were always presented from a single loudspeaker (i.e., width = 1). Wider pure-tone sources were not used, as the superposition of tones from multiple loudspeakers is indistinguishable from a single tone from another location. In total, each participant completed 225 trials: 180 broadband noise trials (3 azimuths x 5 elevations x 4 widths x 3 repeats) + 45 pure tone trials (3 azimuths x 5 elevations x 3 tone frequencies).

Stimuli were 750 ms in duration, modulated by linear on and off ramps (125 ms in duration). All sounds were sampled at 44.1 kHz and presented at a level of 65 dBA.

18 participants completed the experiment (8 female; mean age = 27.4 years, SD = 6.45 years).

#### Model experiment

We simulated the experiment in a virtual replica of the speaker array room, using the procedure described above and employing the same stimuli presented to human participants. To maximize the reliability of the model results, the complete set of sounds and locations was regenerated 32 times, with background noise clips and SNR values randomly sampled for each repeat.

#### Analysis

To evaluate human performance, we plotted the absolute azimuth error as a function of the true azimuthal location, separately for each trial type and width condition. Each data point was obtained by first calculating the median absolute azimuth error for each participant, and then averaging these medians across participants. Error bars represent the standard error of the mean across participants.

### Experiment 9: Source width estimation

#### Human experiment

Participants were played stimuli in exactly the same conditions as in Experiment 7. However, in this experiment, participants were asked to judge the “width” of the sound they heard on a given trial. They chose from options of 1, 3, 5, and 7 speakers, and were told that the speakers playing the sound were physically adjacent in the horizontal plane. Each participant completed 225 trials: 180 broadband noise trials (3 azimuths x 5 elevations x 4 widths x 3 repeats) + 45 pure tone trials (3 azimuths x 5 elevations x 3 tone frequencies). We plotted perceived versus actual width, first averaging responses within subjects and then across subjects; error bars indicate the standard error of the mean.

19 participants with self-reported normal hearing completed the experiment (11 female; mean age = 26.4 years, SD = 7.82 years).

#### Model experiment

We again simulated the experiment in a virtual replica of the speaker array room, using the same stimuli presented to human participants. To maximize the reliability of the model results, the complete set of sounds and locations was regenerated 32 times, with background noise clips and SNR values randomly sampled for each repeat. To extract width judgments from the model, we estimated the spread of the posterior as the 50% highest posterior density interval (in degrees), which we then converted to a number of speakers (10 degrees per speaker, determined by the array spacing).

### Statistics

Repeated measures ANOVAs were implemented using the statsmodels and Pingouin packages for python.

#### Error bars

Error bars in all results figures indicate ±1 standard error of the mean (SEM), computed across participants (for human data) and across the results of five model variants obtained from independent training runs from different random weight initializations (for model data). For the human localization precision data shown in Fig. 8a and 8c, the SEM was estimated by bootstrapping across participants. For the human f0 threshold data in Fig. 7h and 8d, the SEM was estimated by bootstrapping participants (1024 samples), refitting the psychometric function for each sample, and taking the standard deviation of the resulting threshold estimates.

#### Aggregate human-model comparison

To quantify overall correspondence between human and model performance (Fig. 6a&b), we aggregated data from Experiments 3–5. Accuracy measures from Experiments 3 and 5 and sensitivity measures from Experiment 4 were combined after aligning their directionality (sensitivity scores were sign-flipped).

For each measure, data were *z*-scored within each experiment to remove experiment-specific scale differences. Human–model similarity was then quantified as the Pearson correlation (*r*) between human and model data, with 95% confidence intervals obtained by bootstrapping across stimulus conditions (using the same bootstrap samples for humans and models to preserve within-condition correspondence between human and model results).

#### Comparison of uncertainty-aware and cross-entropy model

To test whether the uncertainty-aware model correlated more strongly with human data than the softmax model, we used a one-tailed paired bootstrap test (1024 iterations). For each bootstrap sample, stimulus conditions were resampled with replacement, and the difference in correlation between the two models (Δ*r* = *r*(*M_reg_*) − *r*(*M_cls_*)) was computed. The one-tailed p-value was defined as the proportion of bootstrap samples where Δ*r* ≤ 0.

To assess whether differences in human-model confidence correspondence were a side effect of human-model accuracy correspondence, we repeated the analysis on a subset of stimulus conditions chosen such that the correspondence between human and model accuracy was approximately matched for the uncertainty-aware and softmax models. This matching was achieved by discarding 20% of the experimental conditions, selected to yield similar human-model accuracy correspondence for the two models. To avoid problems with non-independence, condition selection was performed using one randomly chosen half of the human participants together with one randomly chosen half of the model trials, and the subsequent analysis was carried out on the held-out human participants and model trials. Thus, the data used to determine which conditions to retain were independent from the data used to evaluate confidence differences.

### Informed consent

All participants gave informed consent, and all experiments were approved by the Massachusetts Institute of Technology Committee on the Use of Humans as Experimental Subjects (COUHES).

## Data availability

The processed, anonymized human data and simulation data from this study are available at the project repository. There are no restrictions on data availability, and all relevant files are provided in CSV format. No data with mandated deposition are included in this study.

## Code availability

The code used for modeling and data analysis in this study is available at the project repository.

## Acknowledgments

Research supported by National Institutes of Health grant R01 DC017970. The authors thank Adanna Thomas for assistance running Experiments 3, 4, 5, 7 and 8, Iman Khanani for assistance running Experiment 6, Mark Saddler for assistance with pitch model training data and architectures, Andrew Francl and R. Preston Hess for collecting most of the data in Experiment 1, Ajani Stewart for brainstorming during the initial stages of the project, and the McDermott lab, Zoe Boundy-Singer and Ila Fiete for helpful feedback on the manuscript.

## Supplementary Information for Govindarajan et al

**Supplementary Figure 1.**
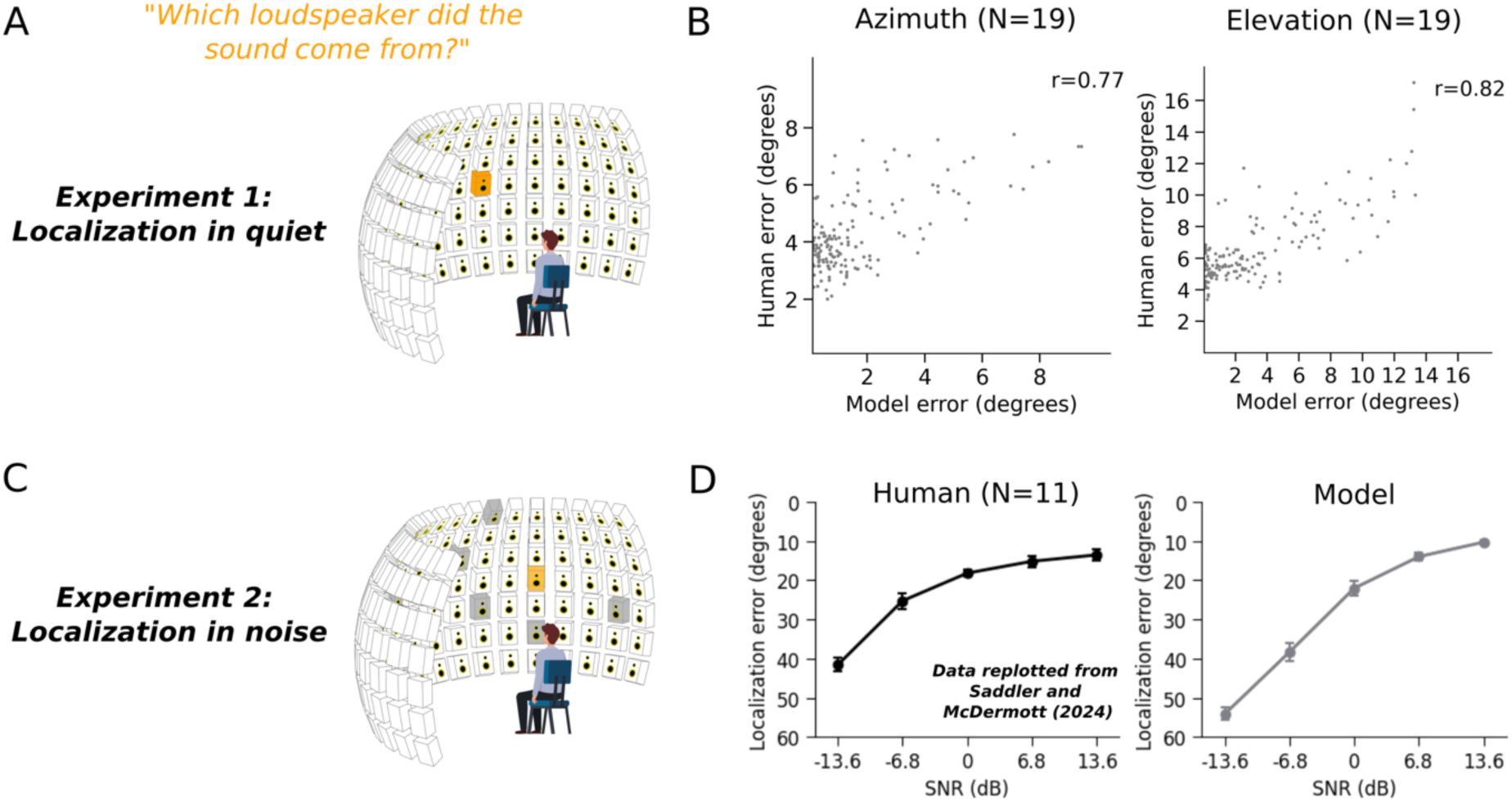
Validation of model of sound localization. **A**. Experiment 1 (measuring localization in quiet). Single natural sounds were presented from one of 133 locations, and the participant judged the location. **B**. Comparison of human and model localization accuracy for individual sounds, calculated and plotted separately for azimuth (left) and elevation (right). **C**. Experiment 2 (measuring localization in noise). Natural sounds were presented from one of 133 locations, accompanied by noise samples (textures) played from some number of other locations. The participant judged the location of the natural sound. **D**. Comparison of human and model azimuthal localization accuracy as a function of signal-to-noise ratio (SNR). Human data are replotted from a previous publication^20^.

**Supplementary Figure 2.**
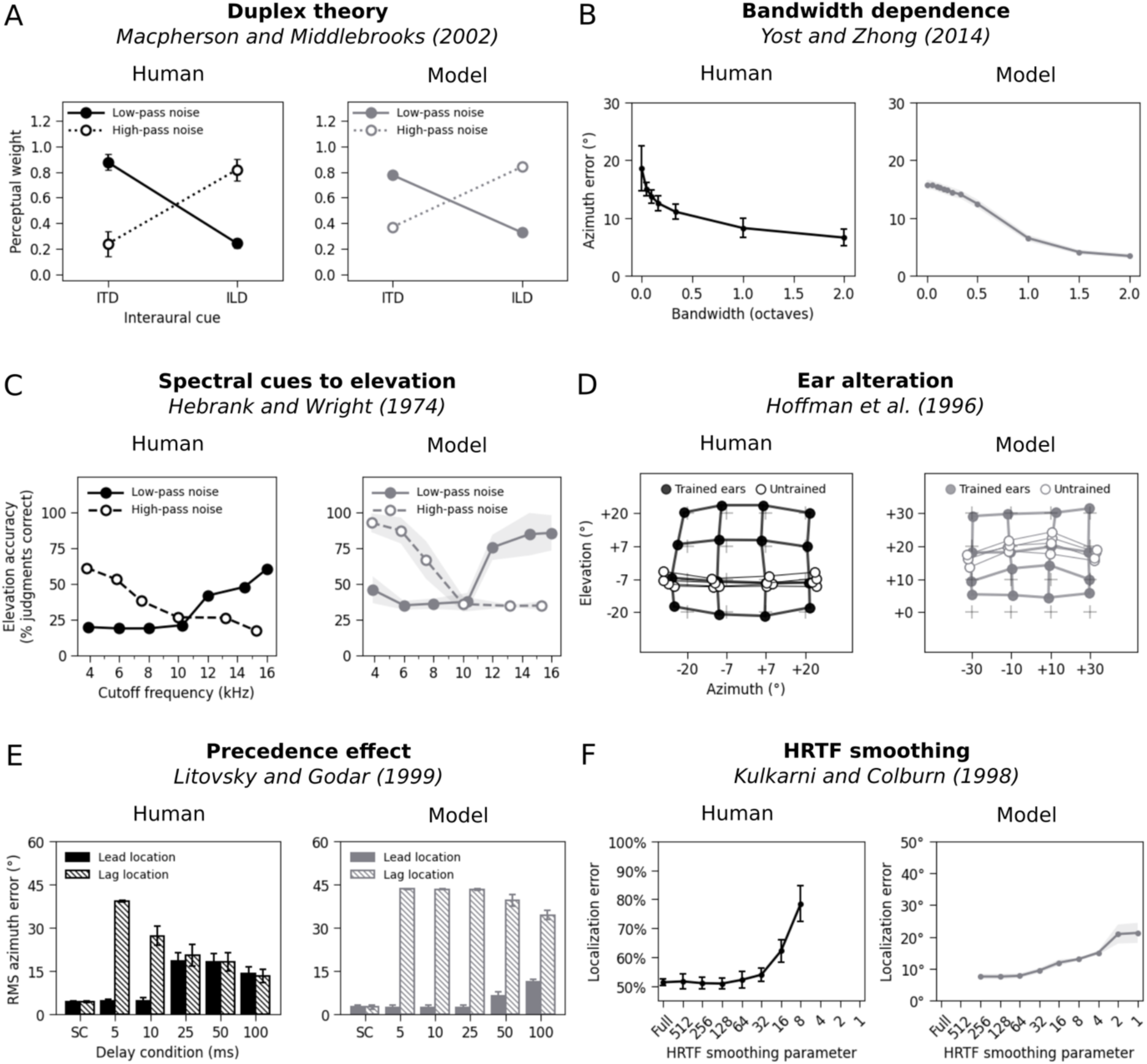
Validation of sound localization model. Panels plots human and model results from the battery of experiments used by Francl and McDermott (2022)^23^ for model validation. Each experiment captures an aspect of human spatial hearing. The model qualitatively reproduces each effect.

**Supplementary Figure 3.**
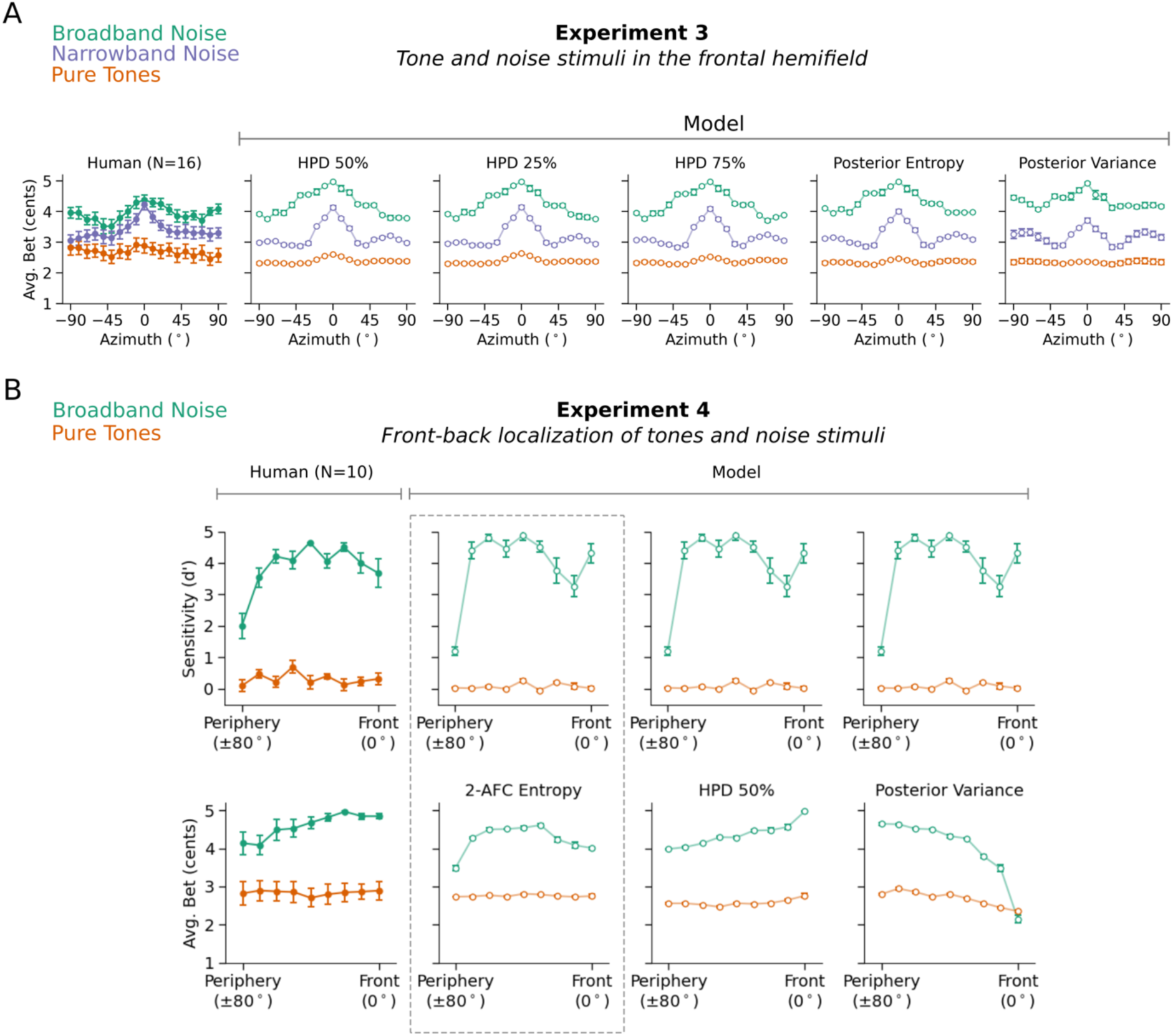
Effect of alternative summary measures of uncertainty. **A.** Model confidence judgments (bets) for Experiment 3 for several summary measures of uncertainty. In each case the summary measure was converted to bets using the same procedure (see Methods). **B.** Model confidence judgments (bets) for Experiment 4, for several summary measures of uncertainty. Here, because the experiment involves a front-back judgment, different summary measures yield different results. In particular, the posterior variance (i.e., the variance over the full posterior distribution over all sound locations) diverges from human confidence judgments, plausibly because it is not closely related to uncertainty of the front-back distinction assayed by the experiment.

**Supplementary Figure 4.**
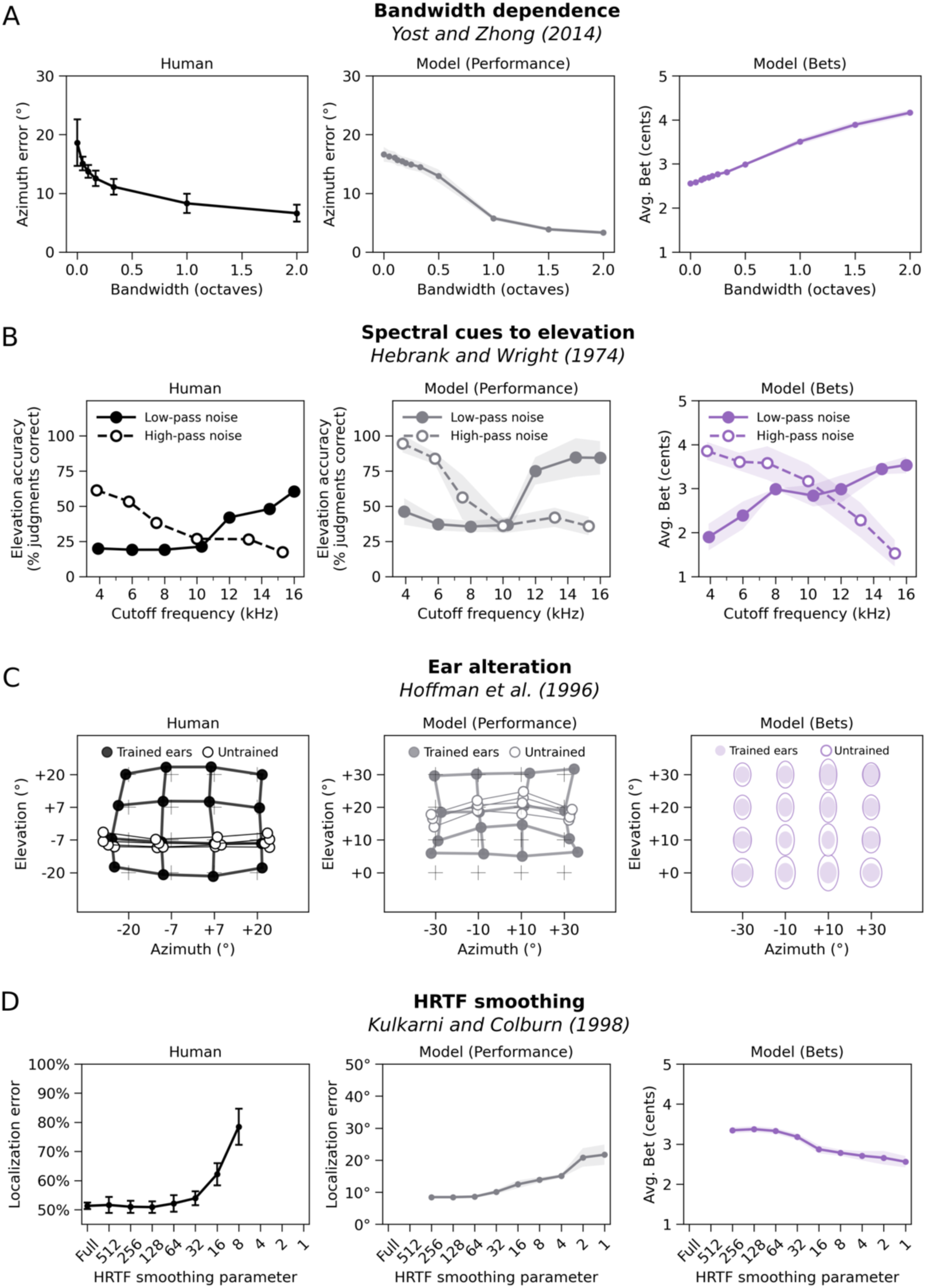
Model confidence judgments across battery of classic localization experiments. Right panels plots model bets for experiments used by Francl and McDermott (2022)^23^ for model validation (including the experiments in which the dependent measure is the accuracy of localization for a single sound). Left and center columns are replicated from Supplementary Figure 2 for ease of comparison. Each experiment manipulates something different about the stimulus. Note that some of the graphs plot error, whereas others plot accuracy. The model uncertainty varies inversely with localization performance in each case. The ellipses in the bet plot in C plot the inverse of the bets for azimuthal and elevation judgments. Bets were scaled by a constant factor to make the variation across conditions visible.

**Supplementary Figure 5.**
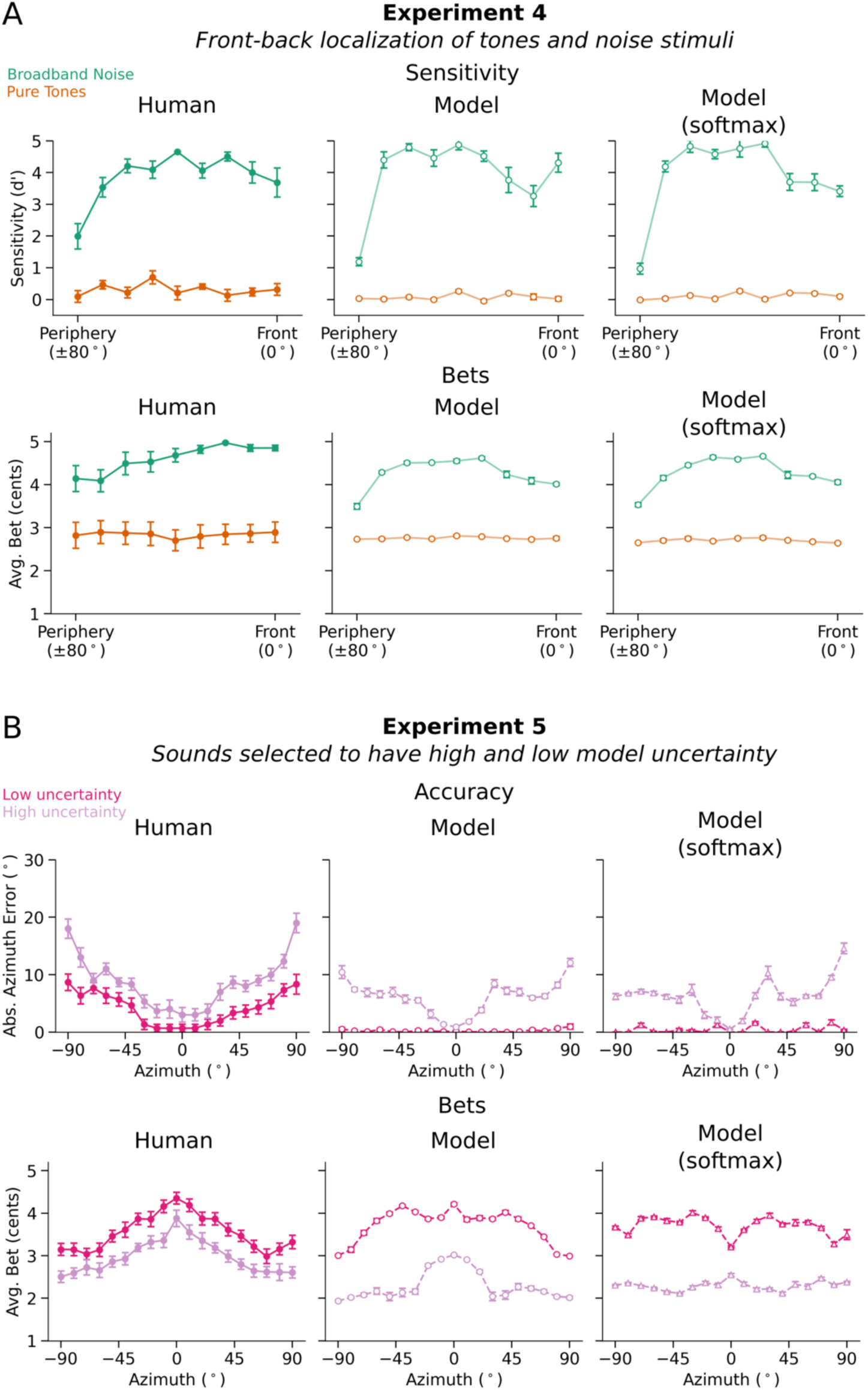
Comparison of human-model similarity between an uncertainty-aware model trained with a log-likelihood loss function and a classification model trained with cross-entropy loss. **A.** Experiment 4: Front-back localization of tones and noise stimuli. **B.** Experiment 5: Natural sounds screened by model uncertainty. Error bars plot SEM.

**Supplementary Figure 6.**
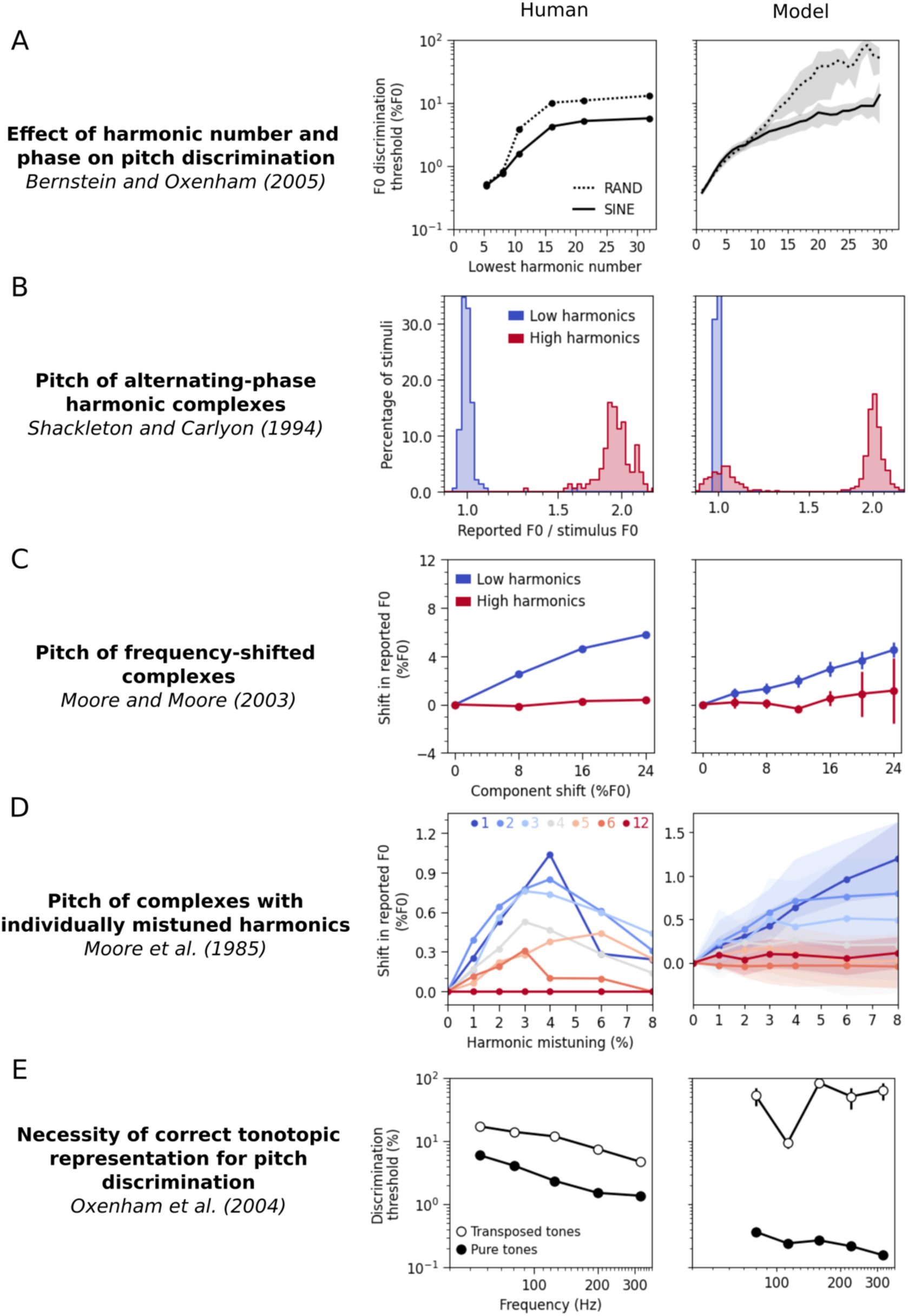
Validation of pitch estimation model. Each panel plots human and model results from the battery of experiments used by Saddler et al. (2021)^24^ for model validation. Each experiment captures an aspect of human pitch perception. The model qualitatively reproduces each effect.

**Supplementary Figure 7.**
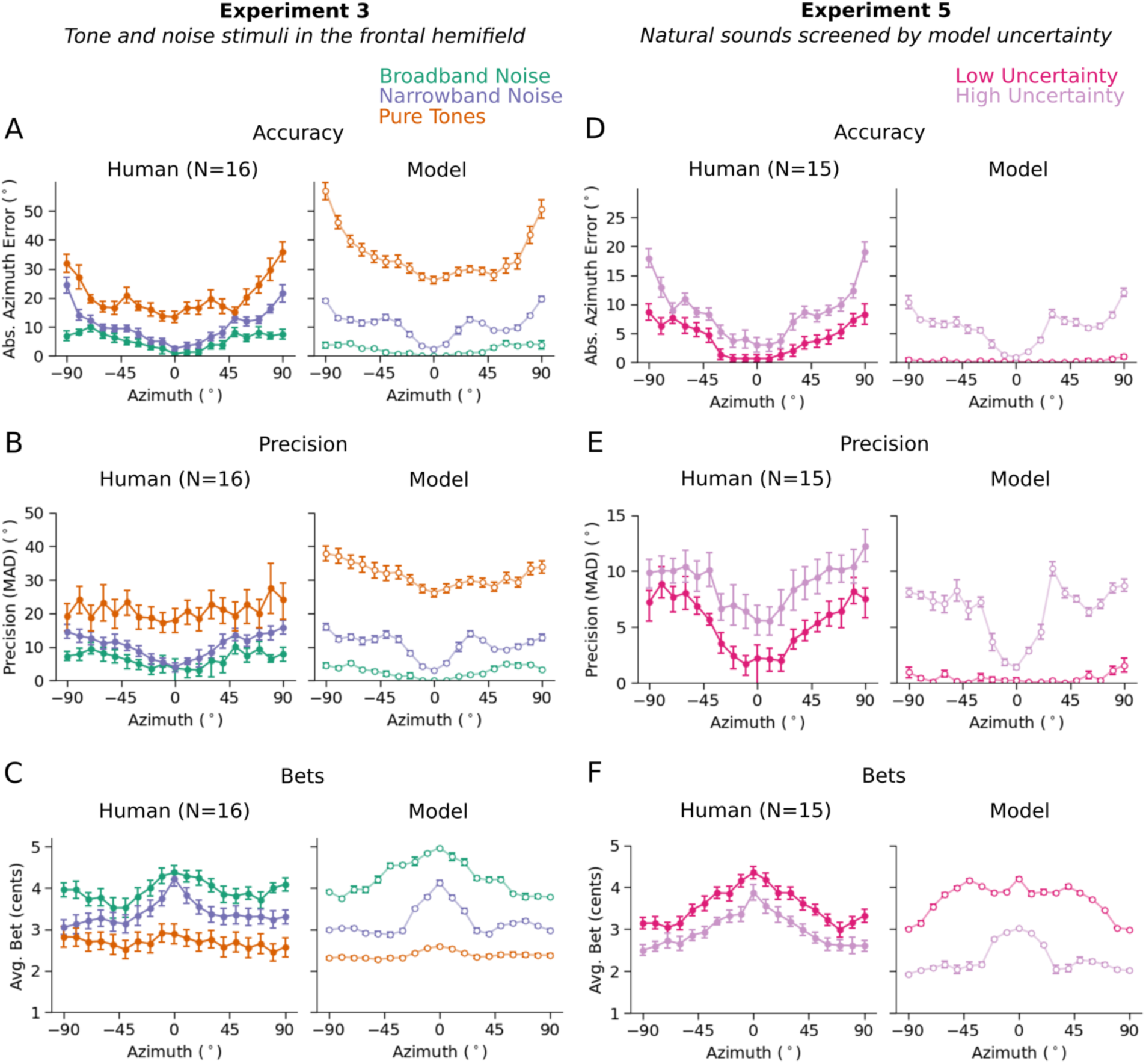
Human (left panels) and model (right panels) performance as a function of azimuth for Experiments 3 and 5. **A&D.** Localization accuracy for each stimulus type at each azimuthal position. **B&E.** Localization precision for each stimulus type at each azimuthal position. **C&F.** Bets for each stimulus type and azimuthal position. Error bars plot SEM.

**Supplementary Table 1.** Sound localization model architecture.

| Layer | Type | Input Shape | Output Shape | Params | Details |
| --- | --- | --- | --- | --- | --- |
| 000_Conv2d | Conv2d | [1, 2, 40, 8000] | [1, 32, 38, 7969] | 6144 | k=(3, 32), s=(1, 1), p=(0, 0) |
| 001_MaxPool2d | MaxPool2d | [1, 32, 38, 7969] | [1, 32, 38, 7969] | 0 | k=(1, 1), s=(1, 1) |
| 002_ReLU | ReLU | [1, 32, 38, 7969] | [1, 32, 38, 7969] | 0 |  |
| 003_BatchNorm2d | BatchNorm2d | [1, 32, 38, 7969] | [1, 32, 38, 7969] | 64 |  |
| 004_Conv2d | Conv2d | [1, 32, 38, 7969] | [1, 32, 37, 7954] | 32768 | k=(2, 16), s=(1, 1), p=(0, 0) |
| 005_MaxPool2d | MaxPool2d | [1, 32, 37, 7954] | [1, 32, 37, 1988] | 0 | k=(1, 4), s=(1, 4) |
| 006_ReLU | ReLU | [1, 32, 37, 1988] | [1, 32, 37, 1988] | 0 |  |
| 007_BatchNorm2d | BatchNorm2d | [1, 32, 37, 1988] | [1, 32, 37, 1988] | 64 |  |
| 008_Conv2d | Conv2d | [1, 32, 37, 1988] | [1, 64, 36, 1957] | 131072 | k=(2, 32), s=(1, 1), p=(0, 0) |
| 009_MaxPool2d | MaxPool2d | [1, 64, 36, 1957] | [1, 64, 36, 1957] | 0 | k=(1, 1), s=(1, 1) |
| 010_ReLU | ReLU | [1, 64, 36, 1957] | [1, 64, 36, 1957] | 0 |  |
| 011_BatchNorm2d | BatchNorm2d | [1, 64, 36, 1957] | [1, 64, 36, 1957] | 128 |  |
| 012_Conv2d | Conv2d | [1, 64, 36, 1957] | [1, 64, 34, 1954] | 49152 | k=(3, 4), s=(1, 1), p=(0, 0) |
| 013_MaxPool2d | MaxPool2d | [1, 64, 34, 1954] | [1, 64, 34, 488] | 0 | k=(1, 4), s=(1, 4) |
| 014_ReLU | ReLU | [1, 64, 34, 488] | [1, 64, 34, 488] | 0 |  |
| 015_BatchNorm2d | BatchNorm2d | [1, 64, 34, 488] | [1, 64, 34, 488] | 128 |  |
| 016_Conv2d | Conv2d | [1, 64, 34, 488] | [1, 128, 32, 481] | 196608 | k=(3, 8), s=(1, 1), p=(0, 0) |
| 017_MaxPool2d | MaxPool2d | [1, 128, 32, 481] | [1, 128, 32, 120] | 0 | k=(1, 4), s=(1, 4) |
| 018_ReLU | ReLU | [1, 128, 32, 120] | [1, 128, 32, 120] | 0 |  |
| 019_BatchNorm2d | BatchNorm2d | [1, 128, 32, 120] | [1, 128, 32, 120] | 256 |  |
| 020_Conv2d | Conv2d | [1, 128, 32, 120] | [1, 256, 30, 119] | 196608 | k=(3, 2), s=(1, 1), p=(0, 0) |
| 021_MaxPool2d | MaxPool2d | [1, 256, 30, 119] | [1, 256, 30, 59] | 0 | k=(1, 2), s=(1, 2) |
| 022_ReLU | ReLU | [1, 256, 30, 59] | [1, 256, 30, 59] | 0 |  |
| 023_BatchNorm2d | BatchNorm2d | [1, 256, 30, 59] | [1, 256, 30, 59] | 512 |  |
| 024_Conv2d | Conv2d | [1, 256, 30, 59] | [1, 512, 29, 52] | 2097152 | k=(2, 8), s=(1, 1), p=(0, 0) |
| 025_MaxPool2d | MaxPool2d | [1, 512, 29, 52] | [1, 512, 29, 52] | 0 | k=(1, 1), s=(1, 1) |
| 026_ReLU | ReLU | [1, 512, 29, 52] | [1, 512, 29, 52] | 0 |  |
| 027_BatchNorm2d | BatchNorm2d | [1, 512, 29, 52] | [1, 512, 29, 52] | 1024 |  |
| 028_Conv2d | Conv2d | [1, 512, 29, 52] | [1, 512, 27, 49] | 3145728 | k=(3, 4), s=(1, 1), p=(0, 0) |
| 029_MaxPool2d | MaxPool2d | [1, 512, 27, 49] | [1, 512, 27, 24] | 0 | k=(1, 2), s=(1, 2) |
| 030_ReLU | ReLU | [1, 512, 27, 24] | [1, 512, 27, 24] | 0 |  |
| 031_BatchNorm2d | BatchNorm2d | [1, 512, 27, 24] | [1, 512, 27, 24] | 1024 |  |
| 032_Conv2d | Conv2d | [1, 512, 27, 24] | [1, 512, 27, 22] | 786432 | k=(1, 3), s=(1, 1), p=(0, 0) |
| 033_MaxPool2d | MaxPool2d | [1, 512, 27, 22] | [1, 512, 27, 22] | 0 | k=(1, 1), s=(1, 1) |
| 034_ReLU | ReLU | [1, 512, 27, 22] | [1, 512, 27, 22] | 0 |  |
| 035_BatchNorm2d | BatchNorm2d | [1, 512, 27, 22] | [1, 512, 27, 22] | 1024 |  |
| 036_Linear | Linear | [1, 304128] | [1, 512] | 155714048 | in=304128, out=512 |
| 037_Linear | Linear | [1, 304128] | [1, 512] | 155714048 | in=304128, out=512 |
| 038_ReLU | ReLU | [1, 512] | [1, 512] | 0 |  |
| 039_BatchNorm1d | BatchNorm1d | [1, 512] | [1, 512] | 1024 |  |
| 040_Dropout | Dropout | [1, 512] | [1, 512] | 0 |  |
| 041_Linear | Linear | [1, 512] | [1, 35] | 17955 | in=512, out=35 |
| 042_Linear | Linear | [1, 512] | [1, 35] | 17955 | in=512, out=35 |
| 043_Softmax | Softmax | [1, 5] | [1, 5] | 0 |  |
| 044_Tanh | Tanh | [1, 20] | [1, 20] | 0 |  |
| 045_ReLUX | ReLUX | [1, 10] | [1, 10] | 0 |  |

**Supplementary Table 2.**
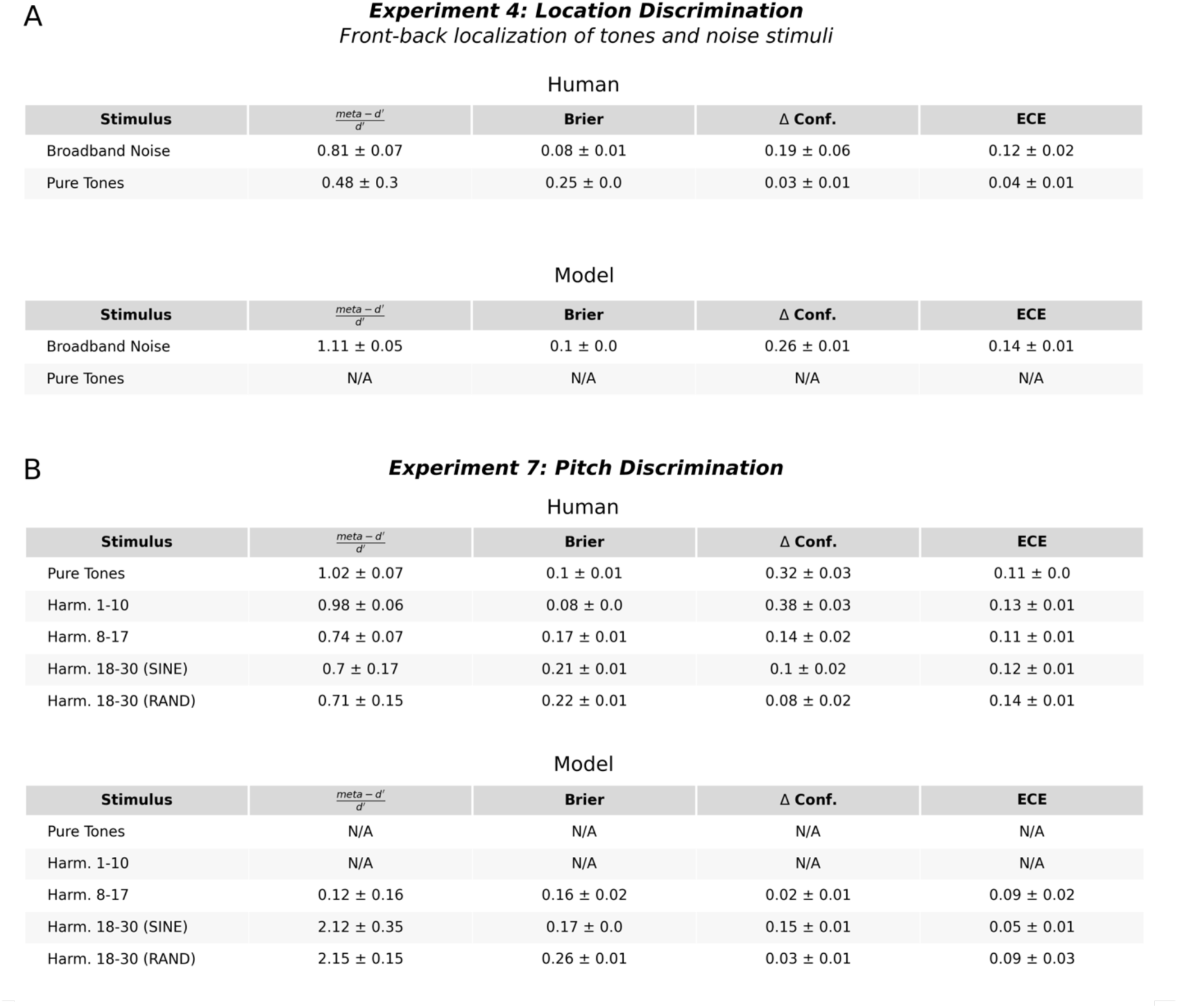
Metrics of meta-cognitive sensitivity. N/A indicates that performance was either at chance or at ceiling (which was the case for come conditions for the model), in which case the metrics were not applicable.

| Human |  |  |  |  |
| --- | --- | --- | --- | --- |
| Stimulus | $\frac{\text{meta} - d'}{d'}$ | Brier | $\Delta$ Conf. | ECE |
| Broadband Noise | $0.81 \pm 0.07$ | $0.08 \pm 0.01$ | $0.19 \pm 0.06$ | $0.12 \pm 0.02$ |
| Pure Tones | $0.48 \pm 0.3$ | $0.25 \pm 0.0$ | $0.03 \pm 0.01$ | $0.04 \pm 0.01$ |

| Model |  |  |  |  |
| --- | --- | --- | --- | --- |
| Stimulus | $\frac{\text{meta} - d'}{d'}$ | Brier | $\Delta$ Conf. | ECE |
| Broadband Noise | $1.11 \pm 0.05$ | $0.1 \pm 0.0$ | $0.26 \pm 0.01$ | $0.14 \pm 0.01$ |
| Pure Tones | N/A | N/A | N/A | N/A |

**Experiment 7: Pitch Discrimination**
| Human |  |  |  |  |
| --- | --- | --- | --- | --- |
| Stimulus | $\frac{\text{meta} - d'}{d'}$ | Brier | $\Delta$ Conf. | ECE |
| Pure Tones | $1.02 \pm 0.07$ | $0.1 \pm 0.01$ | $0.32 \pm 0.03$ | $0.11 \pm 0.0$ |
| Harm. 1-10 | $0.98 \pm 0.06$ | $0.08 \pm 0.0$ | $0.38 \pm 0.03$ | $0.13 \pm 0.01$ |
| Harm. 8-17 | $0.74 \pm 0.07$ | $0.17 \pm 0.01$ | $0.14 \pm 0.02$ | $0.11 \pm 0.01$ |
| Harm. 18-30 (SINE) | $0.7 \pm 0.17$ | $0.21 \pm 0.01$ | $0.1 \pm 0.02$ | $0.12 \pm 0.01$ |
| Harm. 18-30 (RAND) | $0.71 \pm 0.15$ | $0.22 \pm 0.01$ | $0.08 \pm 0.02$ | $0.14 \pm 0.01$ |

**Supplementary Table 2.** Metrics of meta-cognitive sensitivity. N/A indicates that performance was either at chance or at ceiling (which was the case for some conditions for the model), in which case the metrics were not applicable.
| Model |  |  |  |  |
| --- | --- | --- | --- | --- |
| Stimulus | $\frac{\text{meta} - d'}{d'}$ | Brier | $\Delta$ Conf. | ECE |
| Pure Tones | N/A | N/A | N/A | N/A |
| Harm. 1-10 | N/A | N/A | N/A | N/A |
| Harm. 8-17 | $0.12 \pm 0.16$ | $0.16 \pm 0.02$ | $0.02 \pm 0.01$ | $0.09 \pm 0.02$ |
| Harm. 18-30 (SINE) | $2.12 \pm 0.35$ | $0.17 \pm 0.0$ | $0.15 \pm 0.01$ | $0.05 \pm 0.01$ |
| Harm. 18-30 (RAND) | $2.15 \pm 0.15$ | $0.26 \pm 0.01$ | $0.03 \pm 0.01$ | $0.09 \pm 0.03$ |

**Supplementary Table 3.**
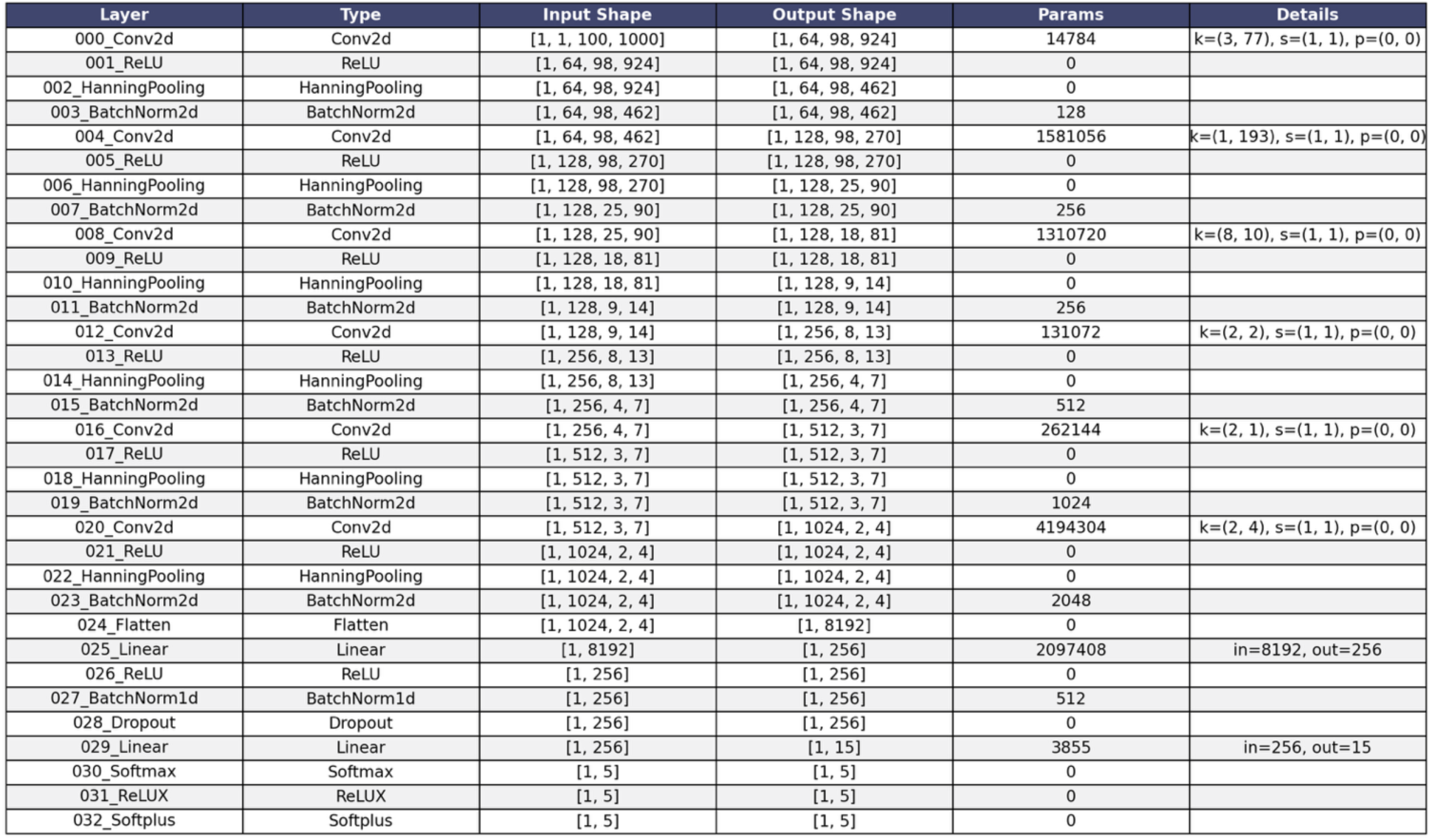
Pitch estimation model architecture.

## References

1 Sahani, M. & Dayan, P. Doubly distributional population codes: Simultaneous representation of uncertainty and multiplicity. Neural Computation 15, 2255–2279 (2003).

2 Pouget, A., Beck, J. M., Ma, W. J. & Latham, P. E. Probabilistic brains: knowns and unknowns. Nature Neuroscience 16, 1170–1178 (2013).

3 Rich, D., Cazettes, F., Wang, Y., Pena, J. L. & Fischer, B. J. Neural representation of probabilities for Bayesian inference. Journal of Computational Neuroscience 38, 315–323 (2015).

4 Landy, M. S., Maloney, L. T., Johnston, E. B. & Young, M. Measurement and modeling of depth cue combination: in defense of weak fusion. Vision Research 35, 389–412, 10.1016/0042-6989(94)00176-M (1995).

5 Ernst, M. O. & Banks, M. S. Humans integrate visual and haptic information in a statistically optimal fashion. Nature 415, 429–433 (2002).

6 Weiss, Y., Simoncelli, E. P. & Adelson, E. H. Motion illusions as optimal percepts. Nature Neuroscience 5, 598–604, doi:10.1038/nn0602-858 (2002).

7 Pouget, A., Drugowitsch, J. & Kepecs, A. Confidence and certainty: distinct probabilistic quantities for different goals. Nature Neuroscience 19, 366–374 (2016).

8 Yeung, N. & Summerfield, C. Metacognition in human decision-making: confidence and error monitoring. Philosophical Transactions of the Royal Society B 367, 1310–1321 (2012).

9 Kepecs, A. & Mainen, Z. F. A computational framework for the study of confidence in humans and animals. Philosophical Transactions of the Royal Society B 367, 1322–1337 (2012).

10 Fleming, S. M. & Lau, H. C. How to measure metacognition. Frontiers in Human Neuroscience 8, 443 (2014).

11 Mamassian, P. Visual confidence. Annual Review of Vision Science 2, 459–481 (2016).

12 Koriat, A. The self-consistency model of subjective confidence. Psychological Review 119, 80–113 (2012).

13 Zylberberg, A., Fetsch, C. R. & Shadlen, M. N. The influence of evidence volatility on choice, reaction time and confidence in a perceptual decision. eLife 5, e17688, doi:10.7554/eLife.17688 (2016).

14 Adler, W. T. & Ma, W. J. Comparing Bayesian and non-Bayesian accounts of human confidence reports. PLoS Comp. Biol. 14, e1006572, doi:10.1371/journal.pcbi.1006572 (2018).

15 Caziot, B. & Mamassian, P. Perceptual confidence judgments reflect self-consistency. Journal of Vision 21, 8 (2021).

16 Boundy-Singer, Z. M., Ziemba, C. M. & Goris, R. L. T. Confidence reflects a noisy decision reliability estimate. Nature Human Behaviour 7, 142–154 (2023).

17 Richards, B. A., Lillicrap, T. P., Beaudoin, P., Bengio, Y., Bogacz, R., Christensen, A., Clopath, C., Costa, R. P., de Berker, A., Ganguli, S., Gillon, C. J., Hafner, D., Kepecs, A., Kriegeskorte, N., Latham, P., Lindsay, G. W., Miller, K. D., Naud, R., Pack, C. C., Poirazi, P., Roelfsema, P., Sacramento, J., Saxe, A., Scellier, B., Schapiro, A. C., Senn, W., Wayne, G., Yamins, D., Zenke, F., Zylberberg, J., Therien, D. & Kording, K. P. A deep learning framework for neuroscience. Nature Neuroscience 22, 1761–1770, doi:10.1038/s41593-019-0520-2 (2019).

18 Rajalingham, R., Issa, E., Bashivan, P., Kar, K., Schmidt, K. & DiCarlo, J. Large-scale, high-resolution comparison of the core visual object recognition behavior of humans, monkeys, and state-of-the-art deep artificial neural networks. Journal of Neuroscience 38, 7255–7269 (2018).

19 Kell, A. J. E., Yamins, D. L. K., Shook, E. N., Norman-Haignere, S. & McDermott, J. H. A task-optimized neural network replicates human auditory behavior, predicts brain responses, and reveals a cortical processing hierarchy. Neuron 98, 630–644, doi:10.1016/j.neuron.2018.03.044 (2018).

20 Saddler, M. R. & McDermott, J. H. Models optimized for real-world tasks reveal the necessity of precise temporal coding in hearing. Nature Communications 15, 10590, doi:10.1038/s41467-024-54700-5 (2024).

21 Alavilli, S. & McDermott, J. H. From sound to source: Human and model recognition of environmental sounds. bioRxiv, 2026.2003.2012.711349, doi:10.64898/2026.03.12.711349 (2026).

22 Griffith, I. M., Hess, R. P. & McDermott, J. H. Optimized feature gains explain and predict successes and failures of human selective listening. Nature Human Behaviour 10, doi:10.1038/s41562-026-02414-7 (2026).

23 Francl, A. & McDermott, J. H. Deep neural network models of sound localization reveal how perception is adapted to real-world environments. Nature Human Behavior 6, 111–133 (2022).

24 Saddler, M. R., Gonzalez, R. & McDermott, J. H. Deep neural network models reveal interplay of peripheral coding and stimulus statistics in pitch perception. Nature Communications 12, 7278, doi:10.1038/s41467-021-27366-6 (2021).

25 Sensoy, M., Kaplan, L. & Kandemir, M. in Proceedings of the 32nd International Conference on Neural Information Processing Systems 3183–3193 (Curran Associates Inc., 2018).

26 Abdar, M., Pourpanah, F., Hussain, S., Rezazadegan, D., Liu, L., Ghavamzadeh, M., Fieguth, P., Cao, X., Khosravi, A., Acharya, U. R., Makarenkov, V. & Nahavandi, S. A review of uncertainty quantification in deep learning: Techniques, applications and challenges. Information Fusion 76, 243–297, 10.1016/j.inffus.2021.05.008 (2021).

27 Gawlikowski, J., Tassi, C. R. N., Ali, M., Lee, J., Humt, M., Feng, J., Kruspe, A., Triebel, R., Jung, P., Roscher, R., Shahzad, M., Yang, W., Bamler, R. & Zhu, X. X. A survey of uncertainty in deep neural networks. Artificial Intelligence Review 56, 1513–1589, doi:10.1007/s10462-023-10562-9 (2023).

28 Murphy, K. P. Probabilistic Machine Learning: An Introduction. (MIT Press, 2023).

29 Blauert, J. Spatial hearing: The psychophysics of human sound localization. (MIT Press, 1997).

30 Goodfellow, I., Bengio, Y. & Courville, A. Deep Learning. (MIT Press, 2016).

31 Terven, J., Cordova-Esparza, D. M., Romero-González, J. A., Ramírez-Pedraza, A. & Chávez-Urbiola, E. A. A comprehensive survey of loss functions and metrics in deep learning. Artificial Intelligence 58, 195 (2025).

32 Huber, P. J. & Ronchetti, E. M. Robust Statistics. (Wiley, 2009).

33 Anthony, M. & Bartlett, P. L. Neural Network Learning: Theoretical Foundations. (Cambridge University Press, 1999).

34 Rothe, R., Timofte, R. & Van Gool, L. Deep expectation of real and apparent age from a single image without facial landmarks. International Journal of Computer Vision 126, 144–157, doi:10.1007/s11263-016-0940-3 (2018).

35 Fu, H., Gong, M., Wang, C., Batmanghelich, K. & Tao, D. in Proceedings of the IEEE Conference on Computer Vision and Pattern Recognition (CVPR) (2018).

36 Bishop, C. M. Mixture density networks. Neural Computing Research Group Report (1994).

37 Zen, H. & Senior, A. in 2014 IEEE International Conference on Acoustics, Speech and Signal Processing (ICASSP). 3844–3848 (IEEE).

38 Prokudin, S., Gehler, P. & Nowozin, S. in The European Conference on Computer Vision (ECCV 2018) 534–551 (Springer, Munich, Germany, 2018).

39 Makansi, O., Ilg, E., Cicek, O. & Brox, T. in The IEEE/CVF Conference on Computer Vision and Pattern Recognition 7144-7153 (IEEE, 2019).

40 Wallach, H. The role of head movements and vestibular and visual cues in sound localization. Journal of Experimental Psychology 27, 339–368, doi:10.1037/h0054629 (1940).

41 Payne, R. S. Acoustic location of prey by barn owls (tyto alba). Journal of Experimental Biology 54, 535–573 (1971).

42 Lertpoompunya, A., Ozmeral, E. J., Higgins, N. C. & Eddins, D. A. Head-orienting behaviors during simultaneous speech detection and localization. Frontiers in Psychology 15, 2024.1425972, doi:10.3389/fpsyg.2024.1425972 (2024).

43 Stevens, S. S. & Newman, E. B. The localization of actual sources of sound. The American Journal of Psychology 48, 297–306 (1936).

44 Hicks, J. M. & McDermott, J. H. Noise schemas aid hearing in noise. Proceedings of the National Academy of Sciences 121, e2408995121, 10.1073/pnas.2408995121 (2024).

45 Macpherson, E. A. & Middlebrooks, J. C. Listener weighting of cues for lateral angle: The duplex theory of sound localization revisited. Journal of the Acoustical Society of America 111, 2219–2236, doi:10.1121/1.1471898 (2002).

46 Yost, W. A. & Zhong, X. Sound source localization identification accuracy: Bandwidth dependencies. Journal of the Acoustical Society of America 136, 2737–2746, doi:10.1121/1.4898045 (2014).

47 Hebrank, J. & Wright, D. Spectral cues used in the localization of sound sources on the median plane. Journal of the Acoustical Society of America 56, 1829–1834, doi:10.1121/1.1903520 (1974).

48 Kulkarni, A. & Colburn, H. S. Role of spectral detail in sound-source localization. Nature 396, 747–749, doi:10.1038/25526 (1998).

49 Hofman, P. M., Van Riswick, J. G. A. & van Opstal, A. J. Relearning sound localization with new ears. Nature Neuroscience 1, 417–421, doi:10.1038/1633 (1998).

50 Litovsky, R. Y., Colburn, H. S., Yost, W. A. & Guzman, S. J. The precedence effect. Journal of the Acoustical Society of America 106, 1633–1654, doi:10.1121/1.427914 (1999).

51 Litovsky, R. Y. & Godar, S. P. Difference in precedence effect between children and adults signifies development of sound localization abilities in complex listening tasks. Journal of the Acoustical Society of America 128, 1979–1991, doi:10.1121/1.3478849 (2010).

52 Brown, A. D., Stecker, G. C. & Tollin, D. J. The precedence effect in sound localization. Journal of the Association for Research in Otolaryngology 16, 1–28, doi:10.1007/s10162-014-0496-2 (2015).

53 Yost, W. A. Cones-of-confusions: Are listeners confused? Journal of the Acoustical Society of America 154, 2769–2771 (2023).

54 Wenzel, E. M., Arruda, M., Kistler, D. J. & Wightman, F. L. Localization using nonindividualized head-related transfer functions. Journal of the Acoustical Society of America 94, 111–123, doi:10.1121/1.407089 (1993).

55 Mills, A. W. On the minimum audible angle. Journal of the Acoustical Society of America 30, 237–246, doi:10.1121/1.1909553 (1958).

56 Macaulay, E. J., Hartmann, W. M. & Rakerd, B. The acoustical bright spot and mislocalization of tones by human listeners. Journal of the Acoustical Society of America 127, 1440–1449 (2010).

57 Gelman, A., Stern, H. S., Carlin, J. B., Dunson, D. B., Vehtari, A. & Rubin, D. B. Bayesian Data Analysis. (Chapman and Hall, 2013).

58 Makous, J. C. & Middlebrooks, J. C. Two-dimensional sound localization by human listeners. Journal of the Acoustical Society of America 87, 2188–2200 (1990).

59 Wightman, F. L. & Kistler, D. J. Resolution of front–back ambiguity in spatial hearing by listener and source movement. Journal of the Acoustical Society of America 105, 2841–2853 (1999).

60 Yost, W. A. & Pastore, M. T. Individual listener differences in azimuthal front-back reversals. Journal of the Acoustical Society of America 146, 2709–2715 (2019).

61 Geurts, L. S., Cooke, J. R. H., van Bergen, R. S. & Jehee, J. F. M. Subjective confidence reflects representation of Bayesian probability in cortex. Nature Human Behaviour 6, 294–305 (2022).

62 Plack, C. J., Oxenham, A. J., Popper, A. J. & Fay, R. R. in *Springer Handbook of Auditory Research* (Springer, New York, 2005).

63. de Cheveigne, A. in The Oxford Handbook of Auditory Science: Hearing Vol. 3 (ed C.J. Plack) (Oxford University Press, 2010).

64 Bernstein, J. G. W. & Oxenham, A. J. An autocorrelation model with place dependence to account for the effect of harmonic number on fundamental frequency discrimination. Journal of the Acoustical Society of America 117, 3816–3831 (2005).

65 Shackleton, T. M. & Carlyon, R. P. The role of resolved and unresolved harmonics in pitch perception and frequency modulation discrimination. Journal of the Acoustical Society of America 95, 3529–3540 (1994).

66 Moore, B. C. J. & Moore, G. A. Discrimination of the fundamental frequency of complex tones with fixed and shifting spectral envelopes by normally hearing and hearing-impaired subjects. Hearing Research 182, 153–163 (2003).

67 Oxenham, A. J., Bernstein, J. G. & Penagos, H. Correct tonotopic representation is necessary for complex pitch perception. Proceedings of the National Academy of Sciences 101, 1421–1425 (2004).

68 Moore, B. C. J., Glasberg, B. R. & Peters, R. W. Relative dominance of individual partials in determining the pitch of complex tones. Journal of the Acoustical Society of America 77, 1853–1860 (1985).

69 Houtsma, A. J. M. & Smurzynski, J. Pitch identification and discrimination for complex tones with many harmonics. Journal of the Acoustical Society of America 87, 304–310 (1990).

70. Pressnitzer, D. & Patterson, R. D. in Physiological and Psychophysical Bases of Auditory Function (ed D.J. Breebaart) 97-104 (Shaker Publishing, 2001).

71 Norman-Haignere, S. & McDermott, J. H. Distortion products in auditory fMRI research: Measurements and solutions. Neuroimage 129, 401–413 (2016).

72 Moore, B. C. J. Frequency differences limens for short-duration tones. Journal of the Acoustical Society of America 54, 610–619 (1973).

73 de Gardelle, V., Le Corre, F. & Mamassian, P. Confidence as a common currency between vision and audition. PLoS ONE 11, e0147901 (2016).

74 Boundy-Singer, Z. M., Ziemba, C. M. & Goris, R. L. T. Sensory population activity reveals downstream confidence computations in the primate visual system. Proceedings of the National Academy of Sciences 122, e2426441122 (2025).

75 Maniscalco, B. & Lau, H. A signal detection theoretic approach for estimating metacognitive sensitivity from confidence ratings. Consciousness and Cognition 21, 422–430 (2012).

76 Brier, G. W. Verification of forecasts expressed in terms of probability. Monthly Weather Review 78, 1–3 (1950).

77 Fernandez-Vargas, J., Tremmel, C., Valeriani, D., Bhattacharyya, S., Cinel, C., Citi, L. & Poli, R. Subject- and task-independent neural correlates and prediction of decision confidence in perceptual decision making. Journal of Neural Engineering 18, 046055 (2021).

78. Potard, G. & Burnett, I. in Proceedings of the 2003 International Conference on Auditory Display (eds E. Brazil & B.G. Shinn-Cunningham) 25-28 (Boston University Publications Production Department, 2003).

79 Santala, O. & Pulkki, V. Directional perception of distributed sound sources. Journal of the Acoustical Society of America 129, 1522–1530, doi:10.1121/1.3533727 (2011).

80 Frank, M. Source width of frontal phantom sources: Perception, measurement, and modeling. Archives of Acoustics 38, 311–319 (2013).

81 Whitmer, W. M., Seeber, B. U. & Akeroyd, M. A. The perception of apparent auditory source width in hearing-impaired adults. The Journal of the Acoustical Society of America 135, 3548–3559 (2014).

82 Obleser, J. Metacognition in the listening brain. Trends Cogn. Sci. 48, 100–112 (2024).

83 Yamins, D., Hong, H., Cadieu, C. F., Solomon, E. A., Seibert, D. & DiCarlo, J. J. Performance-optimized hierarchical models predict neural responses in higher visual cortex. Proceedings of the National Academy of Sciences 111, 8619–8624, doi:10.1073/pnas.1403112111 (2014).

84 Kubilius, J., Bracci, S. & Op de Beeck, H. P. Deep neural networks as a computational model for human shape sensitivity. PLoS Comp. Biol. 12, e1004896, doi:10.1371/journal.pcbi.1004896 (2016).

85. Banerjee, A., Saddler, M. R., Arenberg, J. G. & McDermott, J. H. A deep learning framework for understanding cochlear implants. bioRxiv, 2025.2007.2016.665227, doi:10.1101/2025.07.16.665227 (2025).

86 Geisler, W. S. Contributions of ideal observer theory to vision research. Vision Research 51, 771–781, doi:10.1016/j.visres.2010.09.027 (2011).

87 Kell, A. J. E. & McDermott, J. H. Deep neural network models of sensory systems: windows onto the role of task constraints. Curr. Opin. Neurobiol. 55, 121–132, doi:10.1016/j.conb.2019.02.003 (2019).

88. Guo, C., Pleiss, G., Sun, Y. & Weinberger, K. Q. in Proceedings of the 34th International Conference on Machine Learning (eds Doina Precup & Yee Whye Teh) 1321-1330 (PMLR, 2017).

89 Tian, K., Mitchell, E., Zhou, A., Sharma, A., Rafailov, R., Yao, H., Finn, C. & Manning, C. D. Just ask for calibration: Strategies for eliciting calibrated confidence scores from language models fine-tuned with human feedback. arXiv, 2305.14975 (2023).

90 Leng, J., Huang, C., Zhu, B. & Huang, J. Taming overconfidence in LLMs: Reward calibration in RLHF. arXiv, 2410.09724 (2024).

91. Xiong, M., Hu, Z., Lu, X., Li, Y., Fu, J., He, J. & Hooi, B. Can LLMs express their uncertainty? An empirical evaluation of confidence elicitation in LLMs. arXiv, 2306.13063 (2024).

92 Platt, J. Probabilistic outputs for support vector machines and comparisons to regularized likelihood methods. Advances in Large Margin Classifiers 10, 61–74 (1999).

93 Zadrozny, B. & Elkan, C. in Proceedings of the Eighth ACM International Conference on Knowledge Discovery and Data Mining (KDD) 694–699 (2002).

94. Kumar, A., Liang, P. S. & Ma, T. in Advances in Neural Information Processing Systems Vol. 32 (eds H. Wallach et al.) (2019).

95. Krishnan, R. & Tickoo, O. Vol. 33 (eds H. Larochelle et al.) 18237--18248 (2020).

96 Damani, M., Puri, I., Slocum, S., Shenfeld, I., Choshen, L., Kim, Y. & Andreas, J. Beyond binary rewards: Training LMs to reason about their uncertainty. arXiv, 2507.16806 (2025).

97 Amini, A., Schwarting, W., Soleimany, A. & Rus, D. in Proceedings of the 34th International Conference on Neural Information Processing Systems 1251 (Curran Associates Inc., 2020).

98. Doersch, C. Tutorial on variational autoencoders. *arXiv*, 1606.05908 (2021).

99 Arbel, J., Pitas, K., Vladimirova, M. & Fortuin, V. A primer on Bayesian neural networks: Review and debates. arXiv, 2309.16314 (2023).

100 Zahorik, P., Brungart, D. S. & Bronkhorst, A. W. Auditory distance perception in humans: A summary of past and present research. Acta Acustica 91, 409–420 (2005).

101 Zhong, X. & Yost, W. A. How many images are in an auditory scene? Journal of the Acoustical Society of America 141, 2882–2892, doi:10.1121/1.4981118 (2017).

102 Kiani, R., Corthell, L. & Shadlen, M. N. Choice certainty is informed by both evidence and decision time. Neuron 84, 1329–1342, doi:10.1016/j.neuron.2014.12.015 (2014).

103 Desender, K., Boldt, A., Verguts, T. & Donner, T. H. Confidence predicts speed-accuracy tradeoff for subsequent decisions. eLife 8, e43499, doi:10.7554/eLife.43499 (2019).

104 Meyer, L. Emotion and Meaning In Music. (University of Chicago Press, 1961).

105 Huron, D. Sweet Anticipation: Music and the Psychology or Expectation. (MIT Press, 2006).

106 Di Liberto, G. M., Pelofi, C., Bianco, R., Patel, P., Mehta, A. D., Herrero, J. L., de Cheveigné, A., Shamma, S. & Mesgarani, N. Cortical encoding of melodic expectations in human temporal cortex. eLife 9, e51784 (2020).

107 Sankaran, N., Leonard, M. K., Theunissen, F. & Chang, E. F. Encoding of melody in the human auditory cortex. Science Advances 10, eadk0010 (2024).

108 Saddler, M. R., Dau, T. & McDermott, J. H. Towards individualized models of hearing-impaired speech perception. The 6th Clarity Workshop on Improving Speech-in-Noise for Hearing Devices, 7-11 (2025).

109 Zylberberg, A., Roelfsema, P. R. & Sigman, M. Variance misperception explains illusions of confidence in simple perceptual decisions. Consciousness and Cognition 27, 246–253 (2014).

110 Maniscalco, B., Peters, M. A. & Lau, H. Heuristic use of perceptual evidence leads to dissociation between performance and metacognitive sensitivity. Attention, Perception, and Psychophysics 78, 923–937 (2016).

111 Peters, M. A. K., Thesen, T., Ko, Y. D., Maniscalco, B., Carlson, C., Davidson, M., Doyle, W., Kuzniecky, R., Devinsky, O., Halgren, E. & Lau, H. Perceptual confidence neglects decision-incongruent evidence in the brain. Nature Human Behaviour 1, 0139, doi:10.1038/s41562-017-0139 (2017).

112. Shinn-Cunningham, B. G., Desloge, J. G. & Kopco, N. in 2001 IEEE Workshop on the Applications of Signal Processing to Audio and Acoustics. (IEEE).

113 Yadav, S. & Foster, M. E. GISE-51: A scalable isolated sound events dataset. 2103.12306 (2021).

114. Gemmeke, J. F., Ellis, D. P. W., Freedman, D., Jansen, A., Lawrence, W., Moore, R. C., Plakal, M. & Ritter, M. in IEEE ICASSP 2017.

115 McDermott, J. H. & Simoncelli, E. P. Sound texture perception via statistics of the auditory periphery: Evidence from sound synthesis. Neuron 71, 926–940, doi:10.1016/j.neuron.2011.06.032 (2011).

116 McWalter, R. I. & McDermott, J. H. Adaptive and selective time-averaging of auditory scenes. Current Biology 28, 1405–1418, doi:10.1016/j.cub.2018.03.049 (2018).

117 Kell, A. J. E. & McDermott, J. H. Invariance to background noise as a signature of non-primary auditory cortex. Nature Communications 10, 3958 (2019).

118 Glasberg, B. R. & Moore, B. C. J. Derivation of auditory filter shapes from notched-noise data. Hearing Research 47, 103–138, doi:10.1016/0378-5955(90)90170-T (1990).

119 Gneiting, T. & Raftery, A. E. Strictly proper scoring rules, prediction, and estimation. Journal of the American statistical Association 102, 359–378 (2007).

120 Blanchard, P., Higham, D. J. & Higham, N. J. Accurately computing the log-sum-exp and softmax functions. IMA Journal of Numerical Analysis 41, 2311–2330 (2021).

121 Abromowitz, M. & Stegun, I. A. Handbook of mathematical functions. (United States Government Printing Office, 1972).

122 Wallach, H., Newman, E. B. & Rosenzweig, M. R. The precedence effect in sound localization. American Journal of Psychology 42, 315–336, doi:10.2307/1418275 (1949).

123 Spearman, C. The proof and measurement of association between two things. The American Journal of Psychology 15, 72–101 (1904).

124 Spearman, C. Correlation calculated from faulty data. British Journal of Psychology 3, 271–295 (1910).

125 Moore, B. C. J., Huss, M., Vickers, D. A., Glasberg, B. R. & Alcántara, J. I. A test for the diagnosis of dead regions in the cochlea. British Journal of Audiology 34, 205–224, doi:10.3109/03005364000000131 (2000).

126 White, L. J. & Plack, C. J. Temporal processing of the pitch of complex tones. Journal of the Acoustical Society of America 103, 2051–2063 (1998).

127 Kawahara, H., Morise, M., Takahashi, T., Nisimura, R., Irino, T. & Banno, H. in IEEE ICASSP 3933-3936 (2008).

128 Köhn, A., Stegen, F. & Baumann, T. in The Tenth International Conference on Language Resources and Evaluation (LREC’16). 4644-4647.

129 Engel, J., Resnick, C., Roberts, A., Dieleman, S., Norouzi, M., Eck, D. & Simonyan, K. in Proceedings of the 34th International Conference on Machine Learning-Volume 70. 1068-1077 (JMLR.org).

130 Woods, K. J. P., Siegel, M. H., Traer, J. & McDermott, J. H. Headphone screening to facilitate web-based auditory experiments. Attention, Perception, and Psychophysics 79, 2064–2072 (2017).

131 Woods, K. J. P. & McDermott, J. Schema learning for the cocktail party problem. Proceedings of the National Academy of Sciences 115, E3313–E3322 (2018).

132 McWalter, R. & McDermott, J. H. Illusory sound texture reveals multi-second statistical completion in auditory scene analysis. Nature Communications 10, 5096 (2019).

133 McPherson, M. J. & McDermott, J. H. Time-dependent discrimination advantages for harmonic sounds suggest efficient coding for memory. Proceedings of the National Academy of Sciences 117, 32169–32180, doi:10.1101/2020.05.07.082511 (2020).

134 McPherson, M. J., Dolan, S. E., Durango, A., Ossandon, T., Valdés, J., Undurraga, E. A., Jacoby, N., Godoy, R. A. & McDermott, J. H. Perceptual fusion of musical notes by native Amazonians suggests universal representations of musical intervals. Nature Communications 11, 2786 (2020).

135 Traer, J., Norman-Haignere, S. V. & McDermott, J. H. Causal inference in environmental sound recognition. Cognition 214, 104627 (2021).

136 McPherson, M. J., Grace, R. C. & McDermott, J. H. Harmonicity aids hearing in noise. Attention, Perception, and Psychophysics 84, 1016–1042 (2022).

137 McPherson, M. J. & McDermott, J. H. Diversity in pitch perception revealed by task dependence. Nature Human Behavior 2, 52–66, doi:10.1038/s41562-017-0261-8 (2018).

138. metadpy toolbox, <https://github.com/embodied-computation-group/metadPy> (

